# Phase-Separated RNA Condensates Govern Cas13 Target Accessibility and Cleavage

**DOI:** 10.64898/2026.05.12.724747

**Authors:** Gurjeet Kaur Gill Jagjeet Singh, Wenxin Hu, Carolyn Shembrey, Adrian Hodel, Ilia Voskoboinik, Paul McMillan, Joseph Trapani, Mohamed Fareh

## Abstract

RNA-guided CRISPR-Cas13 nucleases must efficiently locate and target specific transcripts amidst the millions of other RNA molecules that are spatially regulated in the cell. Yet the mechanisms by which Cas13 finds its targets within this crowded, compartmentalized environment remain elusive. Here, we show that diverse Cas13 orthologs assemble into distinct cytoplasmic granules in a crRNA-concentration dependent manner. These granules exhibit hallmark features of phase-separated condensates, displaying liquid-like viscoelasticity and dynamic molecular exchange with the cytoplasm, as demonstrated by FRAP analysis in both human and bacterial cells. Molecular profiling revealed that Cas13 co-localizes with polyadenylated RNAs and noncoding RNAs within condensates that display markers of canonical RNA granules. Biochemical purification coupled with RNA sequencing shows Cas13 associated with thousands of transcripts within condensates, likely mediated by electrostatic interactions with its positively charged surface. Notably, Cas13 retains catalytic activity within these condensates, efficiently cleaving co-localized targets, whereas RNA species excluded from the condensates remain largely protected from Cas13 cleavage. This data indicates that condensate-based spatial organization facilitates efficient sampling and binding of diverse RNA targets by concentrating Cas13 and its substrates within a confined, liquid-like compartment. Together, our findings uncover a conserved spatial mechanism regulating Cas13 activity across bacterial and mammalian cells, where localization within RNA dense granules governs Cas13 activity in cells.

## Introduction

RNA-guided CRISPR-Cas13 nucleases have emerged as powerful tools for programmable transcriptome manipulation, enabling applications from functional genomic screens to therapeutic silencing of oncogenes and pathogenic viral RNAs (Abudayyeh et al., 2017; Abudayyeh et al., 2016; East-Seletsky et al., 2017; East-Seletsky et al., 2016; Hu et al., 2025; McCoullough et al., 2024; Shembrey et al., 2024; Shmakov et al., 2015). Guided by a CRISPR RNA (crRNA) containing a programmable spacer sequence, Cas13 recognises target transcripts through RNA-RNA base pairing, thus catalysing their degradation (O’Connell, 2019; Zhang et al., 2019). Yet for Cas13 to silence RNA in living cells, it must first locate its target transcripts amongst tens of thousands of non-target RNAs in a crowded and highly structured intracellular environment (Abudayyeh et al., 2017; Fareh et al., 2021; Shu et al., 2024). How Cas13 efficiently locates its target transcripts in this complex cellular landscape remains largely unknown.

Rather than being uniformly distributed, cellular RNAs exhibit defined spatial organisation that regulates their processing, translation, and decay, enabling precise control of gene expression (Buxbaum et al., 2015; Czaplinski & Singer, 2006; Ren et al., 2024; Ryder & Lerit, 2018). Many transcripts localize to specific subcellular compartments through directed transport, local anchoring, or partitioning into membraneless biomolecular condensates, which are dynamic, RNA-protein assemblies that form via multivalent interactions. These condensates, including Processing bodies (P-bodies) and Stress Granules (SGs), create a structured landscape where RNAs and RNA-binding proteins are selectively concentrated or excluded, shaping which molecules encounter one another (André & Spruijt, 2020; Banani et al., 2017; Biayna & Dumbović, 2025; Hofmann et al., 2021; Kedersha et al., 2005; Pessina et al., 2025; Stroberg & Schnell, 2018; Strulson et al., 2012; Zhang et al., 2021). Such spatial organization can profoundly influence RNA-protein interactions: compartmentalization may accelerate target recognition by concentrating binding partners within confined volumes, or conversely, restrict access by sequestering substrates away from their cognate enzymes. Phase-separated RISC-microRNA condensates, for instance, enhance silencing efficiency by orders of magnitude through co-concentrating Argonaute complexes and target mRNAs (Liu et al., 2005; Sheu-Gruttadauria & MacRae, 2018).

Mechanistic studies of Cas13 have relied predominantly on *in vitro* biochemical assays and bulk cellular measurements that implicitly assume a well-mixed cellular environment, where target accessibility depends solely on diffusion and sequence complementarity (Abudayyeh et al., 2017; East-Seletsky et al., 2016; Gootenberg et al., 2017; Hu et al., 2024; Kellner et al., 2019; Konermann et al., 2018). These models however cannot account for the spatially compartmentalized transcriptome of living cells. When Cas13 must physically encounter target RNAs for recognition and cleavage, subcellular organization could profoundly shape accessibility. RNAs sequestered away from Cas13 may evade targeting despite perfect sequence complementarity, while colocalizing transcripts should undergo preferential cleavage. Whether and how intracellular RNA organization shapes Cas13 localization, target search, and silencing efficiency remains uncharacterized.

Here, using live-cell imaging we show that *Psp*Cas13b localises within RNA condensates that share markers of cytoplasmic granules, whose formation is dependent on intracellular crRNA abundance. These condensates are highly enriched with cellular RNA, offering enhanced colocalization and interaction between Cas13 and its target RNA. Biochemical purification and pull-down assays demonstrate that Cas13 is catalytically active within these condensates, leading to efficient target cleavage of both exogenous and endogenous transcripts. RNA sequencing further revealed that Cas13 associates with a distinct subset of coding and non-coding RNAs that we termed the *Psp*Cas13b-RNA interactome. Notably, RNA molecules that colocalise with Cas13 within condensates exhibit efficient cleavage, whereas those excluded, show significantly lower cleavage efficiency, further suggesting that spatial colocalization within RNA condensates enhances target search and cleavage.

Together, these findings support a model in which Cas13 target search is governed not only by crRNA-target complementarity, but by spatial partitioning within the cell. Intracellular RNA organisation represents a previously unrecognised and likely general regulator of Cas13 nuclease activity, with broad implications of how spatial localisation shapes productive target encounters and cleavage efficiency across biological systems.

## Results

### d*Psp*Cas13b localises within cytoplasmic granules

We sought to understand how Cas13 locates its target RNA in the crowded and highly compartmentalized cytoplasm of human cells. We first established the baseline subcellular localisation of *Psp*Cas13b using confocal microscopy. HEK293T cells were co-transfected with plasmids encoding a fluorescently tagged catalytically inactive Cas13b (d*Psp*Cas13b-2xmNeonGreen) either alone or together with its cognate non-targeting crRNA (NTcrRNA) **(Figure 1A-B)**.

**Figure 1.**
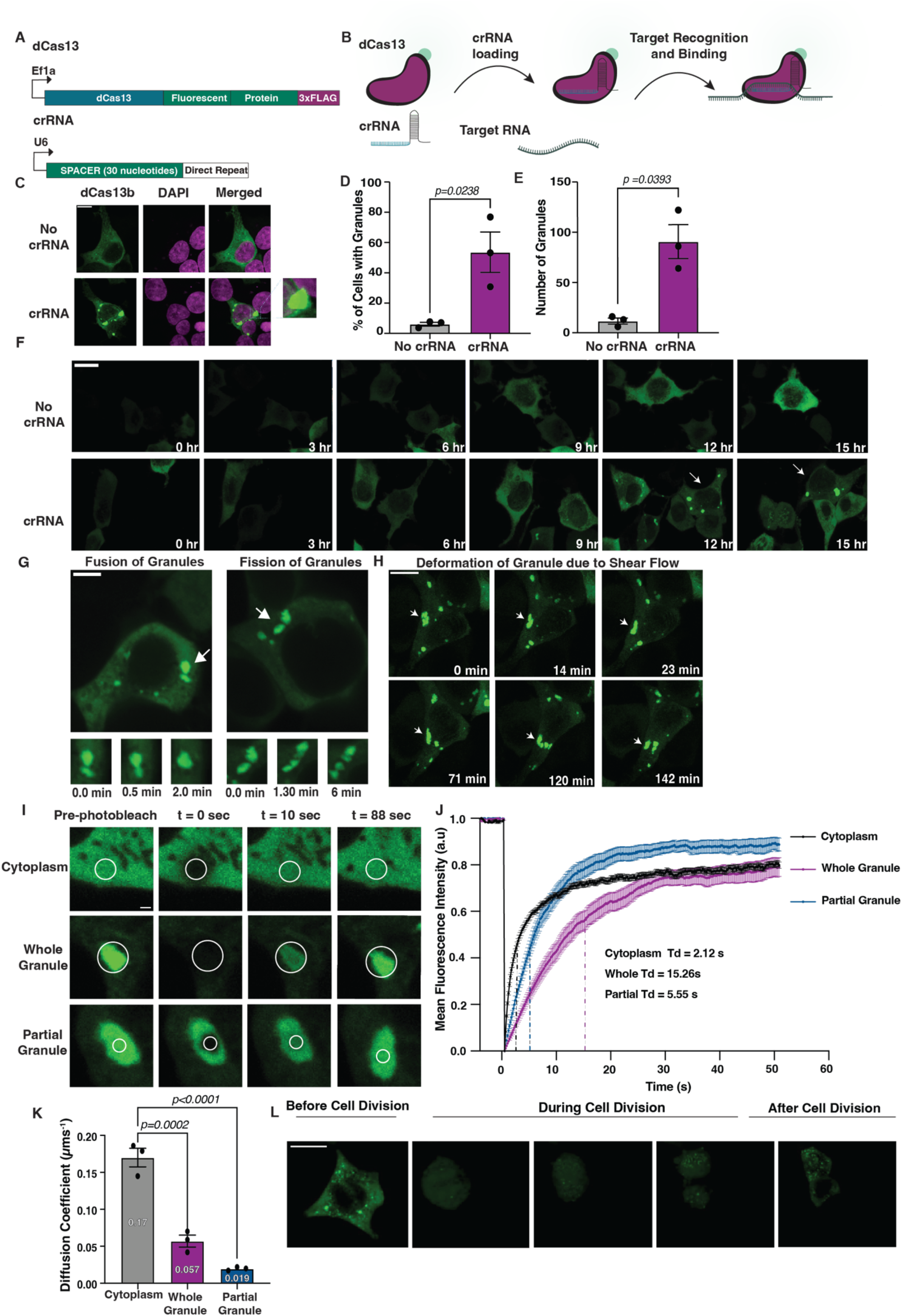
dpspCas13b present as cytoplasmic foci which emulate properties observed in liquid-like particles. **(A)** Schematic illustration of the fluorescently labelled catalytically inactive dpspCas13b fused to 2xmNeonGreen fluorescent proteins (top) and the crRNA (bottom) construct. **(B)** Illustration of the assay used for target RNA search visualisation. **(C)** Confocal microscopy images taken using 100x objective lens of HEK293T cells transfected with dpspCas13b-2xmNeonGreen alone or with its corresponding crRNA counterstained with DAPI. Scale bar = 20μm. **(D-E)** Quantification of percentage of cells with granules and total number of granules per field of view (n=3). **(F)** Representative snapshots of live-cell confocal microscopy of HEK293T cells expressing dpspCas13b-2xmNeonGreen. Imaging was carried out 10 hours post-transfection. The upper panel are cells without crRNA (apo-dpspCas13b), and the lower panel are cells expressing both dpspCas13b and its corresponding crRNA depicting 3-hour intervals snapshots. Scale bar = 10 μm. **(G)** Single-cell snapshots of cells exhibiting granule formation. The panel on the left depicts a granule labelled with a white arrow undergoing fusion events. The panel on the right depicts multiple granules undergoing fission events. Scale bar = 5 μm. **(H)** Snapshots of a single granule undergoing shear force deformation while moving between the nuclear envelope and plasms membrane. Scale bar = 10 μm. **(I)** Representative snapshots of fluorescent recovery after photobleaching with regions of interest highlighted with a white circular bleach radius between 0.5 μm – 0.9 μm. The upper panel depicts cytoplasmic dpspCas13b proteins being photobleached. The middle panel depicts whole granule photobleaching and the bottom panel depicts partial granule photobleaching. Snapshots are taken pre-photobleaching, upon photobleaching (t = 0 seconds) and after recovery (t = 50 seconds). Scale bar = 2 μm. **(J)** Mean fluorescent recovery curves (black = cytoplasm, magenta = whole granule, blue = partial granule (n=30). Halftime of diffusion (Td) is represented with black vertical dashed lines. **(K)** Diffusion coefficients represented as bar graphs with mean diffusion coefficient within each corresponding bar. **(L)** Snapshots of live cell imaging of cells expressing dpspCas13b-2xmNeonGreen and crRNA before, during and after cell division.

When expressed in the absence of its crRNA, d*Psp*Cas13b displayed a diffuse cytoplasmic distribution. Unexpectedly, co-expression with its cognate crRNA induced the formation of discrete cytoplasmic granules containing d*Psp*Cas13b in ∼52.6% of cells **(Figure 1C-E)**. Under these conditions, d*Psp*Cas13b redistributed from a largely diffusive cytoplasmic pool to enrichment within punctate cytoplasmic assemblies. Because a non-targeting crRNA was used, this localisation is unlikely to reflect specific target binding, suggesting instead that crRNA expression itself alters the intracellular distribution of Cas13. Next, we performed live cell imaging 10 hours post-transfection to capture the formation and dynamic movement of these cytoplasmic granules in real time. When co-expressed with its cognate crRNA, d*Psp*Cas31b progressively re-localized from a diffusive cytoplasmic distribution to distinct punctate granules, first detectable ∼12-hours post transfection, and increasing in number and intensity over time **(Figure 1F)**.

Because fluorescent fusion tags can influence condensation behaviour, we next examined the localisation of catalytically active *Psp*Cas13b lacking the fluorescent tag by immunofluorescence (Barkley et al., 2024; Pandey et al., 2024). Under identical crRNA co-expression conditions, *Psp*Cas13b exhibited a similar cytoplasmic granular distribution **(Figure S1)**, indicating that granule localisation is not driven by the fluorescent fusion tag. Collectively, these data establish that *Psp*Cas13b can redistribute from a diffuse cytoplasmic distribution to discrete cytoplasmic granules under crRNA expression conditions.

### dPspCas13b assemblies exhibit liquid-like behaviour

We next asked whether these cytoplasmic assemblies exhibit properties characteristic of biomolecular condensates. Live-cell imaging revealed dynamic morphological behaviours including granule fusion, fission, and deformation during movement within the cytoplasm **(Figure 1G-H)**, consistent with behaviours previously reported for liquid-like condensates (Fare et al., 2021; Wang et al., 2022). Liquid-like particles do not have an encapsulating membrane to limit molecular exchange with their environment and thus are able to rapidly exchange biomolecules with their surrounding cytoplasm (Brangwynne et al., 2009). Additionally, they exhibit higher protein density and weaker molecular motion than the surrounding cytoplasm as they have a higher viscosity (Brangwynne et al., 2009; Taylor et al., 2019). To further assess the biophysical properties of these assemblies, we performed fluorescence recovery after photobleaching (FRAP). Photobleaching of cytoplasmic d*Psp*Cas13b resulted in rapid fluorescence recovery, consistent with fast diffusion in the cytosol. In contrast, photobleaching of entire granules produced slower recovery kinetics, indicating slower exchange between the granule and surrounding cytoplasm. Partial photobleaching within granules recovered more rapidly, suggesting internal molecular mixing **(Figure 1I-J)**.

Quantitative analysis revealed that diffusion of d*Psp*Cas13b within granules was markedly reduced compared with the cytoplasm, with diffusion coefficients of 0.17 µm^2^ s^−1^, 0.057 µm^2^ s^−1^ and 0.019 µm^2^ s^−1^ for cytoplasmic, whole-granule and partial-granule measurements, respectively **(Figure 1K)**.

The slower recovery of whole-granule photobleaching relative to partial-granule photobleaching indicates rapid internal mixing within the condensate but slower exchange across the condensate-cytoplasm interface. In addition to their high viscosity, a defining characteristic of cytoplasmic condensates is their complete dissolution during mitosis and reformation in daughter cells (Draetta & Beach, 1988; Rai et al., 2018; Valverde et al., 2023). Consistent with this behaviour, d*Psp*Cas13b granules dissolved during mitosis and reappeared after cell division **(Figure 1L)**. Collectively, these observations demonstrate that d*Psp*Cas13b accumulates in dynamic cytoplasmic assemblies that exhibit hallmark properties of liquid-like biomolecular condensates.

### dPspCas13b condensates are enriched in RNA and exhibit protein markers of RNA condensates

Having established that d*Psp*Cas13b localises within liquid-like cytoplasmic condensates, we next asked whether these assemblies contain cellular RNA. To address this, we performed RNA fluorescence in situ hybridisation (FISH) using a poly(dT) probe to detect poly(A) RNA. Polyadenylated transcripts were strongly enriched within d*Psp*Cas13b condensates relative to the surrounding cytoplasm **(Figure 2A-B)**, indicating that these structures represent RNA-rich compartments. To determine whether this enrichment extends to additional RNA species, we examined the localisation of ribosomal RNA and a representative cellular transcript. RNAFISH analysis revealed enrichment of 18S rRNA within dPspCas13b condensates compared to the surrounding cytoplasm **(Figure 2C-D)**. Similarly, β-actin (ACTB) mRNA was observed within these condensates **(Figure 2E)**, indicating that multiple RNA species accumulate in these assemblies.

**Figure 2.**
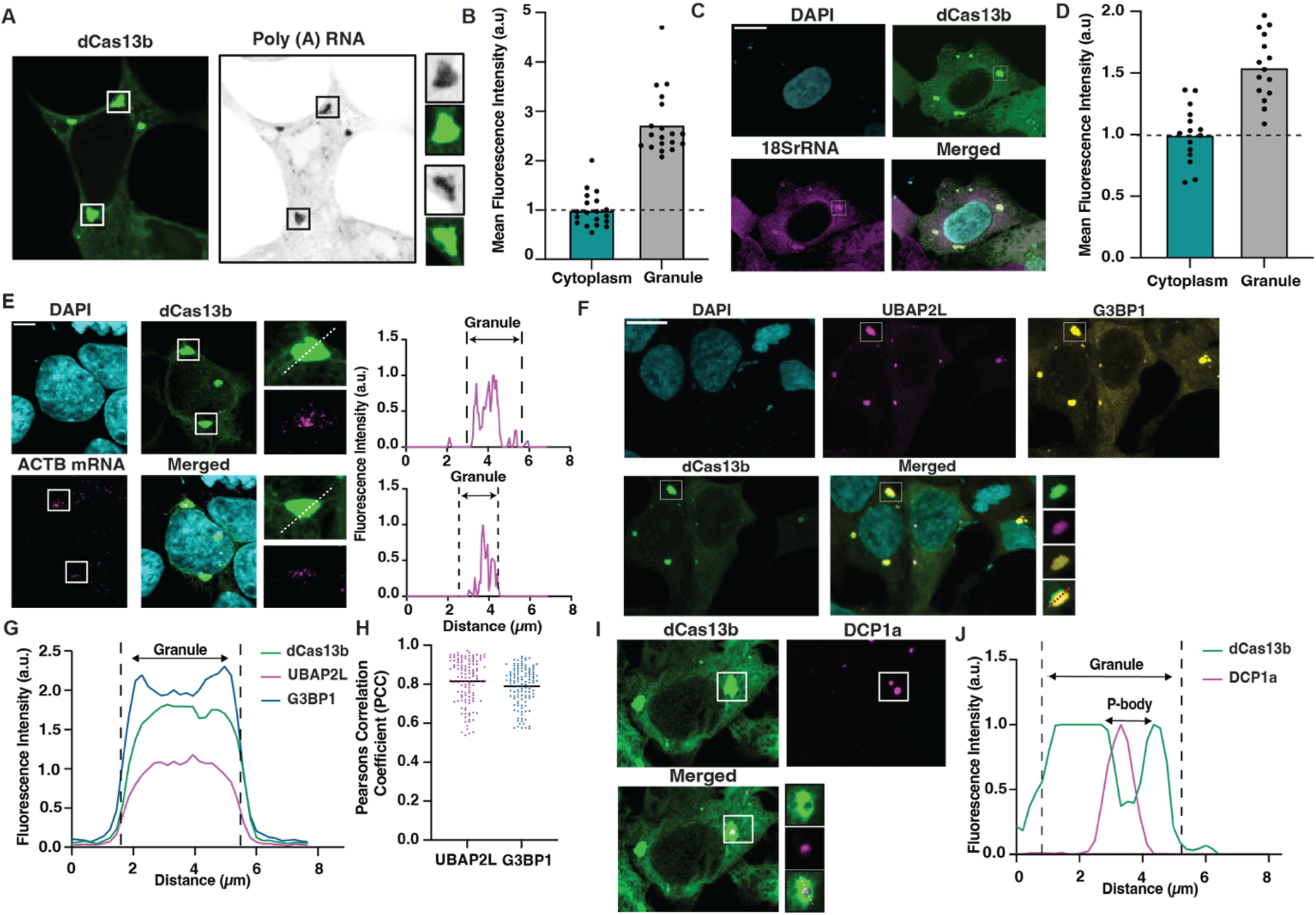
dpspCas13b condensates are enriched in RNA and exhibit markers associated with RNA condensates. **(A)** Representative confocal images of HEK293T cells expressing dPspCas13b-2xmNeonGreen stained for poly(A) RNA using a poly(dT) probe. White boxes indicate condensates shown at higher magnification in adjacent panels. **(B)** Quantification of mean fluorescence intensity of poly(A) RNA in the cytoplasm versus within dPspCas13b condensates (n = 19). **(C)** Representative RNAFISH images showing localisation of 18S rRNA relative to dPspCas13b condensates. **(D)** Quantification of mean fluorescence intensity of 18S rRNA in the cytoplasm versus within condensates (n = 15). **(E)** RNAFISH analysis showing localisation of ACTB mRNA relative to dPspCas13b condensates. Insets highlight individual condensates, and line profiles show fluorescence intensity across the indicated region of interest. Scale bar = 5 µm. **(F)** Representative immunofluorescence images of HEK293T cells expressing dPspCas13b and stained for the stress-granule–associated RNA-binding proteins UBAP2L and G3BP1. White boxes indicate condensates shown at higher magnification in adjacent panels. **(G)** Fluorescence intensity profiles across a condensate showing enrichment of dPspCas13b, UBAP2L and G3BP1. **(H)** Pearson’s correlation coefficient (PCC) quantification of colocalisation between dPspCas13b and UBAP2L or G3BP1 (n = 123). **(I)** Representative confocal images showing spatial association between dPspCas13b condensates and the P-body marker DCP1a. Insets show magnified views of condensates and adjacent P-bodies. **(J)** Fluorescence intensity profile across the region of interest illustrating spatial juxtaposition of dPspCas13b condensates and DCP1a-positive P-bodies without overlap. Scale bars: 10 μm unless otherwise indicated.

Because RNA-rich cytoplasmic condensates frequently correspond to stress granules (SGs) or processing bodies (P-bodies), we next examined whether d*Psp*Cas13b condensates share molecular markers of these structures. Immunofluorescence staining revealed strong colocalization of d*Psp*Cas13b with the SG-associated RNA-binding proteins UBAP2L and G3BP1 **(Figure 2F-H)**, indicating that these assemblies share markers associated with stress granules despite forming in the absence of exogenous stress. In contrast, staining for the P-body marker DCP1a revealed spatial juxtaposition rather than direct colocalization. P-bodies frequently appeared docked adjacent to d*Psp*Cas13b condensates without evidence of fusion **(Figure 2I-J)**, suggesting that these structures remain distinct but interact with neighbouring RNA condensates.

The co-enrichment of G3BP1 and UBAP2L raised the question of whether d*Psp*Cas13b condensate formation mechanistically was dependent on these canonical stress granule nucleators. G3BP1 and G3BP2 are well-established drivers of stress granule assembly, and their loss has been shown to abolish canonical stress granule formation (Nancy Kedersha et al., 2016; McEwen et al., 2005; Yang et al., 2020). To test whether these condensates reflect canonical stress granules, we generated G3BP1/2 double knockout HEK293T cell lines confirmed by immunoblotting and sequencing analyses **(Supplementary Figure S2)**. Strikingly, d*Psp*Cas13b condensates remained detectable in G3BP1/2 deficient cells, though their frequency was reduced compared to the wild-type control **(Supplementary Figure S3)**. This partial dependence indicates that while G3BP1/2 contribute to condensate formation, they are not strictly required. Together, these results demonstrate that d*Psp*Cas13b condensates are RNA-rich cytoplasmic assemblies that share molecular features with stress granules yet form through a partially non-canonical mechanism. Their spatial distinction from P-bodies, combined with their G3BP1/2-independent formation, support a model in which crRNA itself nucleates a distinct class of RNA condensate into which d*Psp*Cas13b is subsequently recruited through a mechanism that engages but does not depend upon canonical stress granule machinery.

### crRNA expression is sufficient to induce formation of cytoplasmic condensates

Having characterised the physiochemical properties of these d*Psp*Cas13b condensates, we next questioned whether crRNA loading was the determining factor mediating formation of these condensates as formation was only observed upon co-expression of both d*Psp*Cas13b and its cognate crRNA. Expression of d*Psp*Cas13b alone was unable to lead to formation of condensates that co-express UBAP2L **(Figure 3A)**.

**Figure 3.**
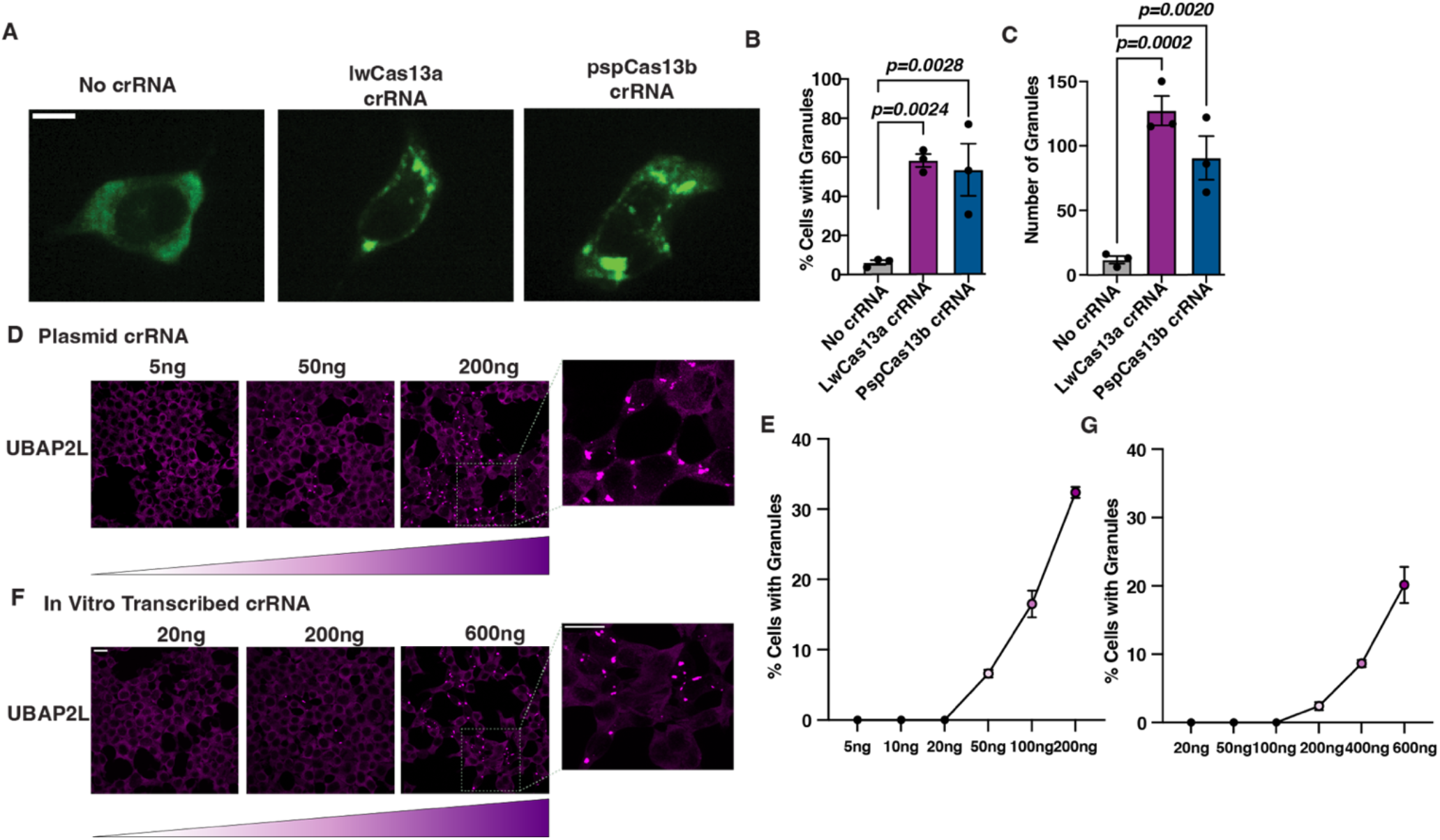
Intracellular RNA abundance promotes formation of cytoplasmic condensates. **(A)** Representative confocal images of HEK293T cells expressing dPspCas13b-2xmNeonGreen in the absence of crRNA or together with crRNAs derived from the LwCas13a or PspCas13b systems. Condensates are observed in the presence of crRNA but not in the absence of crRNA. Scale bar = 10 μm. **(B-C)** Quantification of the percentage of cells containing condensates (B) and the number of condensates per field of view. **(C)** under the indicated conditions. Data represent mean ± SEM from independent experiments (n = 3). **(D)** Confocal images of HEK293T cells transfected with increasing amounts of plasmid-encoded non-targeting crRNA (NTcrRNA). UBAP2L immunostaining was used as a marker for condensate formation. Insets show magnified regions containing condensates. Scale bar = 20 μm. **(E)** Dose-dependent quantification of the percentage of cells containing condensates following plasmid-encoded NTcrRNA delivery. Data represent mean ± SEM (n = 3). **(F)** Confocal images of HEK293T cells transfected with increasing amounts of in vitro-transcribed (IVT) NTcrRNA. UBAP2L staining was used to visualise cytoplasmic condensates. Insets show representative magnified regions of condensates. Scale bar = 20 μm. **(G)** Dose-dependent quantification of the percentage of cells containing condensates following IVT NTcrRNA delivery. Data represent mean ± SEM (n = 3).

We ectopically expressed d*Psp*Cas13b together either with its cognate crRNA or a paralogue crRNA that belongs to the *Lw*Cas13a family. This crRNA possesses different RNA features compared to the crRNA from the *Psp*Cas13b family, such as a different direct repeat sequence and a flipped orientation of the crRNA which prohibits any crRNA loading into d*Psp*Cas13b (Slaymaker et al., 2019). Therefore, this experimental setting allows for decoupling of crRNA expression from crRNA loading, thus allowing us to assess which of these processes are contributing to condensate formation. We observed significant condensate formation in all conditions except for the No crRNA condition **(Figure 3A - C)** suggesting that it is not the loading of the cognate crRNA that is the driving force mediating formation of these condensates, but the intracellular concentration of the crRNA. Previous studies have shown that elevated RNA concentrations can promote cytoplasmic condensation through multivalent RNA-RNA and RNA-protein interactions (Aumiller et al., 2016; Bounedjah et al., 2014; Hofmann et al., 2021; Van Treeck et al., 2018). Consistent with this, increasing intracellular crRNA abundance resulted in a dose-dependent increase in UBAP2L-positive condensates following plasmid-based crRNA expression **(Figure 3D-E)**.

To determine whether crRNA abundance alone is sufficient to drive condensate formation, we delivered in vitro-transcribed (IVT) crRNA directly into cells. Increasing amounts of IVT crRNA similarly induced formation of UBAP2L-positive cytoplasmic condensates in a dose-dependent manner **(Figure 3F-G)**. Together, these results indicate that elevated intracellular RNA abundance is sufficient to promote formation of cytoplasmic condensates.

### Condensate partitioning is conserved across Cas13 orthologues

Cas13 effectors comprise of a diverse family of single-effector, RNA-guided RNases that are highly divergent at the sequence level outside of their HEPN catalytic domains, implying substantial evolutionary distance across subfamilies (Makarova et al., 2025; Smargon et al., 2017; Yan et al., 2018). Despite this sequence divergence, Cas13 proteins share a broadly conserved organisation consisting of an RNA recognition lobe and a nuclease lobe containing two HEPN domains that form the catalytic centre (Liu et al., 2017; Slaymaker et al., 2019).

To determine whether localisation to RNA-rich cytoplasmic condensates is conserved across the Cas13 family, we examined orthologues spanning the three major subtypes Cas13a, Cas13b and Cas13d **(Figure 4A-C)**.

**Figure 4.**
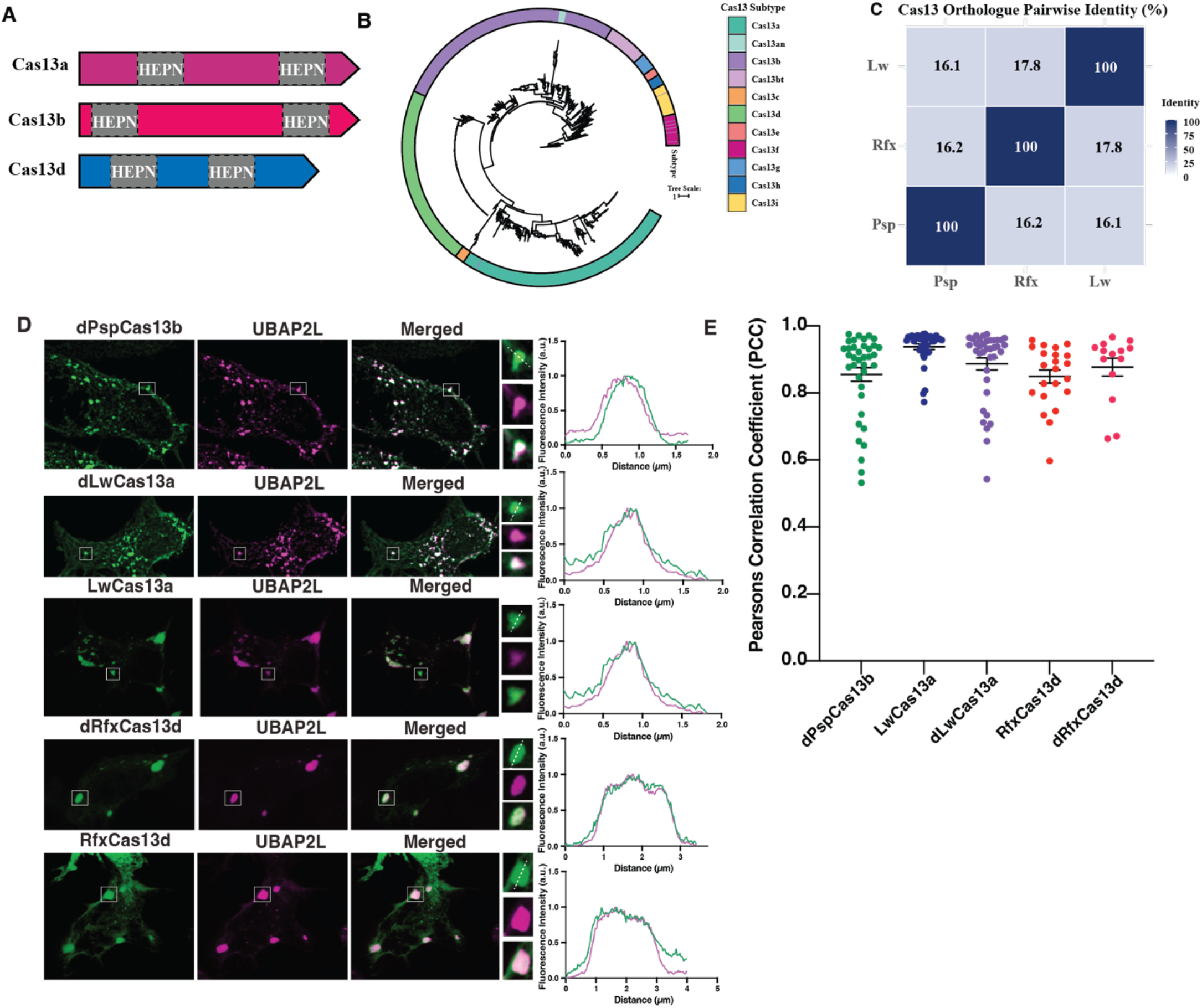
Condensate localisation is conserved across evolutionarily distinct Cas13 orthologues. **(A)** Schematic domain organization of Cas13a, Cas13b, and Cas13d proteins highlighting the conserved dual HEPN RNase domains (grey; schematic not to scale). **(B)** Maximum-likelihood phylogeny of Cas13 proteins constructed from a multiple sequence alignment (MAFFT), alignment trimming (trimAl), and tree inference using IQ-TREE2 with ModelFinder and branch support estimation (ultrafast bootstrap and SH-aLRT). Branch lengths represent substitutions per site (evolutionary scale bar shown = 0.7). Tip annotations are displayed as a circular colour ring indicating subtype categories. Sequence accession numbers and subtype calls were curated from published Cas13 phylogenies and datasets, including the accession list reported for Cas13 phylogeny construction by Adler et al., metagenomic subtype definitions for Cas13Bt-A/B and Cas13e–i by Hu et al., the original Cas13X/Y description by Xu et al., and additional subtype sampling including Cas13an from recent structure-guided discovery (Adler et al., 2022; Hu et al., 2022; C. Xu et al., 2021). **(C)** Pairwise amino-acid identity (%) between the three Cas13 orthologues selected for experimental characterisation (LwCas13a, PspCas13b, RfxCas13d), calculated using EMBOSS needle global alignments. **(D)** Confocal microscopy images depicting single cell representatives of different Cas13 orthologs. Green = Cas13 (“d” nomenclature indicates a catalytically inactive effector) Magenta = UBAP2L. White box indicates a granule, with panels showing magnified view. Dotted line indicates region of interest used for line intensity analysis plots. **(E)** Colocalization analysis performed using Pearson’s Correlation Coefficient (PCC).

We used *Lw*Cas13a (*Leptotrichia wadei*), *Psp*Cas13b (*Prevotella sp*.) and *Rfx*Cas13d (*Ruminococcus flavefaciens*; CasRx), which are widely used platforms for programmable RNA targeting (Abudayyeh et al., 2017; Gootenberg et al., 2017; Konermann et al., 2018). Pairwise sequence comparisons confirmed that these enzymes are highly divergent, sharing only ∼16–18% amino acid identity between subtypes **(Figure 4C)**.

To assess whether condensate partitioning represents a conserved property of Cas13 enzymes, we examined the localisation of both catalytically active and inactive variants in mammalian cells. Across all orthologues tested, Cas13 proteins were enriched within UBAP2L-positive cytoplasmic condensates **(Figure 4D)**. Quantification of colocalization using Pearson’s correlation coefficient revealed strong overlap between Cas13 and UBAP2L signals across orthologues, including *Lw*Cas13a (∼0.94), *dLw*Cas13a (∼0.89), *Rfx*Cas13d (∼0.85), d*Rfx*Cas13d (∼0.88) and d*Psp*Cas13b (∼0.81) **(Figure 4E)**. Together, these observations demonstrate that partitioning of Cas13 into RNA-rich cytoplasmic condensates is conserved across evolutionarily divergent Cas13 subtypes and occurs independently of HEPN-mediated catalytic activity.

### PspCas13b forms discrete intracellular assemblies in bacteria and localises to rRNA-rich regions

Having established that *Psp*Cas13b, *Lw*Cas13a, and *Rfx*Cas13d preferentially partition into RNA-rich condensates in mammalian cells, we next asked whether this spatial behaviour extends to a bacterial context. Cas13 effectors evolved as RNA-guided RNases within Class 2 Type VI CRISPR-Cas systems in bacteria, however, it remains unclear whether the partitioning observed in mammalian cells reflects an intrinsic property of Cas13 or emerges from features of the mammalian cytoplasmic environment. We therefore examined Cas13 localization in bacteria to determine whether it similarly forms discrete intracellular assemblies and associates with RNA-enriched regions. To investigate this, we examined the intracellular localisation of *Psp*Cas13b in *Escherichia coli* BL21(DE3) cells. *Psp*Cas13b was expressed as an EGFP fusion under an IPTG-inducible promoter and imaged by fluorescence microscopy following induction **(Figure 5A)**. In contrast to EGFP alone, which remained diffusely distributed throughout the cytoplasm, *PspCas13b* frequently formed discrete intracellular assemblies exhibiting heterogenous localisation patterns, including polar foci, membrane-associated patches, and multipunctate structures **(Figure 5B-C)**. Because Cas13 is an RNA-guided effector and bacterial RNA exhibits defined spatial organisation, we next asked whether these assemblies reflect RNA-rich regions. RNA fluorescence in situ hybridisation (RNA-FISH) targeting 16S rRNA revealed frequent spatial overlap between *Psp*Cas13b-EGFP assemblies and 16S rRNA signal (**Figure 5D-G**). Quantitative analysis demonstrated strong intensity correlation between *Psp*Cas13b and 16S rRNA (Pearson’s r ≈ 0.7) together with high fractional overlap of *Psp*Cas13b signal with rRNA (Manders’ M ≈ 0.8), whereas a smaller fraction of total rRNA signal overlapped Cas13b assemblies (Manders’ M ≈ 0.4) **(Figure 5H-I)**. By contrast, EGFP alone exhibited substantially lower overlap with 16S rRNA under matched conditions **(Figure 5D-F)**. To determine whether these assemblies represent static aggregates or dynamic structures, we performed fluorescence recovery after photobleaching (FRAP) combined with fluorescence loss in photobleaching (FLIP) to assess molecular exchange between intracellular foci. Selective photobleaching of a polar *Psp*Cas13b focus resulted in partial fluorescence recovery with a mean mobile fraction of ∼39.8% and a recovery half-time of ∼30.7s **(Figure 5J-K)**. Concomitantly, fluorescence decreased at the adjacent unbleached pole while total cellular fluorescence remained constant, indicating redistribution of *Psp*Cas13b molecules between spatially distinct foci.

**Figure 5.**
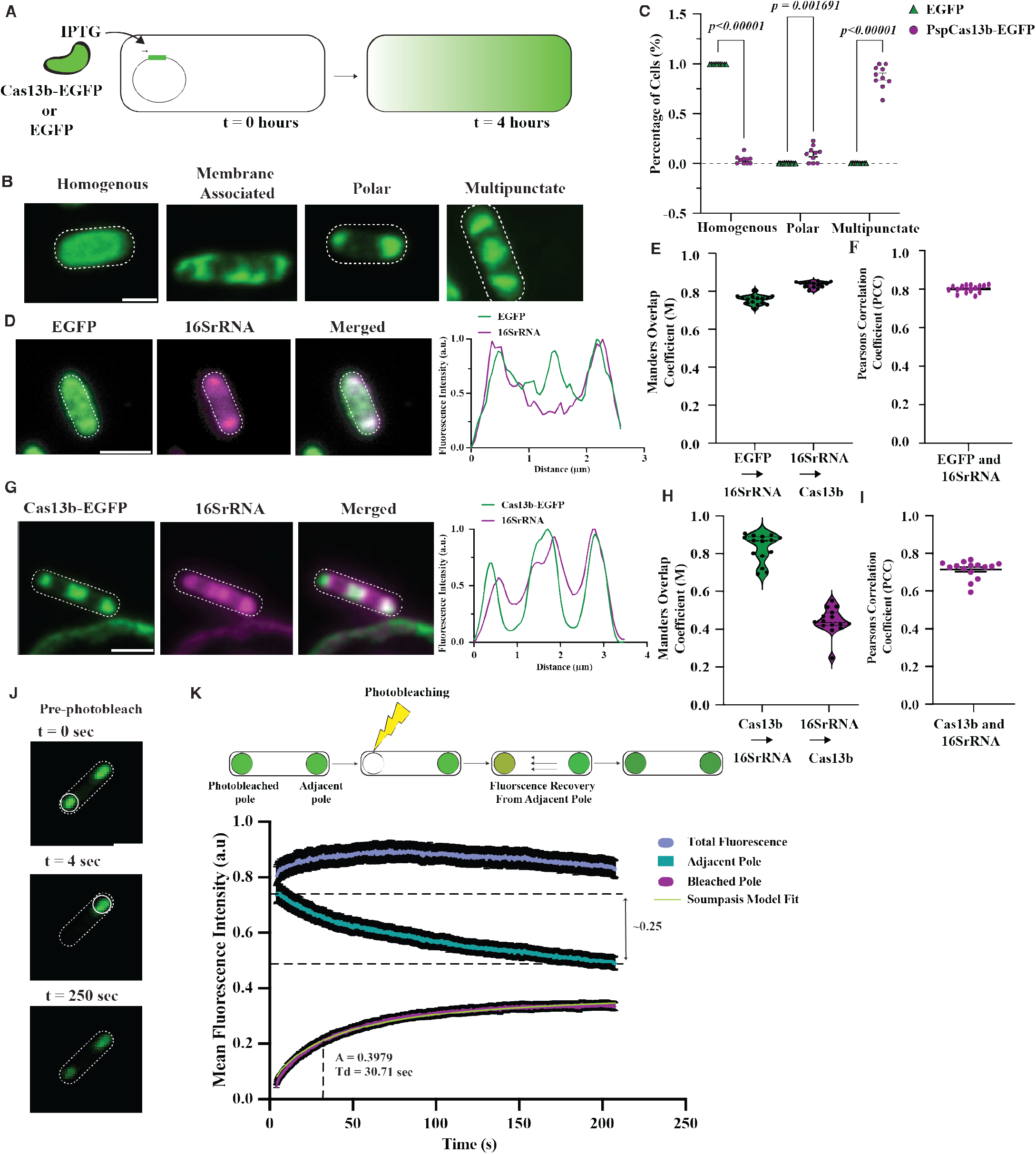
*Psp*Cas13b forms dynamic intracellular assemblies in bacteria that are enriched with 16S rRNA. **(A)** Schematic of the IPTG-inducible expression system used to express PspCas13b-EGFP or EGFP in *E. coli* BL21(DE3). Cells were induced with 0.05 mM IPTG and imaged after 4 h. **(B)** Representative fluorescence images showing distinct localisation patterns observed in cells expressing PspCas13b-EGFP or EGFP, categorised as homogeneous, membrane-associated, polar, or multipunctate. Dashed outlines indicate cell boundaries. Scale bar = 1 μm. **(C)** Quantification of localisation phenotypes in EGFP- and PspCas13b-EGFP–expressing cells. Points represent independent biological replicates and bars indicate mean ± SEM. Approximately n = 300 cells per condition. Statistical comparisons were performed using multiple unpaired t-tests on biological replicate means (n = 3). (**D)** RNA fluorescence in situ hybridisation (RNA-FISH) targeting 16S rRNA (magenta) in cells expressing EGFP (green). Line-scan intensity profiles across the indicated cells show minimal spatial correlation between EGFP and rRNA signals. Scale bar = 1 μm. **(E–F)** Quantification of spatial overlap between EGFP and 16S rRNA using Manders’ overlap coefficient (M) (E) and Pearson’s correlation coefficient (PCC) (F). Each point represents the mean coefficient calculated per field of view. Horizontal bars indicate median with interquartile range (IQR). n = 15 fields of view across 3 independent biological replicates. **(G)** Representative RNA-FISH images of 16S rRNA (magenta) in cells expressing PspCas13b-EGFP (green). Merged images and line-scan intensity profiles demonstrate spatial co-enrichment of Cas13b assemblies with rRNA-rich regions. Scale bar = 1 μm. **(H–I)** Quantification of spatial overlap between PspCas13b-EGFP and 16S rRNA using Manders’ coefficient (H) and Pearson’s correlation coefficient (I). Each point represents the mean coefficient calculated per field of view. Horizontal bars indicate median with IQR. n = 15 fields of view across 3 independent biological replicates. **(J)** Representative time-lapse snapshots of fluorescence recovery after photobleaching (FRAP). A polar focus (white circle) was selectively photobleached and recovery monitored over time. Scale bar = 1 μm. **(K)** FRAP–FLIP analysis of Cas13b mobility between intracellular poles. Fluorescence recovery curves show signal recovery at the bleached pole (magenta), fluorescence loss at the adjacent pole (teal), and total cellular fluorescence (blue). The Soumpasis model fit is shown in green. The mobile fraction (A) was ∼0.40 with a recovery half-time (Td) of ∼30.7 s (n = 39 cells).

Together, these results indicate that *Psp*Cas13b assemblies are dynamic and that *Psp*Cas13b molecules can exchange between spatially distinct intracellular foci. Collectively, the localization, RNA association and FRAP dynamics observed indicate that *Psp*Cas13b preferentially partitions into RNA-rich microenvironments in bacteria while remaining dynamic.

### PspCas13b retains catalytic activity in condensates

While our previous results demonstrate that Cas13 localises to RNA-rich cytoplasmic condensates, it remained unclear whether this localisation has functional consequences for Cas13 activity. Biomolecular condensates can either sequester proteins and RNAs or enhance biochemical reactions by increasing local molecular concentration (Banani et al., 2017; Banani et al., 2016; Cochard et al., 2022; Li et al., 2012; Maharana et al., 2018). We therefore asked whether *Psp*Cas13b retained catalytic activity within these assemblies.

To address this, we first examined whether Cas13b and its RNA substrate co-partition within the same condensates. HEK293T cells were co-transfected with plasmids encoding *Psp*Cas13b-BFP-FLAG, an mCherry reporter transcript, and either a non-targeting (NT) or mCherry-targeting crRNA **(Figure 6A)**. RNA-FISH targeting the mCherry transcript revealed enrichment of mCherry mRNA within these cytoplasmic condensates, as confirmed by line-scan intensity analysis across condensates **(Figure 6B-C)**. Quantification of fluorescence intensities showed increased mCherry mRNA signal within condensates relative to the surrounding cytoplasm, whereas mCherry protein did not display comparable enrichment **(Figure 6D)**. These observations indicate that Cas13 and its RNA substrate can co-localise within condensates.

**Figure 6.**
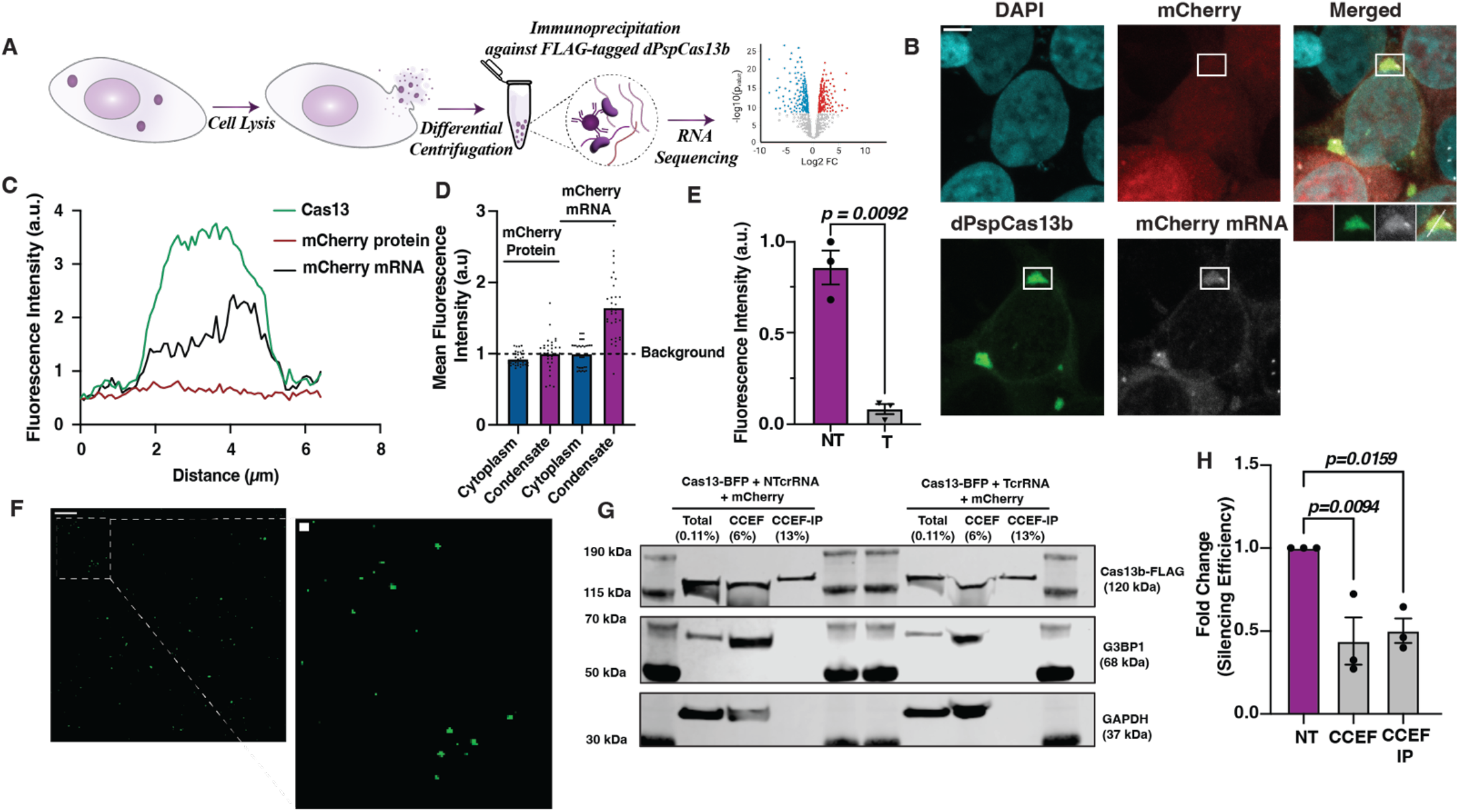
Isolation of dpspCas13b condensates show evidence of mRNA cleavage indicative of Cas13 activity. **(A)** Schematic of isolation protocol of cytoplasmic granules from HEK293T cells. **(B)** RNAFISH image of mCherry mRNA localisation within dpspCas13b condensates. Scale bar =10 μm. **(C)** Fluorescence intensity profile of mCherry mRNA enrichment within the condensate. **(D)** Quantification of Mean fluorescent intensity of mCherry mRNA and mCherry protein (n= 30). **(E)** Bar chart representing normalised mean fluorescence from 3 representative field of views per experiment imaged, n=3. The data is represented in arbitrary units (a.u.). Error bars are standard error of the mean and p-values of one-way Anova test are indicated (95% confidence interval). **(F)** Confocal representative image of granules in organelle enriched cytosolic extract (CCEF). Scale bar = 10μm. Dotted box represents magnified field of view. Scale bar = 1 μm. **(G)** Western blot showing immunoprecipitation of PspCas13b-BFP-FLAG. **(H)** RT-qPCR assay measuring silencing of mCherry mRNA as an indicator for mRNA cleavage within different fractions. n=3. Data are normalised means and errors are SEM. Results were analysed using two-way Anova with p-values indicated (95% confidence interval).

We next tested whether condensate localised *Psp*Cas13b remain catalytically active. Cells expressing mCherry and *Psp*Cas13b with either NT or targeting crRNA were subjected to a cytoplasmic condensate isolation workflow to generate a Cas13 condensate-enriched fraction (CCEF) using differential centrifugation (Hubstenberger et al., 2017; Khong et al., 2018).

FLAG-tagged PspCas13b was subsequently immunoprecipitated from this fraction (CCEF-IP) for downstream analysis **(Figure 6A)**. Coomassie staining and immunoblotting confirmed enrichment of *Psp*Cas13b-BFP-FLAG in the CCEF-IP fraction with minimal background **(Figure 6G; Supplementary Figure 6C)**. Notably, the stress granule marker G3BP1 was not detected in the immunoprecipitated fraction, indicating that although G3BP1 co-localises with *Psp*Cas13b by immunofluorescence, this association does not represent a stable biochemical interaction under the isolation conditions used. To assess *Psp*Cas13b catalytic activity, mCherry transcript abundance was quantified by RT-qPCR using primers spanning the crRNA-targeted region. In the targeting condition, mCherry RNA levels were significantly reduced relative to NT controls across multiple fractions, including the CCEF-IP fraction **(Figure 6H; Supplementary Figure 6D)**. These results indicate depletion of intact target RNA consistent with Cas13-mediated cleavage.

Together, these findings demonstrate that Cas13 recovered from condensate-enriched fractions retains target-dependent catalytic activity. This suggests that localisation within RNA-rich condensates does not sequester Cas13 away from its substrates but instead permits active RNA cleavage within these compartments.

### Cas13 localisation generates distinct RNA association profiles

Previous studies have shown that RNA populations associated with cytoplasmic condensates can be identified through immunoprecipitation coupled with RNA sequencing (Hubstenberger et al., 2017; Khong et al., 2018). We therefore asked **(i)** whether RNA is recovered with d*Psp*Cas13b from cytoplasmic and condensate-enriched fractions, and **(ii)** which transcripts comprise these recovered RNA pools.

To address this, HEK293T cells expressing d*Psp*Cas13b-2xmNeonGreen and a non-targeting crRNA (NTcrRNA) were subjected to biochemical fractionation followed by d*Psp*Cas13b-FLAG immunoprecipitation **(Figure 7A)**. Four RNA populations were analysed: total cytoplasmic RNA (Total), RNA associated with cytoplasmic Cas13 (Total-IP), the condensate-enriched fraction (CCEF), and RNA associated with condensate-localised Cas13 (CCEF-IP). Immunoblotting confirmed robust enrichment of FLAG-tagged dPspCas13b in immunoprecipitated fractions, with no detectable co-purification of condensate-associated proteins (G3BP1, UBAP2L) and absence of the cytoplasmic marker β-actin, supporting specificity of the fractionation and pulldown workflow **(Figure 7B)**. As an orthogonal specificity control, anti-FLAG immunoprecipitation performed in cells expressing HA-tagged dPspCas13b failed to recover Cas13 protein or RNA, confirming that RNA recovery is dependent on FLAG-mediated Cas13 pulldown **(Supplementary Figure 7A)**. RNA integrity and recovery from IP fractions were further validated by TapeStation and RT–qPCR analyses **(Supplementary Figure 7C-D)**.

**Figure 7.**
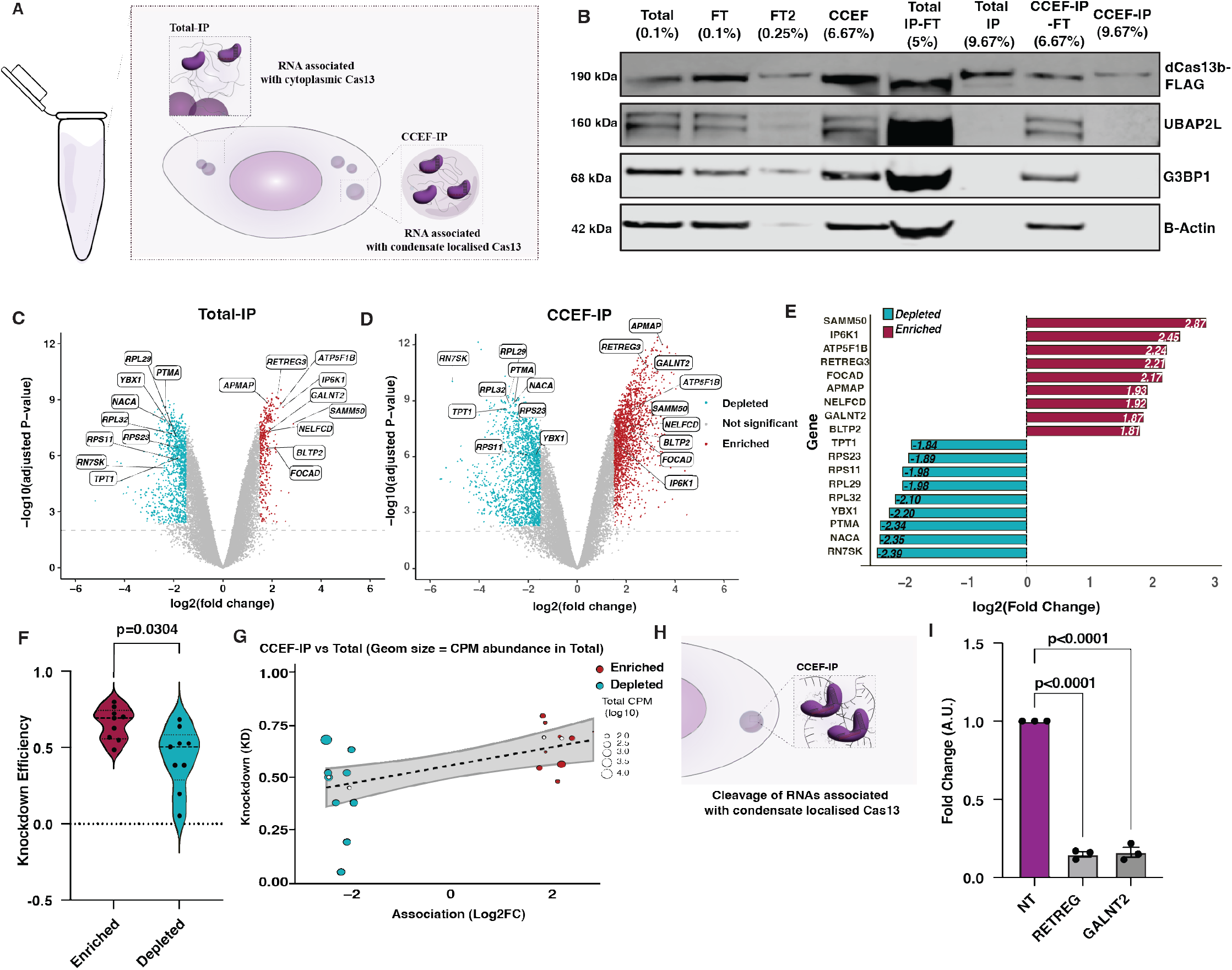
Cas13-RNA association profiles reveal preferential interaction with a distinct subset of transcripts that predicts silencing efficiency. **(A)** Schematic of biochemical fractionation workflow used for Ribo-depletion RNA sequencing. RNA associated with cytoplasmic Cas13 was recovered by immunoprecipitation from total lysate (Total-IP), while RNA associated with condensate-localised Cas13 was recovered from the Cas13 condensate-enriched fraction followed by immunoprecipitation (CCEF-IP). **(B)** Representative immunoblot of FLAG-tagged dPspCas13b-2xmNeonGreen across fractionated samples. Lanes correspond to different fractions of the granule isolation workflow. FT = flow-through; CCEF = Cas13 condensate-enriched fraction. Immunoblotting was performed against the RNA-binding proteins G3BP1 and UBAP2L, with β-actin as a cytoplasmic marker. **(C–D)** Differential RNA association identified by RNA-seq. Volcano plots comparing (C) Total-IP vs Total cytoplasmic RNA and (D) CCEF-IP vs Total RNA. Transcripts significantly enriched are shown in red and depleted transcripts in cyan (adjusted *p* < 0.01, |log_2_FC| > 1.5). **(E)** Log_2_(fold change) values for the top 10 enriched and top 10 depleted transcripts identified in both IP fractions relative to total cytoplasmic RNA. **(F)** Distribution of Cas13-mediated knockdown efficiencies for transcripts enriched or depleted in Cas13-associated fractions. **(G)** Relationship between RNA association with Cas13 (log_2_FC) and observed knockdown efficiency for selected transcripts. **(H)** Schematic illustrating the experimental workflow used for the validation silencing assay. **(I)** RT-qPCR validation of selected transcripts following immunoprecipitation from the condensate-enriched fraction. Silencing of RETREG and GALNT2 indicates Cas13-mediated cleavage of RNAs associated with condensate-localised Cas13. Error bars represent SEM, *n* = 3 biological replicates.

Ribo-depleted RNA sequencing identified selective enrichment and depletion of transcripts associated with dPspCas13b across biochemical fractions **(Figure 7C-D)**. Replicates showed high reproducibility across all sample types **(Supplementary Figure S8A)**.

Differential expression analysis revealed distinct RNA association profiles between cytoplasmic Cas13 (Total-IP) and condensate-localised Cas13 (CCEF-IP). Notably, Cas13 recovered from condensate-enriched fractions was associated with a broader and more strongly enriched set of transcripts compared with cytoplasmic Cas13 **(Figure 7C-D)**. Additional comparisons across biochemical fractions (CCEF-IP vs Total, CCEF-IP vs CCEF, and CCEF-IP vs Total-IP) further confirmed that Cas13-associated RNA populations vary as a function of subcellular localisation **(Supplementary Figure S8-S9)**. Heatmap visualisation of enriched and depleted transcripts showed clear segregation between RNA populations associated with cytoplasmic versus condensate-localised Cas13 **(Supplementary Figure S9)**. Together, these analyses demonstrate that Cas13 associates with a structured and localisation-dependent subset of cellular RNAs in mammalian cells.

### Cas13-associated RNA enrichment correlates with transcript silencing efficiency

We next asked whether Cas13-associated RNA enrichment correlates with functional cleavage efficiency. Candidate targets were selected based on consistent classification as enriched or depleted across both immunoprecipitation fractions and sufficient baseline expression to enable reliable quantification.

Across this panel, transcripts enriched in dPspCas13b immunoprecipitation fractions exhibited greater silencing than depleted transcripts, indicating that Cas13-associated RNAs are preferentially targeted for cleavage. When summarised by group, enriched transcripts showed significantly higher knockdown efficiency compared to depleted transcripts (66% vs 43%, **Figure 7F)**. Importantly, this relationship was not explained by baseline transcript abundance, as low-abundance transcripts that were enriched in Cas13 IP fractions were also efficiently silenced.

To directly assess whether cleavage occurs within condensate-associated Cas13 material, we examined two enriched targets, RETREG and GALNT2. RT–qPCR analysis of RNA recovered from the condensate-associated immunoprecipitated fraction (CCEF-IP) showed marked depletion of both transcripts (∼85% and ∼84% knockdown, respectively) relative to non-targeting controls (Figure 7H). These results provide evidence consistent with active RNA cleavage within Cas13-containing condensates.

Together, these findings demonstrate that Cas13-associated RNA enrichment is predictive of cleavage efficiency, linking spatial RNA association with functional outcome.

### Transcript features of Cas13-associated RNAs reveal bias in biotype and transcript architecture

We next asked whether transcripts enriched in Cas13-associated fractions share common structural features. Transcript annotation revealed that the majority of enriched RNAs were protein-coding transcripts, whereas depleted sets showed a broader representation of non-coding RNA biotypes **(Supplementary Figure S11A)**. In addition, enriched transcripts were generally longer and contained longer coding sequences and extended 3′ untranslated regions, whereas 5′ UTR length showed little difference between enriched and depleted sets **(Supplementary Figure S11B-D)**. These trends were observed across both cytoplasmic and condensate-associated IP fractions. Together, these results suggest that Cas13 association is influenced by broad transcript features, and that localisation within RNA-rich condensates reshapes the RNA populations accessible to Cas13.

### Surface electrostatics of Cas13 suggest a structural basis for broad RNA association

The transcriptomic analyses above revealed reproducible association of Cas13 with a selective subset of cellular RNAs across both cytoplasmic and condensate-associated contexts **(Figure 7)**. These observations raised the possibility that intrinsic structural features of Cas13 may contribute to RNA association beyond canonical crRNA-target recognition. To explore this, we generated a structural model of PspCas13b using AlphaFold2 and calculated electrostatic surface (Jumper et al., 2021; Jurrus et al., 2018). Electrostatic mapping revealed multiple positively charged surface patches distributed across the protein surface, including a prominent positively charged groove-like region **(Figure 8A)**. This feature is consistent with guide RNA-binding regions described in structural studies of Cas13 enzymes (Ishikawa et al., 2025; Liu et al., 2017; Zhang et al., 2018). In contrast, negatively charged regions were distributed across the remaining surface, producing a patchy and asymmetric electrostatic landscape rather than a uniformly charged surface **(Figure 8A)**.

**Figure 8.**
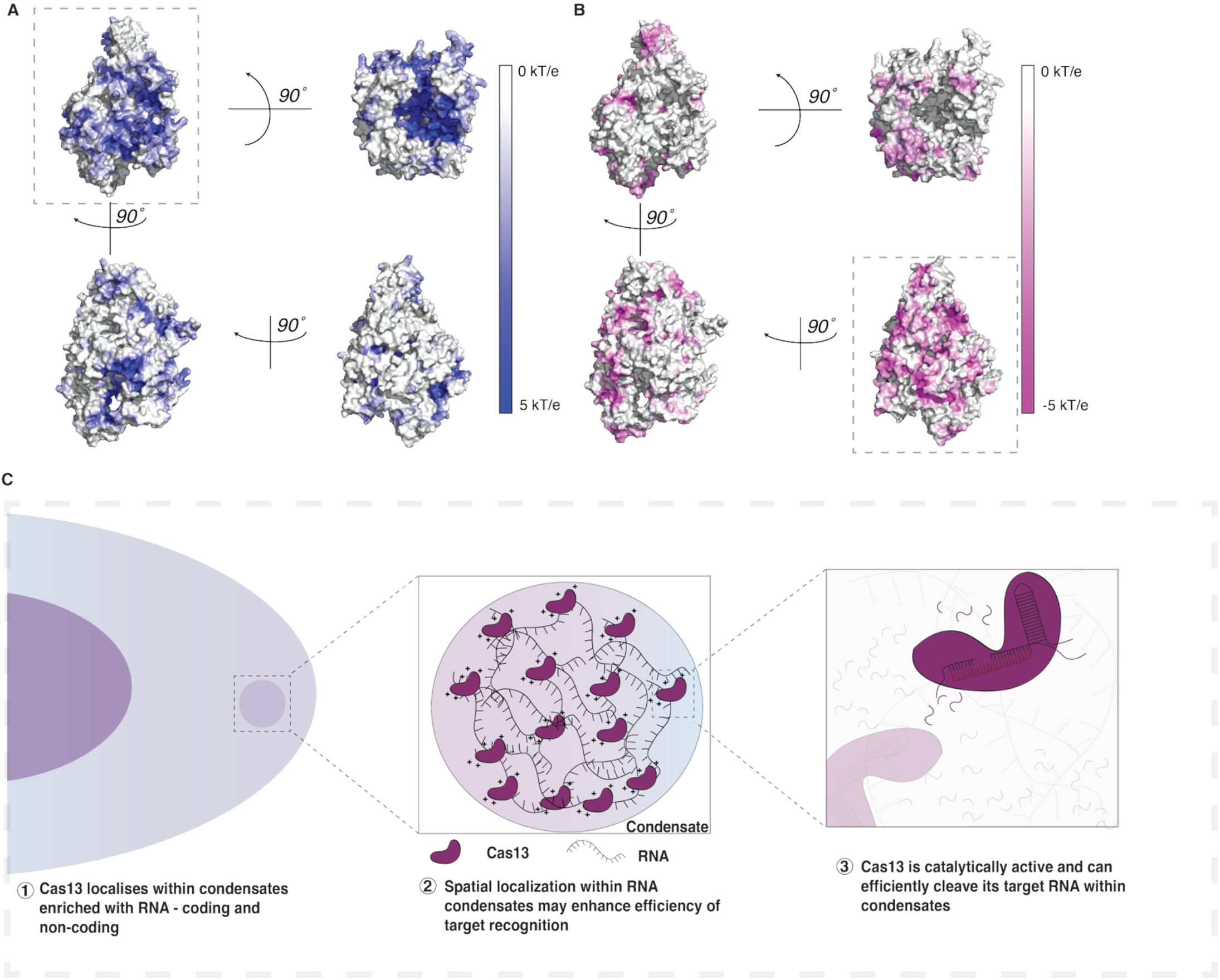
Spatial model for Cas13 RNA targeting in RNA-rich assemblies. **(A)** Electrostatic surface representation of PspCas13b predicted using AlphaFold2 and calculated using APBS in PyMOL. Structures are shown in multiple orientations (90° rotations) to visualise surface charge topology. Positively charged regions are shown in blue (+5 kT/e) and negatively charged regions in magenta (−5 kT/e). The protein surface displays an asymmetric distribution of electrostatic charge with multiple positively charged surface patches, including a prominent groove consistent with previously described RNA-binding regions of Cas13 enzymes. **(B)** Electrostatic surface representation of PspCas13b highlighting negatively charged regions (magenta). The overall electrostatic landscape is patchy and non-uniform, suggesting the potential for transient electrostatic interactions with RNA molecules. **(C)** Proposed model for Cas13 partitioning into RNA-rich assemblies. Cas13 proteins preferentially localise to RNA-rich cytoplasmic condensates where elevated local RNA concentration facilitates transient RNA interactions. Within these assemblies Cas13 remains catalytically active and can efficiently engage and cleave RNA substrates enriched in these compartments.

To assess whether these features are unique to *Psp*Cas13b or reflect a broader property of Cas13 effectors, we performed analogous analyses on additional Cas13 orthologues. *Lw*Cas13a and *Rfx*Cas13d similarly exhibited asymmetric surface charge distributions with prominent positively charged regions suggesting that electrostatic features compatible with RNA interaction may be shared across Cas13 subtypes **(Supplementary Figure S14)**.

These analyses provide a structural context compatible with the RNA association profiles observed experimentally, but do not establish a causal mechanism. Together with the biochemical and transcriptomic data, these findings support a model in which Cas13 partitioning into RNA-rich environments increases the likelihood of productive interactions with associated transcripts **(Figure 8C)**.

## Discussion

While Cas13 activity is widely attributed to crRNA-target complementarity, the role of intracellular spatial localisation has remained largely unexplored. Here, we show that intracellular spatial organisation of Cas13 constitutes an additional layer of regulation, governing target accessibility and nuclease activity. Across both mammalian and bacterial cells, Cas13 preferentially localises within RNA-dense granules rather than remaining uniformly distributed within the cytoplasm.

In mammalian cells, d*Psp*Cas13b is diffusely distributed when expressed alone but is recruited into distinct cytoplasmic granules upon crRNA co-expression. The dose-dependent formation of these condensates – observed with both plasmid-encoded and in vitro-transcribed crRNA, indicates that intracellular crRNA abundance nucleates RNA condensate assembly, which in turn drives the recruitment of Cas13 to these foci. This is consistent with the principle that RNA can self-assemble into condensate-nucleating scaffolds through RNA-RNA and RNA-protein interactions in a concentration-dependent manner (Banani et al., 2017; Guillén-Boixet et al., 2020; Van Treeck et al., 2018; Wadsworth et al., 2024). Live-cell imaging and FRAP analyses demonstrated that these assemblies exhibit hallmarks of condensates, including fusion and fission behaviour, molecular exchange with the surrounding cytoplasm, and fluorescence recovery kinetics consistent with liquid-like material properties (Lyon et al., 2021; Sprague & McNally, 2005; Wang et al., 2022). d*Psp*Cas13b granules dissolved during mitosis and reformed in daughter cells, a hallmark behaviour that distinguishes phase-separated condensates from stable protein aggregates (Rai et al., 2018; Valverde et al., 2023). Notably, these condensates display markers of stress granules including G3BP1 and UBAP2L. However, Cas13 condensates persisted despite the ablation of G3BP1/2, indicating that these are non-canonical stress granule condensates. This is consistent with emerging evidence that biomolecular condensates can be induced through multiple redundant pathways, including elevated intracellular RNA concentration (N. Kedersha et al., 2016; Roden & Gladfelter, 2021).

This localisation behaviour was conserved across multiple Cas13 orthologues. *Lw*Cas13a, *Psp*Cas13b, and *Rfx*Cas13d all enriched within RNA-rich cytoplasmic condensates despite sharing only ∼16 –18% pairwise amino acid identity outside their conserved HEPN domains, implying that condensate partitioning is governed by a general physicochemical property, such as the capacity to associate with RNA-dense environments through surface charge and multivalent RNA contacts, rather than any conserved sequence motif. Consistent with this, *Psp*Cas13b exhibited comparable localisation behaviour in *E. coli*, displaying polar, membrane-associated, and multipunctate foci enriched in 16S rRNA, which showed dynamic behaviour similar to RNA condensates observed in mammalian cells (Guo et al., 2024; Monterroso et al., 2024; Passos et al., 2025; Pei et al., 2025). The observation that Cas13 partitions into RNA-rich microenvironments across both prokaryotic and eukaryotic contexts reinforces the idea that this behaviour may be an intrinsic property encoded in Cas13’s structure and surface properties.

From a functional perspective, localisation within RNA-dense condensates hubs in prokaryotes could increase the likelihood of encountering viral transcripts during infection, thereby enhancing Cas13-mediated immune surveillance. This is particularly pertinent given that viral infections themselves trigger the assembly of RNA-rich condensates in both prokaryotic and eukaryotic cells, including stress granules, RNase L-induced bodies, and viral replication factories, creating discrete RNA-dense compartments at sites of active infection (Al-Husini et al., 2018; Burke, 2023; Castellana et al., 2016; Nandana et al., 2023).

RNA enrichment within these assemblies places Cas13 in compartments containing abundant potential substrates. Biomolecular condensates are increasingly recognised as microenvironments that reshape biochemical reactions by altering local concentration, diffusion, and encounter probability (Banani et al., 2017; Holehouse & Alberti, 2025; Peeples & Rosen, 2021; Riback et al., 2020). For an RNA-guided nuclease, localisation within an RNA-dense phase may therefore influence not only where the enzyme resides but which transcripts it samples and how frequently productive interactions occur. Interestingly, in mammalian RNA interference, Argonaute-containing silencing complexes partition into RNA-rich cytoplasmic assemblies, enhancing small RNA-mediated repression (Liu et al., 2005; Sheu-Gruttadauria & MacRae, 2018). Our findings suggest that Cas13 may similarly exploit localisation within RNA-rich condensates to facilitate target search and cleavage.

This localisation within RNA-rich condensates has potential implications for the downstream fate of its cleavage products. Beyond their role in RNA storage and translational regulation, condensates concentrate a diverse array of RNA-binding proteins and RNA metabolism factors, including components of the RNA decay, surveillance, and repair machinery (Banani et al., 2017; Ivanov et al., 2019; Ouyang et al., 2019; Pessina et al., 2025). Recent work has demonstrated that RNA cleavage by CRISPR nucleases generate defined RNA ends that are subject to cellular repair. Nemudraia and colleagues showed that RNA fragments produced by type III CRISPR-mediated cleavage are repaired in human cells, highlighting RTCB (HSPC117) – the only well-characterised eukaryotic 3′-5′ RNA ligase, with established roles in tRNA maturation and XBP1 mRNA splicing, as a plausible ligase for sealing the relevant RNA ends (Moncan et al., 2023; Nemudraia et al., 2024). In addition, *Psp*Cas13b targeting has been reported to generate unexpected RNA nicking and ligation events near cleavage sites, consistent with downstream cellular processing of Cas13-generated RNA ends, a finding that has been exploited to develop RNA segment editing as a therapeutic strategy (Hu et al., 2025; Lam et al., 2026). Our data indicates that endogenous RTCB is detectable within d*Psp*Cas13b condensates **(Supplementary Figure S15)**, consistent with prior reports of RTCB co-partitioning with stress granule-associated RNA granules (Ning et al., 2019). The co-concentration of Cas13 with RNA processing machinery such as RTCB, raises the possibility that the condensate microenvironment may shape the downstream fate of Cas13 cleavage products, whether through ligation-mediated repair, decay, or other RNA processing events. While prior mechanistic studies of Cas13 have largely focused on the rules of target recognition and catalytic activation *in vitro* (Abudayyeh et al., 2017; East-Seletsky et al., 2016; Hu et al., 2024), our findings suggest that the intracellular spatial context of target search is non-random and systematically biased by condensate partitioning. Transcriptomic profiling revealed a Cas13-RNA association profile in which transcripts enriched within Cas13-containing condensates exhibited greater silencing than transcripts depleted from these fractions, directly linking spatial localisation to functional outcome. Condensate-enriched transcripts tended to be longer with extended coding sequences and 3′ UTRs, consistent with known enrichment of long poorly translated mRNAs in stress granule-like condensates, suggesting that transcript physicochemical properties may contribute to Cas13 encounter probability independently of crRNA design (Khong et al., 2017).

The molecular basis of Cas13 recruitment to RNA-rich environments remains unclear.

Our structural modelling identified positively charged surface patches on *Psp*Cas13b, suggesting that electrostatic interactions with RNA may contribute to condensate partitioning, a mechanism observed for other RNA-binding proteins that associate with biomolecular condensates through charge-mediated non-specific contacts (Dutagaci et al., 2021; Sesé-Sansa et al., 2022; Vernon et al., 2018). Consistent with this, Cas13 immunoprecipitation recovered a broad population of RNAs in the absence of a targeting crRNA, suggesting capacity for sequence-nonspecific RNA association.

Taken together, these findings support a two-layer model of Cas13 function in cells. crRNA-target complementarity defines which transcripts are targetable, whereas intracellular RNA organisation determines which transcripts are encountered. Cas13 therefore operates within a spatially structured transcriptome in which RNA-rich microenvironments bias encounter probability and reshape effective target accessibility. More broadly, these findings indicate that the activity of programmable RNA-targeting nucleases is shaped not only by the rules of molecular base pairing, but also by the spatial architecture of the cellular transcriptome it navigates.

## Author Contributions

G.K.G.J.S. and M.F. conceived the study. M.F., I.V., P.M., and J.T. supervised the study. G.K.G.J.S. and M.F. designed the experiments. G.K.G.J.S. performed all experiments and analysed the data. A.H. set up the analysis for the FRAP experiments. W.H. and C.S. provided training in experimental techniques and cloning. P.M. provided microscopy expertise and infrastructure support. G.K.G.J.S. generated the figures with input from M.F. G.K.G.J.S. and M.F. wrote the manuscript. All authors read, commented on, edited and approved the manuscript.

## Acknowledgments

The authors thank all lab members from the Fareh and Trapani lab for facilitating experiments and discussions. We also acknowledge the Centre for Advanced Histology and Microscopy (CAHM) at the Peter MacCallum Cancer Centre for providing all microscopy infrastructure.

## Data Availability

All data are available in the main text and/or supplementary materials. Source data are provided with this paper. All key plasmids constructed in this study will be deposited to Addgene upon publication. The reference human transcriptome (GRCh38) was used for read alignment. Raw RNA sequencing data will be deposited to NCBI Gene Expression Omnibus (GEO) upon publication (accession pending). Raw RNA-seq data, processed differential expression data and analysis code are publicly available at https://github.com/FarehLab/Cas13IP-RNA-Seq-Profiles, with an interactive data explorer accessible at https://farehlab.github.io/Cas13IP-RNA-Seq-Profiles/.

## Supplementary Data

Supplementary data are available online.

## Materials and Methods

### Cell Culture

HEK 293T (ATCC CRL-3216), HCT-116 (ATCC-CCL-247) and HeLa (ATCC-CCL-2) cell lines were cultured in DMEM high glucose media (Thermo Fisher, 11965092) containing 10% or 5% heat-inactivated fetal bovine serum (Thermo Fisher, 10100147), 100mg/ml Penicillin/-Streptomycin (Thermo Fisher, 730 151401220), and 2mM GlutaMAX (Thermo Fisher, A1286001). The cell lines were incubated at 37 °C and 10% CO_2_. Cells were passaged twice a week and were routinely tested for mycoplasma.

### crRNA design and cloning

*Psp*Cas13b crRNA spacers were designed as 34-nt single-stranded forward and reverse oligos containing CACC and CAAC overhangs, respectively, allowing for ligation into BbsI-digested plasmid that encodes the *Psp*Cas13b direct repeat (Addgene #103854). Oligos were ordered as single-stranded DNA (IDT) at a concentration of 100 μM. All crRNA spacer sequences used in this study are available in **(Table S1)**. To produce double-stranded DNA oligos suitable for plasmid ligation, each forward and reverse DNA oligos (3 μM) were annealed in a T4 DNA ligase buffer (NEB) by heating to 95°C for 5 min then cooling at a rate of 4°C every 2 min until ambient temperature was reached. The *Psp*Cas13b crRNA backbone plasmid (Addgene #103854), a gift from F. Zhang, was digested using BbsI (NEB) in 1x NEBuffer r3.1 at 37°C for 1 hour. Digested plasmids were separated on a 1% (w/v) agarose gel in 1x TAE (tris-acetate-EDTA) buffer at 100 V for 1 hour. The linearized fragment was then excised and purified using the NucleoSpin Gel and PCR Clean-up Kit (Macherey-Nagel). The annealed oligos (400 nM) were ligated with 0.25ng of the digested crRNA backbone using T4 ligase (Promega) in 1x T4 ligase buffer for 3 hours at room temperature (RT)

### Plasmid amplification and purification

Five microliters of the ligation product or 50 ng of the circularized plasmid was transformed into 25 μl of Top10F or STBL3 competent bacteria by heat shock at 42°C for 45s, followed by 2 min on ice. The transformed bacteria were recovered in 450 μl of LB media for 1 hour at 37°C in a shaking incubator (200 rpm). The bacteria were pelleted by centrifugation at 8000 rpm for 30 secs, resuspended in 100 μl of LB broth, and plated onto prewarmed LB agar plates containing ampicillin (75 μg/ml). Following overnight incubation at 37°C, single colonies were picked and transferred into 3-ml bacterial starter cultures and incubated for ∼16 hours prior to mini- or maxi-prep DNA purification using the NucleoSpin Plasmid kit (Macherey-Nagel). Successful cloning of all plasmids was verified by Sanger sequencing (Australian Genome Research Facility).

### Transfections Assays for Cas13 visualisation and RNA silencing assays

Transfection experiments were performed using an optimised Lipofectamine-3000 transfection protocol (Life Technologies, L3000015). Transfection assays were carried out as follows: 100,000 cells/500 μL were seeded in 24-well tissue culture treated flat-bottom plates (Corning) ∼18 hours prior to transfection on poly-l-lysine coated coverslips. For each well, a total of 200ng DNA plasmid (100ng of pspdCas13b Addgene#132403, 100ng crRNA Addgene#103854) was mixed with 1 μL of p3000 in 25 μL Opti-MEM Serum-free Medium (Master-Mix 1) (Gibco). In a separate tube, 25 μL Opti-MEM was mixed with 1.5 μL Lipofectamine-3000 (Master -2). Master mixes 1 and 2 were then added together and incubated for 20 minutes at room temperature. ∼50 μL of the transfection mixture was added to each well.

For RNA silencing assays, transfection was carried out as follows: 150,000 cells/500 µL were seeded in 24-well tissue culture treated flat-bottom plates. For each well. A total of 460ng of DNA plasmids (100ng PSpCas13b-NES-3xFLAG-T2A-BFP Addgene#173029, 100ng of crRNA, 260ng MSCV-mCherry Addgene#52114) was mixed with 1 µL of P3000 in 25 µL Opti-MEM Serum-free Medium. In a separate tube, 25 μL Opti-MEM was mixed with 1.5 μL Lipofectamine-3000 (Master -2). Master mixes 1 and 2 were then added together and incubated for 20 minutes at room temperature. ∼50 μL of the transfection mixture was added to each well. After transfection, cells were incubated at 37 °C. 10% CO_2_ and the transfection efficiency as well as mCherry mRNA silencing was monitored 24-72 hours post transfection by fluorescent microscopy.

### Cloning of Cas13 orthologues to produce mNeonGreen fusion constructs

Cas13 orthologues dRfxCas13d and dLwCas13a was cloned into a plasmid encoding two mNeonGreen fluorescent proteins (Addgene #132403). The backbone was linearised via double digest using Not1 and Xba1 enzymes. The inserts/fragments were obtained using PCR-based cloning using primers that flank the insert. drfxCas13d was obtained from Addgene#132411, and dLwCas13a was obtained from Addgene#91905. All PCR reactions were done using Q5® High-Fidelity DNA Polymerase (NEB M0491S). The assembly of products was carried out using Gibson Assembly (NEB #E5510S) following manufacturers protocol. Colony PCR was carried out to screen for positive colonies and successful cloning was verified using Sanger Sequencing.

### Western Blotting

Cells were washed with ice-cold PBS ± and lysed on ice in RIPA lysis 762 buffer [50 mM Tris (Sigma-Aldrich, T1530), pH 8.0, 150 mM NaCl, 1% NP-40 (Sigma Aldrich, 118896), 0.1% SDS, 0.5% sodium deoxycholate (Sigma-Aldrich, #D6750)] containing protease inhibitor cocktail (Roche, #04693159001) and phosphatase inhibitor cocktail (Roche, #4906845001). Samples were incubated for 30min at 4 °C with rotation (25 rpm), and centrifuged at 16,000 g for 10 min, 4 °C. Supernatant was transferred to a new tube. Protein concentrations were quantified using the Pierce 768 BCA Protein Assay Kit (Thermo Fisher, 23225) according to the manufacturer’s instructions. A total of 20 μg of protein diluted in 1x Bolt LDS sample buffer (Thermo Fisher, B007) and 1x Bolt sample reducing agent (Thermo Fisher, B009) were denatured at 95 °C for 5 min. Samples were resolved by Bolt Bis-Tris Plus 4–12% gels (Thermo Fisher, NW04120BOX) in 1x MES SDS running buffer (Thermo Fisher, B0002) for 1.5 hrs at 100V. Following electrophoresis, proteins were transferred onto 0.45 μM PVDF membranes (Thermo Fisher, 88518) activated in 100% methanol for 5 seconds, rinsed with MilliQ H_2_O, then soaked in 1x transfer buffer (Tris-HCl 23mM [pH 7.3], glycine 192mM, 20% v/v methanol) using a Trans-Blot Semi-Dry electrophoretic transfer cell (Bio-Rad, CA, USA) apparatus at 20 Volt for 30 min. After transfer, membranes were blocked in 5% (w/v) BSA (Sigma-Aldrich, A3059) in TBST with 0.15% 780 Tween 20 (Sigma-Aldrich, P1379) for 1h at RT and probed overnight with primary antibodies at 4 °C. Blots were washed three times in TBST with 0.15% Tween20, 782 followed by incubation with fluorophore-conjugated secondary antibodies diluted in intercept Blocking Buffer (LI-COR) for 1h at RT. Membranes were washed in TBST (0.15% Tween20) three times and fluorescence was detected using the Odyssey CLx 785 Imager 9140 (Li-COR).

### Immunofluorescence

#### Mammalian Cells

Immunofluorescence was carried out to visualise the distribution of psp-Cas13b-BFP-3xFLAG. Fixation was carried out by incubating cells in 4% PFA in PBS for 5 minutes at room temperature. Cells were then washed three times with ice-cold PBS and permeabilization was then carried out by incubating cells in 100% ethanol at -20 °C overnight. The cells were washed with PBS three times and permeabilised with 0.1% Triton X-100 in PBS for 10 minutes at room temperature. Cells were then washed with PBS three times for 5 minutes. Cells were then blocked with 3% bovine serum albumin (BSA) (0.3g BSA in 10ml PBST (PBS + 0.1% Tween 20)) for 30 minutes. Cells were then incubated in the appropriate primary antibodies overnight at 4°C. The following day, the solution was aspirated, cells were washed three times in PBS for 5 minutes and were incubated with appropriate secondary antibodies for an hour at room temperature in the dark. The coverslips were then washed with PBS three times for 5 minutes, counter stained with DAPI and mounted using ProLongTM Glass Antifade Mountant (P36980). Samples were incubated for 30 minutes at 37°C to allow drying and polymerization of mounting media, and the coverslips were sealed using nail polish.

#### Bacteria Cells

Immunofluorescence was carried out to visualise PspCas13b (Addgene#115219) localisation within BL21-DE3 *E*.*Coli* Cells (Thermofisher, C600003). A fresh colony was inoculated into 7mL of Terrific Broth medium containing 7 µL of Ampicillin (1:100) and was left to shake overnight at 200rpm at 37 °C. Dilute cells to reach OD_600_ of 0.01. Cool the cultures to 18 °C and add IPTG to final concentration of 0.05mM or 0.1mM. to induce expression of pspCas13b. (-IPTG was used as a negative control). Incubate tubes at 18 °C at 200rpm overnight. Once cells have reached OD_600_ of 0.5, the cells were fixed with 4% PFA for 30 minutes at R.T.P. Cells were then permeabilised with 70% ethanol at -20 °C overnight. Cells were then treated with Lysozyme (25 µg/mL in TEG buffer (5mM Tris (HCl), 10mM EDTA, 50mM glucose, pH 8.0)) for 30 minutes at R.T.P. Immunostaining was carried out by first blocking cells in blocking buffer (0.1% ultrapure BSA, 0.05% Tween-20, 1x PBS) for 1 hour at 37 °C. Cells were then incubated in primary antibody (Sigma #SAB2702218) diluted in blocking buffer (1:500) and incubated for 1.5h at R.T.P. Cells were then incubated in secondary antibody (1:500) at R.T.P for 1h. The cells were counted stained with DAPI and mounted using ProLongTM Glass Antifade Mountant (P36980).

### RNA extraction, cDNA synthesis, and RT-PCR

Total RNA was isolated from around using TriZol (Invitrogen #15596026) according to manufacturer’s instructions. For RNA extraction to determine silencing efficiency of mCherry mRNA, DNase treatment was carried out with RQ1 RNase-Free DNase according to manufacturer’s instruction (Promega, M6101). RNA was converted to cDNA using the high-capacity cDNA reverse transcription kit (Thermo Fisher, 4368814) following the manufacturer’s instructions. Quantitative RT-PCR reaction was performed in duplicate in a StepOne Real-Time PCR system (Thermo Fisher) using PowerUp™ SYBR™ Green Master Mix (Thermo 799 Fisher, A25742). *GAPDH* was used as a housekeeping gene. The 2^−ΔΔ*C*t^ method was used to normalize the expression of a transcript of interest. Primers for RT–qPCR are detailed in **(Table S2)**.

### Isolation of RNA from HEK293T cells and Granule Cores using Immunoprecipitation

HEK293T cells expressing dPspCas13b-2xmNeonGreen and its cognate crRNA via transient transfection was grown to ∼80% confluency in 10 T175cm^2^ U-shaped Angled Neck Cell Culture Flasks (Corning #431080). Once sufficient granule formation was observed, cells were washed once with media, transferred to falcon tubes, and pelleted at 1500g for 3 mins. Upon aspirating the media, the pellets were flash-frozen in liquid nitrogen and stored at -80 °C until isolation of mammalian Granule cores was performed.

The isolation of granule cores was adapted from (Khong et al., 2017). Briefly, the pellets were thawed in ice, resuspended in 1mL lysis buffer (50mM Tris, pH 7,4, 1mM EDTA, 150mM NaCl, 0.2% Triton X-100) in the presence of 65 U/mL RNaseOut ribonuclease inhibitor (Promega) and EDTA-free protease inhibitor cocktail (Roche, #04693159001). To accelerate lysis, extracts were passed through a 25 Gauge X 1” (0.5mm x 25mm) needle attached to a 2mL syringe 7 times. They lysates were spun at 1000g for 5 mins at 4 °C to pellet cell debris. 50 µL, 150 µL and 800 µL (volumes would vary) of the supernatant was transferred to new microcentrifuge tubes for isolating total RNA (50 µl), RNAs interacting with cytoplasmic Cas13 (150 µL) and RNAs interacting with Cas13 enriched within the granule (800 µL). For isolating total RNA, TRIzol (Invitrogen, #15596018) was added and sequential extraction of both RNA and protein was carried our according to manufacturer’s protocol.

The following steps were performed to isolate mammalian SG cores and extract its RNA. The supernatants were spin at 18,000g for 20 mins at 4 °C to pellet SG cores. The supernatant was discarded, and the pellet was resuspended in 1mL lysis buffer and spun again at 18,000g for 20 mins at 4 °C to enrich for SG cores. The resulting pellet was resuspended in 300 µL lysis buffer and spun at 850g for 2 mins at 4 °C. The supernatant was transferred to a new microcentrifuge tube which represents the mammalian SG enriched fraction (SGEF).

Immunoprecipitation was carried out using Pierce™ Anti-DYKDDDK Magnetic Agarose beads (Invitrogen, #A36797) as per manufacturer’s instructions. Briefly, the beads were allowed to come to room temperature. 100 µL of slurry was used and a MagnaRack (Invitrogen, #CS15000) was used to collect the beads. The supernatant was removed, and the beads were washed twice with 500 µL of lysis buffer. The lysates were added, and the volume was made up to 300 µL. Samples were incubated at 4 °C on a horizontal tube roller for 3 hours. The beads were then washed once with 500 µL lysis buffer and 500 µL of Rnase-free water. Acid elution was carried out using twice using 100 µL of 0.1M glycine, pH 2.8. The beads were incubated for 10 minutes in a thermomixer, and the supernatant was collected. 15 µL of 1M Tris, pH 8.5 was added per 100 µL of eluate. Sequential RNA and protein extraction was carried out from the SG cores using TRIzol (Invitrogen, #15596018) according to manufacturer’s protocol.

### Mammalian library construction and RNA-Sequencing

After RNA isolation from whole cells and granule core fractions, 100 ng of RNA was used for ribosomal RNA-depleted RNA-sequencing. RNA quality was assessed using High-Sensitivity RNA ScreenTape (Agilent, #5067-5579) on the Agilent TapeStation 4150 (Peter MacCallum Molecular Genomics Core Facility). Library preparation was performed using the Illumina Ribo-Zero Plus kit (#20040526) according to the manufacturer’s instructions. Quality control of libraries was performed using TapeStation followed by qPCR-based quantification RNA-sequencing was carried out at the Australian Genome Research Facility (AGRF, Melbourne) on the Illumina NextSeq platform, generating paired-end cDNA libraries. Image analysis and base calling were performed in real-time using NovaSeq Control Software (NCS v1.2.0.28691) and Real-Time Analysis (RTA v4.6.7). Demultiplexing and FASTQ generation were performed using Illumina DRAGEN BLC Convert v07.021.645.4.0.3.Reads were aligned to the GRCh38 reference genome using STAR aligner with default settings.

Gene-level counts were generated using featurecounts, and normalization was performed using the limma-voom pipeline (limma 3.58.1) (Law et al., 2014; Ritchie et al., 2015). Differential expression analysis was conducted between cytoplasmic (Total), total IP, and condensate-enriched IP (CCEF-IP) fractions. Genes with log_2_ fold change (log_2_FC) > 1.5 were considered enriched; those with log_2_FC < –1.5 were considered depleted. Statistical significance was determined using an adjusted p-value < 0.05 (Benjamini-Hochberg method, FDR 5%). Genes highlighted in volcano plots were selected for further validation and functional assays of Cas13-mediated cleavage. To assess transcript-level properties, gene annotations were retrieved using the biomaRt R package (v2.58.2), querying the ENSEMBL Genes 110 (GRCh38.p13) dataset(Aken et al., 2016). Transcript features including total transcript length, CDS length, and UTR lengths were extracted, and biotype classification was based on the most expressed transcript per gene. Comparative analyses between enriched and depleted gene sets were visualized in R using ggplot2 (v3.4.4) for plotting and pheatmap (v1.0.12) for heatmap generation. Statistical significance for differences in transcript features was assessed using two-sided Mann–Whitney U tests. Venn diagrams were generated using the VennDiagram (v1.7.3) package.

### Microscopy Analysis

#### Confocal

Fixed and live cell imaging was carried out using the FLUOVIEW FV3000 Confocal Laser Scanning Microscope (Olympus). All images were taken at 16 bits per pixel with a scan size of 1024 by 1024 with a zoom magnification of 2.5 using a PLAPON60x 1.518 NA objective lens oil. Z stacks varied from cell to cell, but slice size was kept constant from 0.4 to 0.6 um per slice. Mean fluorescence intensity was quantified using FIJI and the data was analysed using GraphPad Prism.

#### Fluorescent Microscopy Analysis

For RNA silencing experiments, the fluorescence intensity was monitored using EVOS 750 M5000 FL Cell Imaging System (Thermo Fisher). Images were taken 48 h post transfection, and the fluorescence intensity of each image was quantified using a lab written macro in ImageJ software. Briefly, all images obtained from a single experiment are simultaneously processed using a batch mode macro. First, images were converted to 8-bit, threshold adjusted, converted to black and white using Convert to Mask function, and fluorescence intensity per pixel measured using Analyze Particles function. Each single mean fluorescence intensity was obtained from four different field of views for each crRNA and subsequently normalized to the non-targeting (NT) control crRNA. Two-fold or higher reduction in fluorescence intensity is considered as biologically relevant.

### Fluorescence Recovery After Photobleaching

FRAP experiments were carried out on a FLUOVIEW FV3000 Confocal Laser Scanning Microscope (Olympus) using filter sets and scanning mirrors provided by the manufacture. Imaging was performed using a PLAPON 60 × 1.518 NA objective lens. The imaging area was further narrowed with a 28x zoom which contained a circular bleach ROI with a radius of 0.5um to 0.9um. Circular bleaching was carried out in 10 different cells with a 488nm argon laser (20mW) using a scan size of 64 by 64 with total frames ranging from 300 to 500 frames at 177.246 frames per msec. Synchronization was done without rest after 20 frames of normal laser scanning confocal imaging to obtain initial intensity measurements. Stimulation (photobleaching) was carried out for a duration of 147.000 msec at 50% 488nm laser intensity. Either the cytoplasm, the whole dCas13b granule, or part of the granule was photobleached. All FRAP experiments were performed at 37 °C with 5% CO2. Intensity profiles were obtained using the z-axis profile tool on ImageJ. The data was graphed using GraphPad PRISM and data fitting was done using a non-linear regression Soumpasis fit. The mobile fraction (Mf) was obtained using the following equation:

The immobile fraction (If) was then calculated as If = 1 – Mf. Diffusion coefficients were obtained using the following equation:

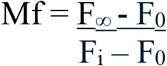

Where;

F_∞_ = fluorescence after recovery,

F_0_ = fluorescence after photobleaching

F_i_ = initial fluorescence

The immobile fraction (If) was then calculated as If = 1 – Mf. Diffusion coefficients were obtained using the following equation:

Td (Half-life of diffusion) = R2/(4*D), where D is the diffusion coefficient and R is the radius.

### Fluorescence Recovery After Photobleaching (FRAP) and Fluorescence Loss in Photobleaching (FLIP) in *BL21-DE3 E*.*Coli Cells*

For bacteria, FRAP and FLIP experiments were carried out on a FLUOVIEW FV4000 Confocal Laser Scanning Microscope (Olympus) using filter sets and scanning mirrors provided by the manufacture. Imaging was performed using a PLAPON 60X 1.518 NA objective lens. The imaging area was further narrowed with a 10x zoom which contained a circular bleach ROI. Circular bleaching was carried out with 488nm argon laser (20mW) using a scan size of 128 by 128 with total frames of 1000 frames at speed 208 frames per msec. Synchronization was done without rest after 20 frames of normal laser scanning at 0.1% 488nm confocal imaging to obtain initial intensity measurements. Stimulation (photobleaching) was carried out for a duration of 40.000 msec at 50% 488nm laser intensity. One polar focus was bleached, and fluorescence intensity was measured in both the bleached and adjacent pole to obtain the fluorescence loss in photobleaching. All experiments were performed at 37 °C. Intensity profiles were obtained using the z-axis profile tool on ImageJ. The data was graphed using GraphPad PRISM and data fitting was done using a non-linear regression Soumpasis fit.

### Single molecule inexpensive fluorescent *in situ* hybridisation (smiFISH)

smiFISH was carried out as described by Mueller and colleagues (Tsanov et al., 2016). Probes were designed according to the Oligostan R script. Sequence of probes listed in **Table S3**. Transfected cells were fixed with 4% Paraformaldehyde, and permeabilised overnight in 100% ethanol at - 20 °C. Cells were then rinsed with PBS three times, and RNAse treatment was carried out by incubating samples in 1x RNase A (1mg/mL) for 30 minutes at room temperature on samples for the negative control. Following this, hybridisation was carried out, and samples were incubated overnight at 37 °C. The following day the samples were washed, counter stained with Dapi and mounted on glass slides and stored at 4 °C.

### Bacteria single molecule inexpensive fluorescent *in situ* hybridisation (smiFISH)

smiRNAFISH within BL21-DE3 cells was adapted from previously published protocols.(Skinner et al., 2013; Tsanov et al., 2016) Briefly, BL21-DE3 cells were transformed with plasmids expressing either EGFP or PspCas13b-EGFP. A fresh colony was inoculated into 7mL of Terrific Broth containing 7 μL of Ampicillin (25mg/ml) and shaken overnight. The culture was diluted to an OD_600_ of 0.01 and IPTG was added to a final concentration of 0.05mM. Samples were incubated at 37 °C until OD_600_ of 0.5 was reached. Cells were fixed in 4% Paraformaldehyde and permeabilised overnight in 70% ethanol at -20 °C overnight. Cells were rehydrated in PBS and treated with Lysozyme (0.5mg/mL) in TEG buffer (5mM Tris (HCl), 10mM EDTA, 50mM glucose, pH 8.0)) for 5 minutes at R.T.P. Following this, hybridisation was carried out (for 16SrRNA, final probe concentration of 250nM was used), and samples were incubated overnight at 37 °C. The following day, samples were washed, counter stained with Dapi and mounted on glass slides and stored at 4 °C.

### Structure Prediction and Electrostatic Surface Potential Analysis

The structure of PspCas13b, LwCas13a and RfxCas13d was predicted using AlphaFold2(Jumper et al., 2021; Mirdita et al., 2022). The top-ranked model by pLDDT score was selected for downstream analysis. Electrostatic surface potentials were calculated using the Adaptive Poisson-Boltzmann Solver (APBS) implemented via the PyMol APBS plugin(Baker et al., 2001). Surfaces were visualised across four 90° orientations, with electrostatic potentials mapped between -5 and +5 kT/e. Positively charged surfaces were coloured blue, and negatively charged surfaces were coloured magenta.

### Phylogenetic Analysis of Cas13 Orthologs

Representative Cas13 protein sequences were retrieved from the NCBI protein database (accessed May 2026) using subtype-specific search terms based on official Type VI CRISPR protein name annotations. Sequences were retrieved for subtypes Cas13a, Cas13b, Cas13d, Cas13bt, and Cas13e using keyword searches of the NCBI protein database via the rentrez package in R, with a maximum of 200 sequences per subtype to ensure balanced representation. Cas13c was retrieved by direct accession (WP_013921683.1, WP_005959253.1) (Smargon et al., 2017). Cas13e-i representative sequences were retrieved from linked nucleotide locus records deposited by Hu et al (OL636992–OL636996) and filtered to proteins within the expected Cas13 size range (700–1,500 aa) to isolate effector sequences from flanking CRISPR locus genes (Hu et al., 2022). Curated sequences from published studies were additionally incorporated for subtypes with limited NCBI representation. An AbiF protein (WP_003231729.1) was included as an outgroup based on its identification as an evolutionary precursor to Cas13 effectors. Sequences were subsampled to a maximum of 200 per subtype to ensure balanced phylogenetic representation, yielding 687 sequences across 12 subtypes. All sequences and associated metadata are provided in the Source Data.

All sequences were analysed at the amino-acid level. Multiple sequence alignment was performed using MAFFT (v7), employing the --localpair algorithm with --maxiterate 1000 to optimise alignment quality for divergent protein sequences (Katoh & Standley, 2013). Poorly aligned and highly gapped regions were removed using trimAl with the -automated1 heuristic, which selects trimming parameters based on alignment characteristics to improve phylogenetic signal (Capella-Gutiérrez et al., 2009).

Maximum likelihood phylogenetic inference was carried out using IQ-TREE2. The best-fit amino-acid substitution model was selected automatically using ModelFinder (-m MFP) (Kalyaanamoorthy et al., 2017; Minh et al., 2020). Branch support was assessed using ultrafast bootstrap approximation (1000 replicates; -bb 1000) (Hoang et al., 2018) and SH-aLRT branch tests (1000 replicates; -alrt 1000) (Guindon et al., 2010).(Guindon et al., 2010) Computational threading was determined automatically (-nt AUTO). Nodes with ultrafast bootstrap support ≥95% and SH-aLRT support ≥80% were considered strongly supported. The resulting maximum likelihood tree was visualised in R using the ape and ggtree packages (Paradis & Schliep, 2019; Yu et al., 2017). Trees were plotted using a radial (fan) layout, with branch lengths proportional to inferred evolutionary distances. Tip labels correspond to Cas13 ortholog identifiers, and scale bars indicate amino-acid substitutions per site. Final tree visualisations were exported as high-resolution PNG and PDF files for downstream analysis and figure preparation.

#### Data Analysis

All data analysis and visualisation (graphs) were performed in GraphPad Prism software version 10, unless stated otherwise. All figures were assembled using Adobe illustrator 2024. Statistical tests and numbers of independent biological replicated are mentioned in respective figure legends. The silencing of mCherry crRNAs was analysed using unpaired two-tailed Student’s t-test or two-way Anova followed by Dunnett’s multiple comparison test where we compare every mean to a control mean as indicated in the figures. The P values (P) are indicated in the figures. P<0.05 is considered statistically significant. Colocalization of Cas13 with RNA-Binding Protein Granule markers were quantified using Pearson Correlation Coefficient. Briefly, using ImageJ/Fiji, suitable ROIs that met the criteria; sufficient intensity and dual presence of markers, were chosen manually. Colocalization analysis was performed with the Bioimaging and Optics Platform (BIOP) JACoP (Just another ColocalizationPlugin) version 9(BOLTE & CORDELIÈRES, 2006). A pipeline was established using CellProfiler to carry out spot counting to quantify number of bacteria (Stirling et al., 2021). Line intensity analysis was carried out by plotting the Z-axis profile along a region of interest. All figures were assembled using Adobe Illustrator 2025.

