## Supplementary material for "Phase-Separated RNA Condensates Govern Cas13 Target Accessibility and Cleavage": Suppl. Files

**Supplementary Information**

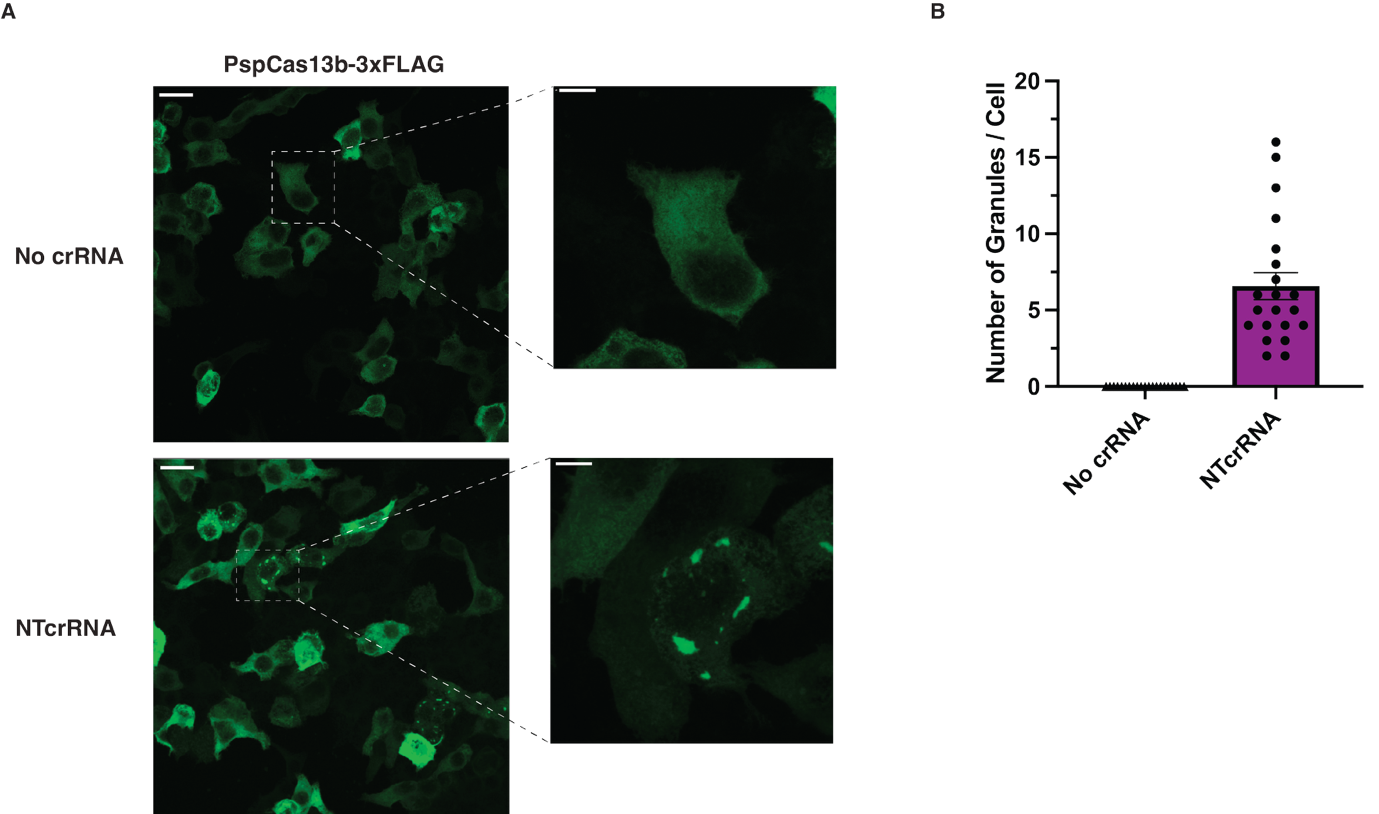

**Figure S1 | Untagged PspCas13b forms granules dependent on crRNA expression**

**(A)** Immunofluorescence of HEK293T cells expressing untagged PspCas13b-3xFLAG, either alone (No crRNA), or with its cognate non-targeting crRNA (NTcrRNA). Cells were stained with an anti-FLAG antibody. Granule structures are observed only upon crRNA co-expression. Scale bar = 20 μm. Insets show zoomed regions of representative cells depicting a single cell. Scale bar = 10 μm. **(B)** Quantification of number of granules per cell for each condition from a single replicate (n=1). Data represent individual cells.

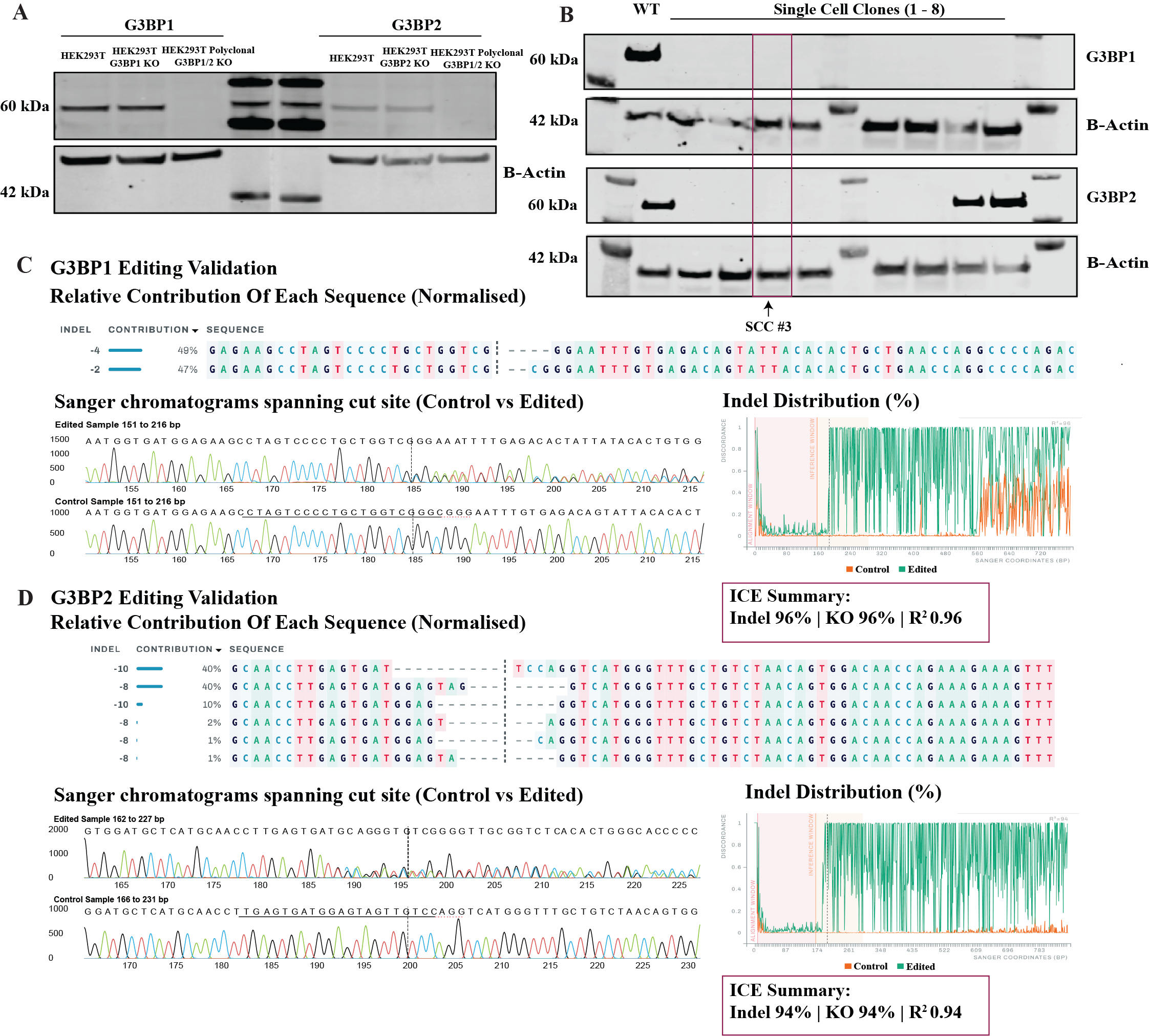

**Figure S2 | Generation and validation of G3BP1/2-knock-out HEK293T cells.**
**(A)** Representative immunoblot confirming loss of **G3BP1** and **G3BP2** protein in a CRISPR-edited **polyclonal G3BP1/2 knockout (KO)** HEK293T population relative to parental HEK293T controls. β-Actin serves as a loading control. **(B)** Immunoblot screening of individual **single-cell clones (SCC1–SCC8)** derived from the polyclonal edited population of G3BP1 and G3BP2. The clone used for downstream experiments **(SCC3)** is indicated. β-Actin serves as a loading control. **(C)** **G3BP1** editing validation by Sanger sequencing of the targeted locus followed by ICE deconvolution. Top: inferred indel composition and relative contribution of sequence variants. Bottom: Sanger chromatograms spanning the cut site for control versus edited cells and the corresponding indel distribution plot. ICE summary metrics (indel percentage, KO percentage, and R²) are shown. **(D) G3BP2** editing validation as in (C), showing inferred indel composition, control versus edited chromatograms across the cut site, indel distribution, and ICE summary metrics.

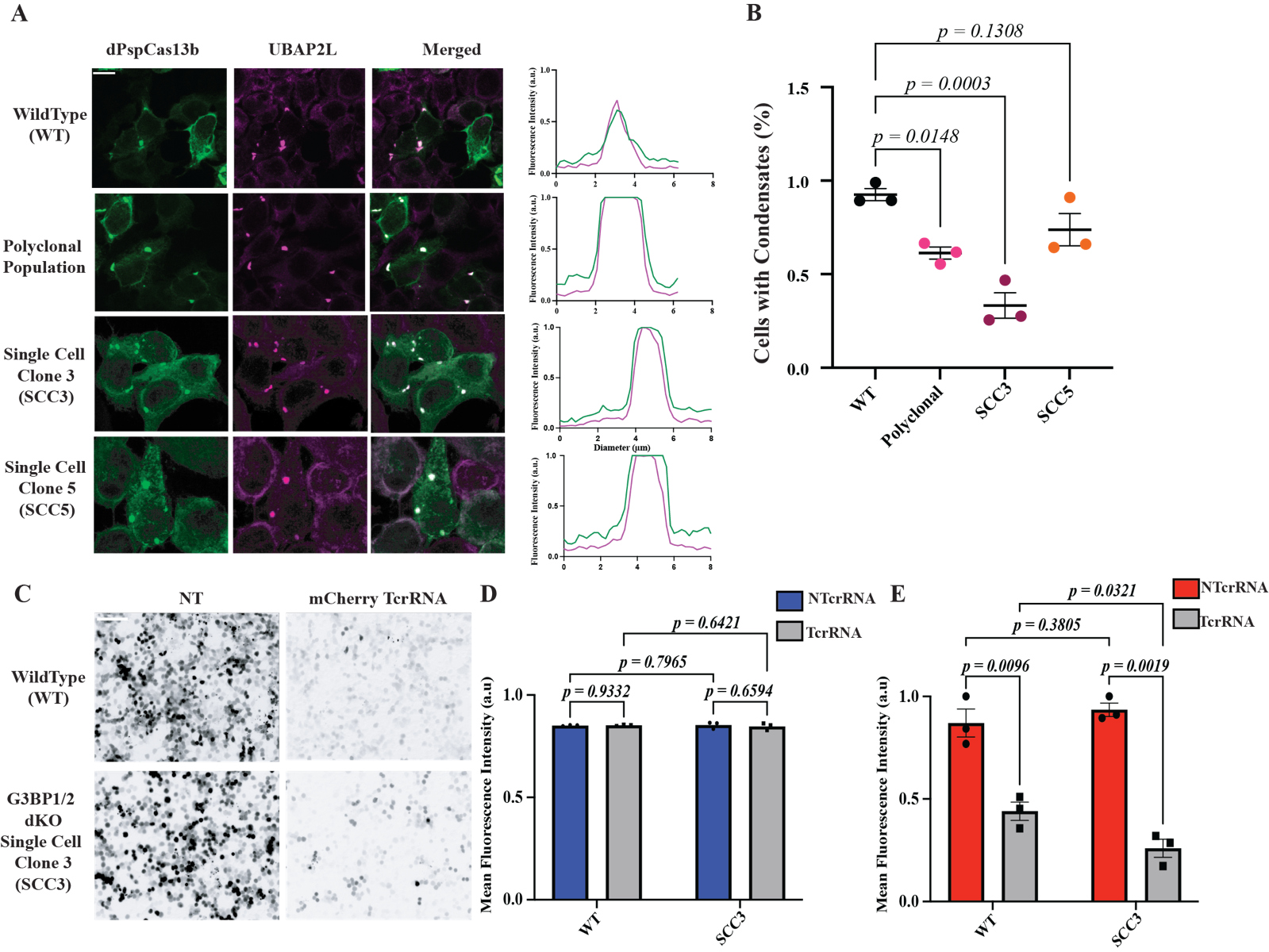

**Figure S3 | G3BP1/2 loss reduces but does not abolish UBAP2L-positive dPspCas13b condensates.
(A)** Representative confocal images of d*Psp*Cas13b (green) and endogenous UBAP2L (magenta) in wild-type (WT), a polyclonal G3BP1/2-edited population, and single-cell clones (SCC3 and SCC5), with corresponding line intensity profiles. Scale bar = 10 µm. **(B)** Quantification of the percentage of cells containing condensates across the indicated genotypes. Points represent biological replicates and p-values are shown as indicated. n = 3. Error bars are standard error of the mean (S.E.M) **(C)** Representative images of mCherry fluorescence used for image-based quantification (WT and SCC3 shown) for mCherry silencing using PspCas13b-T2A-BFP. Scale bar = 50 µm. **(D–E)** Quantification of mean fluorescence intensity measurements for BFP **(D)** and mCherry **(E).** p-values obtained using 2-way ANOVA test. Error bars are S.E.M. n=3

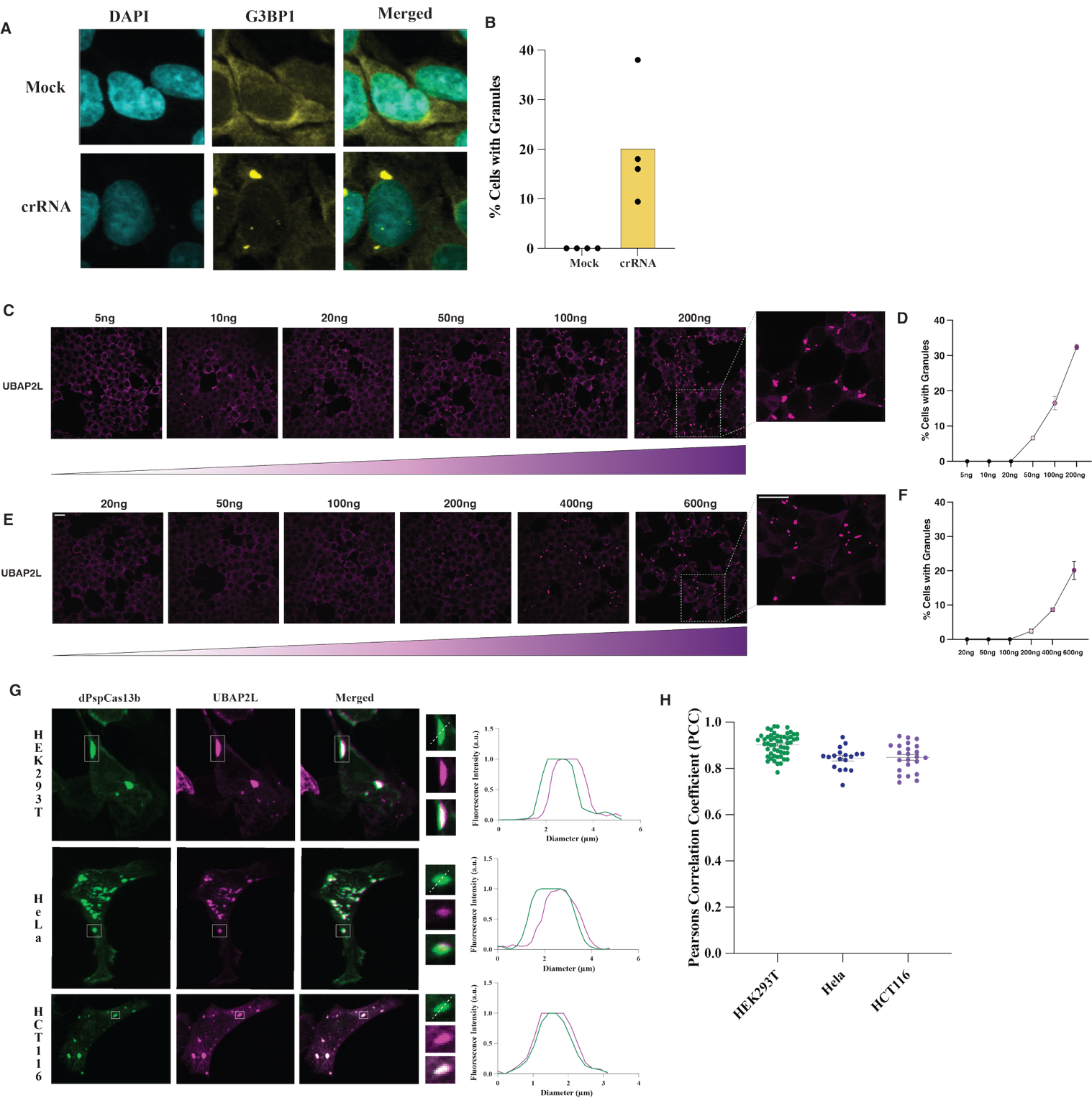

**Figure S4 | crRNA overexpression induces formation of condensates which is conserved in multiple cell lines**

**(A)** Immunofluorescence of HEK293T cells expressing the NTcrRNA alone stained with an anti-G3BP1antibody (granule marker) and DAPI. Granule formation was sufficiently observed in cells expressing the NTcrRNA, even in the absence of Cas13, suggesting crRNA overexpression alone can induce granule formation. **(B)** Quantification of the percentage of cells with visible G3BP1-positive granules in mock transfected (transfected with no plasmid load) and NTcrRNA transfected conditions. **(C, E)** Dose-dependent formation of condensates following increased amounts of NTcrRNA. 5ng – 200ng for plasmid delivery and 20ng – 600ng for IVT NTcrRNA delivery in HEK293T cells. UBAP2L was used as a positive granule marker. Insets showing a representative image of cells showing granule formation. Scale bar = 20µm. **(D, F)** Quantification of number of cells with granules from panels (C) and (E). **(G)** Representative confocal images showing colocalization of dPspCas13b with UBAP2L-mCherry in HEK293T, HeLa and HCT116 cells. White boxes indicate condensates used for line intensity profiles which depict overlapping intensity quantified across the white dotted lines. Magenta = UBAP2L-mCherry, Green = dPspCas13b-2xmNeonGreen. **(H)** Pearson’s correlation coefficients (PCC) quantifying dPspCas13b and UBAP2L colocalising across cell types. Data is shown as mean.

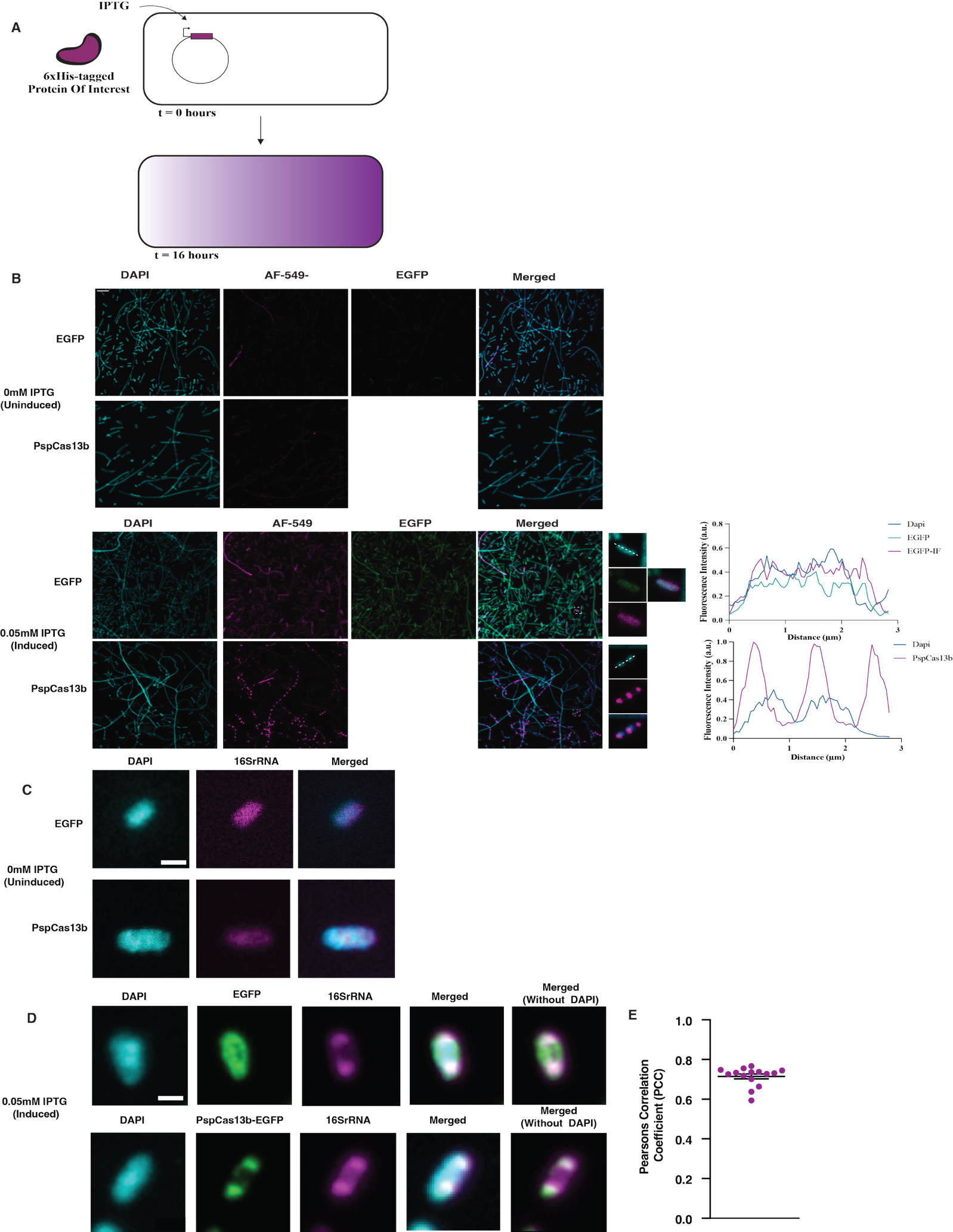

**Figure S5 | IPTG-inducible expression of PspCas13b in *E.coli* reveal distinct subcellular localisation**

**(A)** Schematic depicting IPTG-inducible expression system used to express 6xHis-tagged PspCas13 or 6xHis-tagged EGFP in *E.coli* BL21(DE3) cells. Cells were induced at 0.05mM IPTG and grown at 16 °C overnight before fixation. **(B)** Confocal images of *E.coli* cells expressing either EGFP alone or EGFP tagged PspCas13b under uninduced (0mM IPTG) or induced (0.5mM IPTG) conditions. Cells were fixed 16 hours post-inducted and stained with anti-6xHistidine antibody (shown at AF 549). Insets show zoomed-in view of representative single cells. Right panels depict line intensity profiles of fluorescence intensity across representative single cells (white dotted lines) showing colocalization of EGFP and DAPI and nucleoid exclusion of PspCas13b puncta. No crRNA was co-expressed in these experiments. **(C)** Representative single cell image of uninduced BL21-DE3 cells transformed with EGFP (top panel) and PspCas13b-EGFP (bottom panel). 16SrRNA staining is depicted using magenta. Scale bar = 1 μm. **(D)** Representative single cell images of induced BL21-DE3 cells expressing EGFP (top panel) PspCas13b-EGFP (bottom panel). Puncta localisation of 16SrRNA was observed in both EGFP and PspCas13b-EGFP expressing cells. **(E)** Pearsons’s correlation (p ~ 0.7) between PspCas13b-EGFP and 16SrRNA fluorescence. n = 15 FOVs, 3578 cells across 3 BRs.

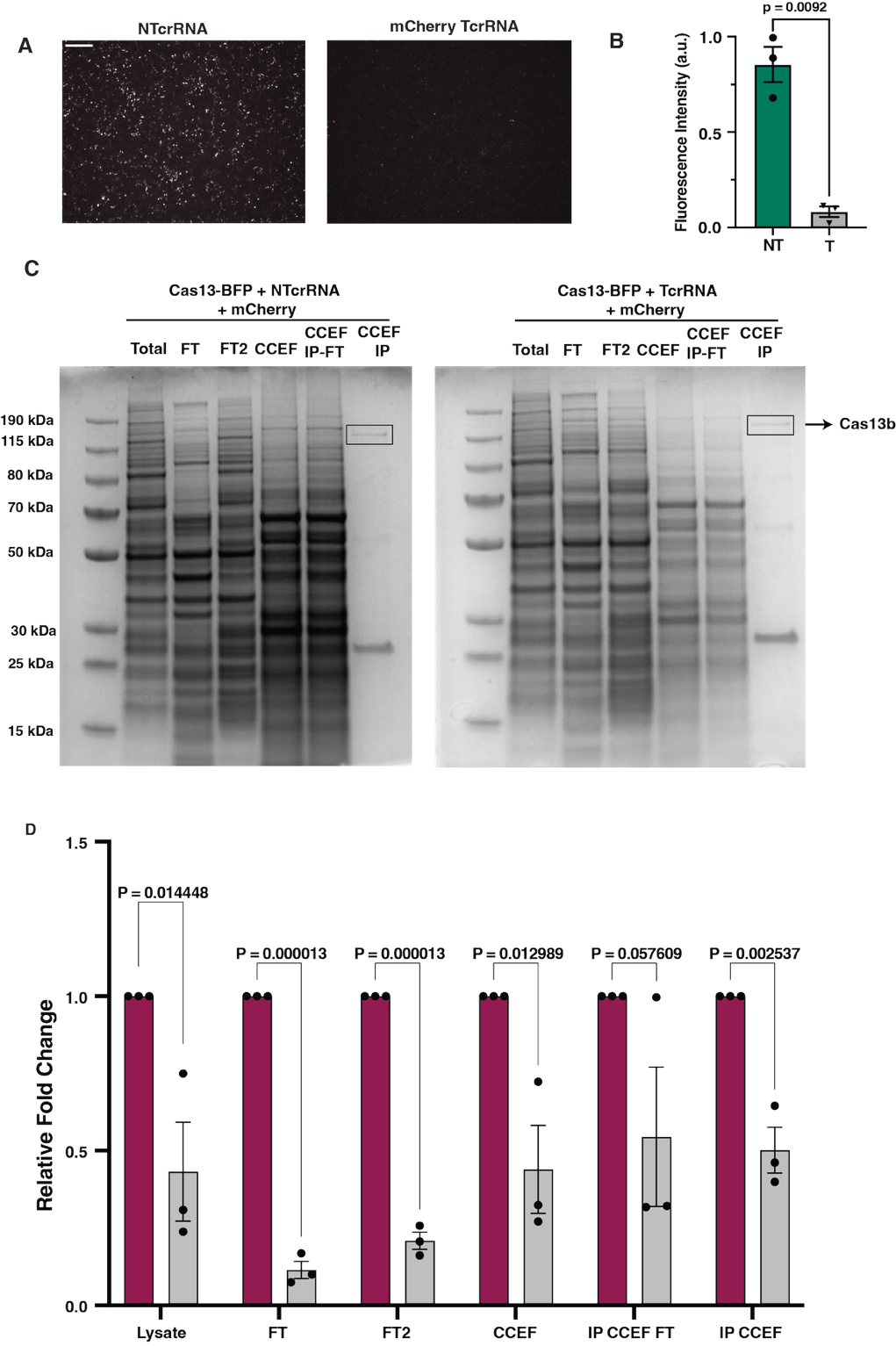

**Figure S6 | Cas13b condensates exhibit catalytic activity and mediates target mRNA cleavage (A)** Fluorescence microscopy images showing HEK293T cells expressing Cas13b-BFP and mCherry with either a non-targeting (NTcrRNA) or mCherry-targeting crRNA (TcrRNA), showing reduced mCherry fluorescence in the presence of a targeting crRNA. n = 3 biological replicates. Scale bar = 10 μm. **(B)** Quantification of mean fluorescence intensity. Error bars represent mean ± SEM. p-value calculated using unpaired two-tailed t-test. **(C)** Coomassie-stained SDS-PAGE gels showing input (Total), flow-through (FT) and immunoprecipitated fraction from cytosolic condensate-enriched fractions (CCEF) for cells expressing NTcrRNA or mCherry TcrRNA. Clean enrichment of Cas13b is evident in CCEF IP lanes. A protein band at the expected size of Cas13b is indicated. **(D)** RT-qPCR analysis of mCherry transcript levels across all fractions in non-targeting (magenta) and targeting (grey) conditions. mCherry RNA is significantly depleted across all fractions including condensate enriched fractions as well as CCEF-IP, consistent with robust RNA cleavage by Cas13b. Data shown as mean ± SEM from three biological replicates. p-values from multiple unpaired t-test.

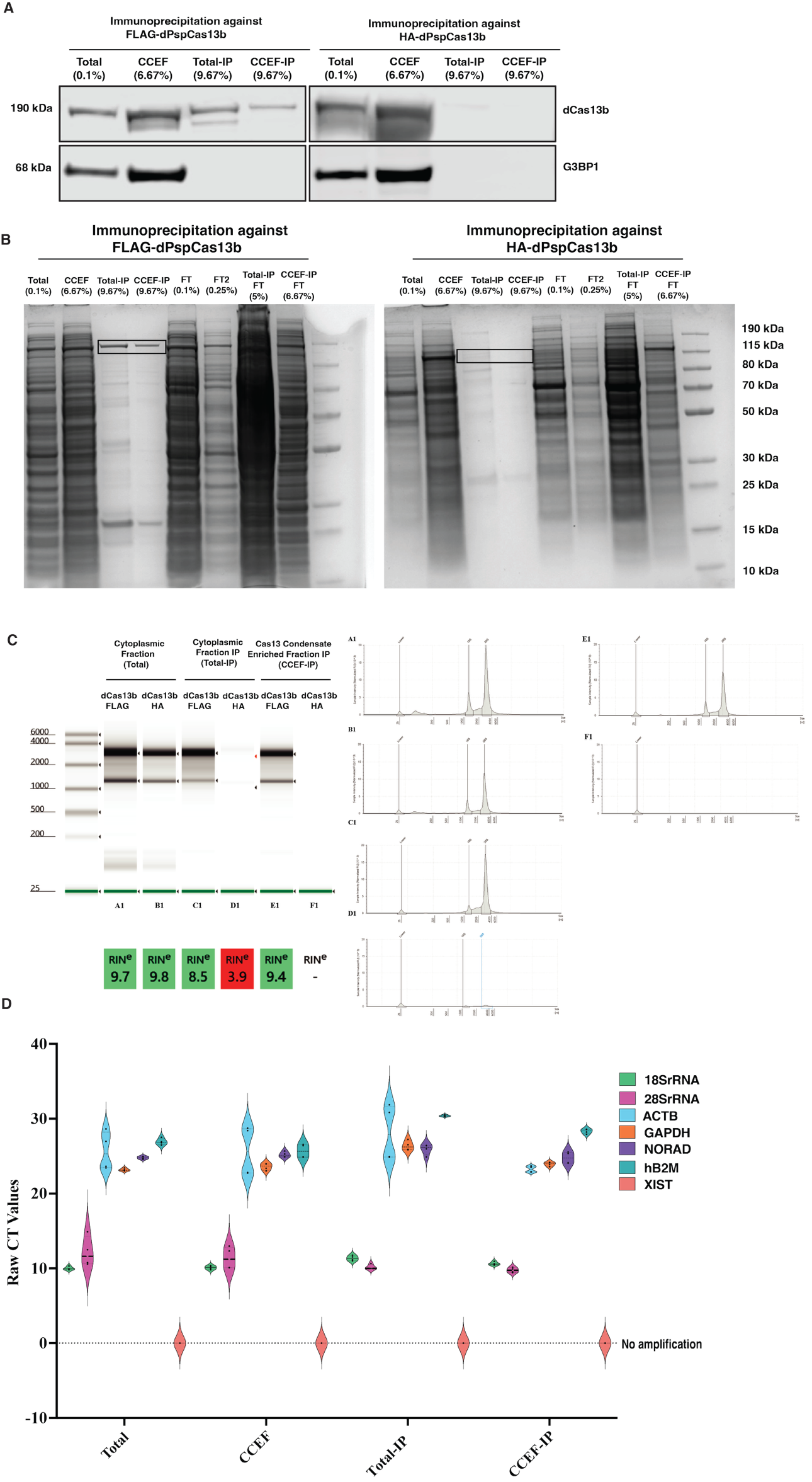

**Figure S7 | Validation of immunoprecipitation and RNA integrity of samples obtained from condensates enriched for dPspCas13b. (A)** Western blot of immunoprecipitated 3xFLAG-tagged Cas13 and G3BP1 from four different fractions, Total, CCEF, Total-IP and CCEF-IP. Immunoprecipitation was carried out using beads conjugated with a monoclonal FLAG antibody against a FLAG-tagged dPspCas13b and a HA-tagged dPspCas13b (Negative Control). Blots were probed with anti-FLAG and anti-G3BP1 antibodies to assess IP efficiency as well as background from non-specific binding to the beads. Total: Total cytoplasmic lysate input. **(B)** Coomassie-stained SDS-PAGE gels showing input (Total), flow-through (FT) , CCEF and immunoprecipitated fractions from both the input and CCEF for cells expressing dPspCas13b-3xFLAG and dPspCas13b-HA proteins. A protein band at the expected size of Cas13b is indicated with a black box. Blot on the right shows no evidence of Cas13 pulldown using HA-tagged dpspCas13b. Coomassie blots were used to further validate the specificity of the IP, however non-specific bands were seen in the Total IP and CCEF-IP lanes, for both Flag and HA-tagged dPspCas13b **(C)** RNA quality assessment of RNA from three fractions, Cytoplasmic (Total), Cytoplasmic IP (Total-IP) and Cas13-condensate enriched fraction IP (CCEF-IP). RNA was extracted and analysed using High Sensitivity RNA TapeStation on the Agilent Tapestation 4150 bioanalyzer. Representative electropherograms (A1-F1) and Rin^e^ values are shown. No presence of RNA was seen in HA-tagged dPspCas13b IP fractions. **(D)** RT-qPCR analysis visualised as a violin plot showing raw CT values. Targets are shown in the legend. Quality control of the RNA was carried out using RT-qPCR, using a panel of primers, from non-coding to coding RNAs, as well as Xist as an indicator for nuclear contamination. Mean CT values are plotted. Dashed line represents no amplification (no CT values recovered). Mean values are plotted.

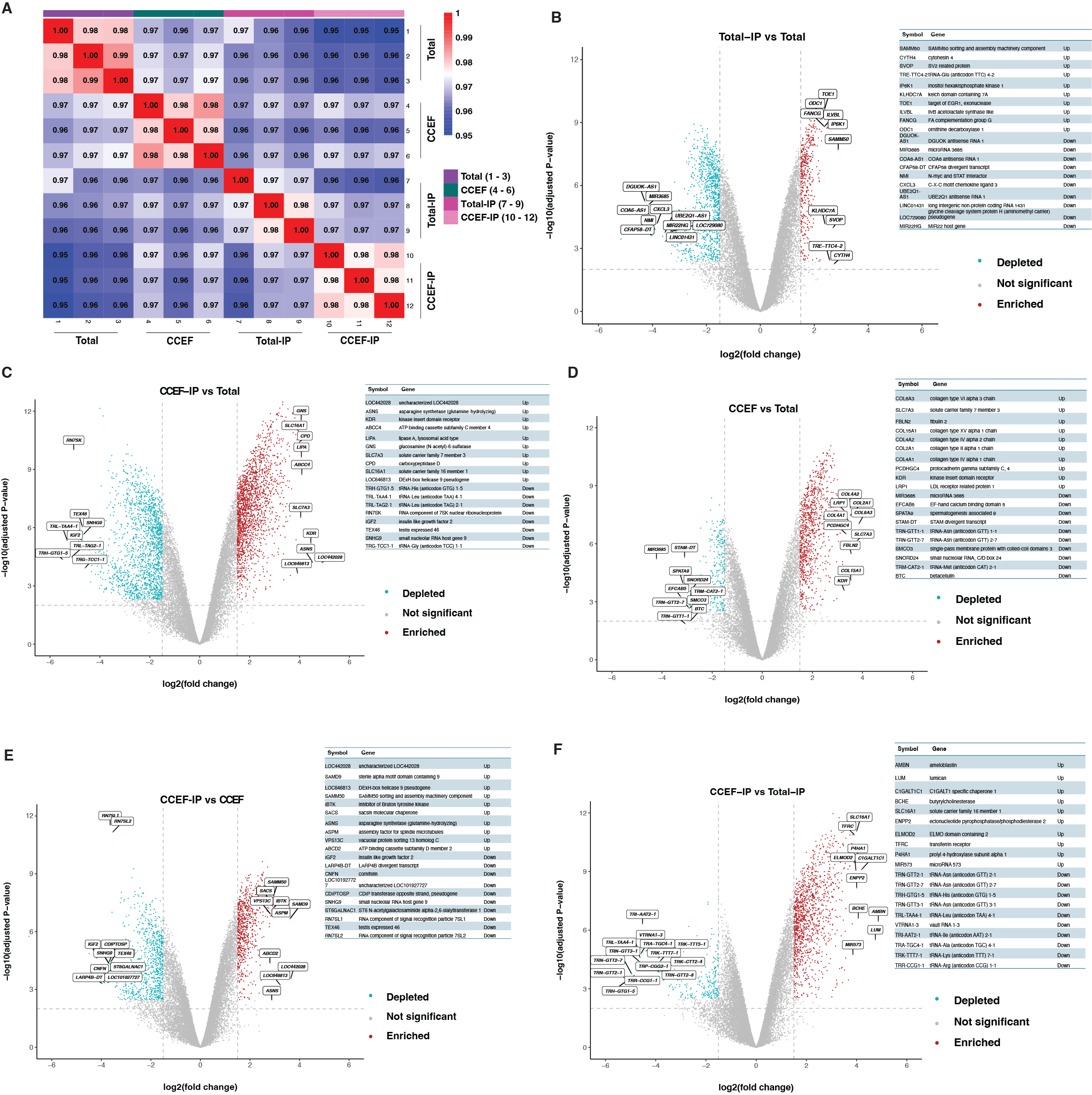

**Figure S8 | Transcriptomic profiling of RNA enrichment and depletion across different biochemical fractions analyzed by Ribo-depletion Bulk RNA-Sequencing. (A)** Spearman correlation matrix of raw RNA-seq counts across all samples. Columns correspond to biological replicates from four sample types: total cytoplasmic RNA (Total, n=3), cytoplasmic condensate-enriched fraction (CCEF, n=3), dPspCas13b immunoprecipitation from Total (Total-IP, n=3) and dPspCas13b immunoprecipitation from cytoplasmic condensate-enriched fraction (CCEF-IP, n=3). The correlation matrix shows strong reproducibility across biological replicates. **(B-F)** Volcano plots showing differential expression analysis of RNA across sample comparisons: **(B)** Total-IP vs Total, **(C)** CCEF-IP vs Total, **(D)** CCEF vs Total, **(E)** CCEF-IP vs CCEF and **(F)** CCEF-IP vs Total-IP. Log_2_ (fold change) is plotted against -log_10_(adjusted p-value) for each transcript. Genes significantly enriched (red) or depleted (cyan) in the respective comparisons are highlighted, with transcripts selected for the silencing assay labelled by gene symbol. Thresholds: adjusted p-value <0.01, log_2_FC > 1.5 and < -1.5. Tables adjacent to each volcano plot show top enriched and depleted genes highlighted on the volcano plot with accompanying gene symbol and description. The differential expression highlights distinct RNA subsets enriched in Cas13b condensates compared to input fractions.

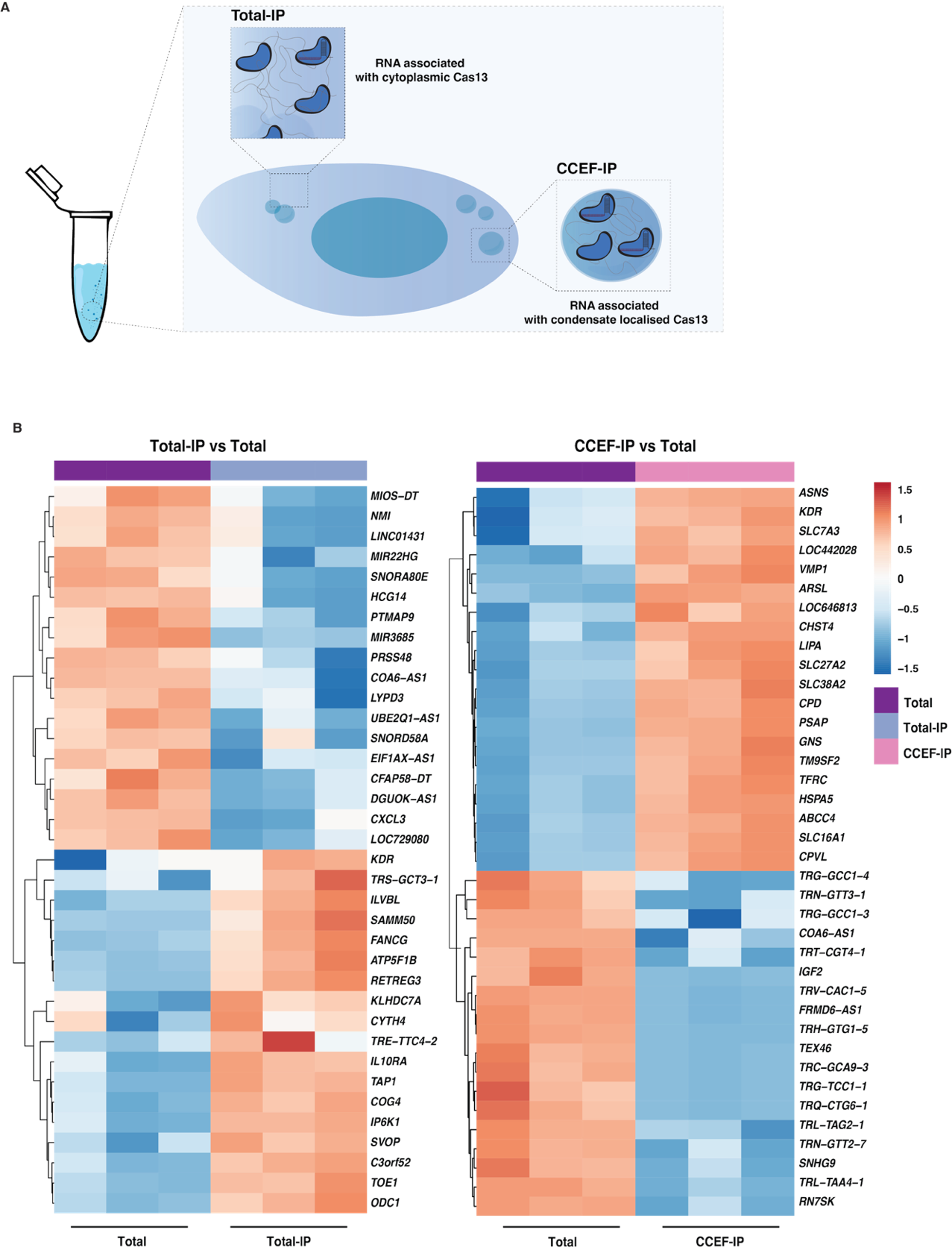

**Figure S9 | Expression profiles of top differentially enriched and depleted RNA associated with Cas13 based on its subcellular localisation. (A)** Schematic illustrating the experimental setup used: RNA populations associated with cytoplasmic Cas13 (Total-IP) and RNA populations associated with condensate-localised Cas13 (CCEF-IP) were isolated from cells.  **(B)** Heatmaps showing the expression of the top 20 enriched and 20 depleted transcripts identified from differential expression analysis of Total-IP vs Total (left) and CCEF-IP vs Total (right) as shown in Figure 5C – D. Expression values are voom-nomalised log_2_ counts per million (logCPM) and each column represents an individual biological replicate from one of four sample types: Total cytoplasmic RNA (Total), Condensate-enriched fraction (CCEF), Immunoprecipitation of dPspCas13b from Total (Total-IP) and dPspCas13b immunoprecipitation from cytoplasmic condensate-enriched fraction (CCEF-IP). Hierarchical clustering was applied to rows (genes) only using Euclidean distance and visualised using the *pheatmap* R package. These profiles suggest that Cas13b associates with distinct RNA subsets depending on its subcellular localization which may influence RNA-target accessibility.

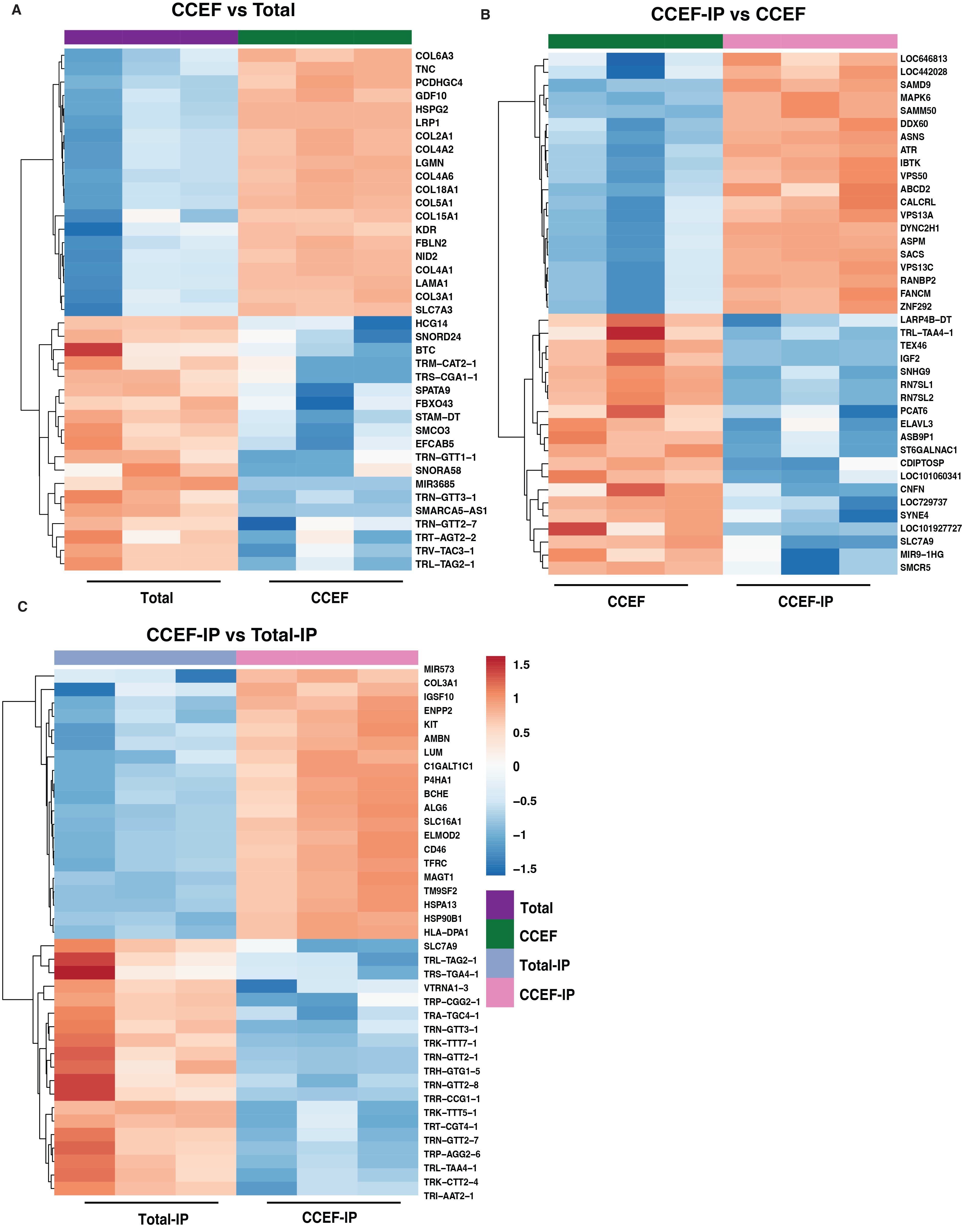

**Figure S10 | Distinct RNA expression profiles of top differentially enriched and depleted genes highlight Cas13b association in condensate vs cytoplasmic contexts. (A)** Heatmap of the top 50 enriched and top 50 depleted transcripts in **CCEF vs Total**. **(B)** Heatmap of the top differentially expressed transcripts in **CCEF-IP vs CCEF. (C)** Heatmap of the top differentially expressed transcripts in **CCEF-IP vs Total-IP.** Expression values are voom-nomalised log_2_ counts per million (logCPM) and each column represents an individual biological replicate from one of four sample types: Total cytoplasmic RNA (Total), Condensate-enriched fraction (CCEF), Immunoprecipitation of dPspCas13b from Total (Total-IP) and dPspCas13b immunoprecipitation from cytoplasmic condensate-enriched fraction (CCEF-IP). Hierarchical clustering was applied to rows (genes) only using Euclidean distance and visualised using the *pheatmap* R package. Columns (samples) are ordered manually to reflect the experimental design. These profiles suggest that Cas13b RNA association landscape is shaped by its subcellular localization, with condensate partitioning revealing a distinct set of RNAs.

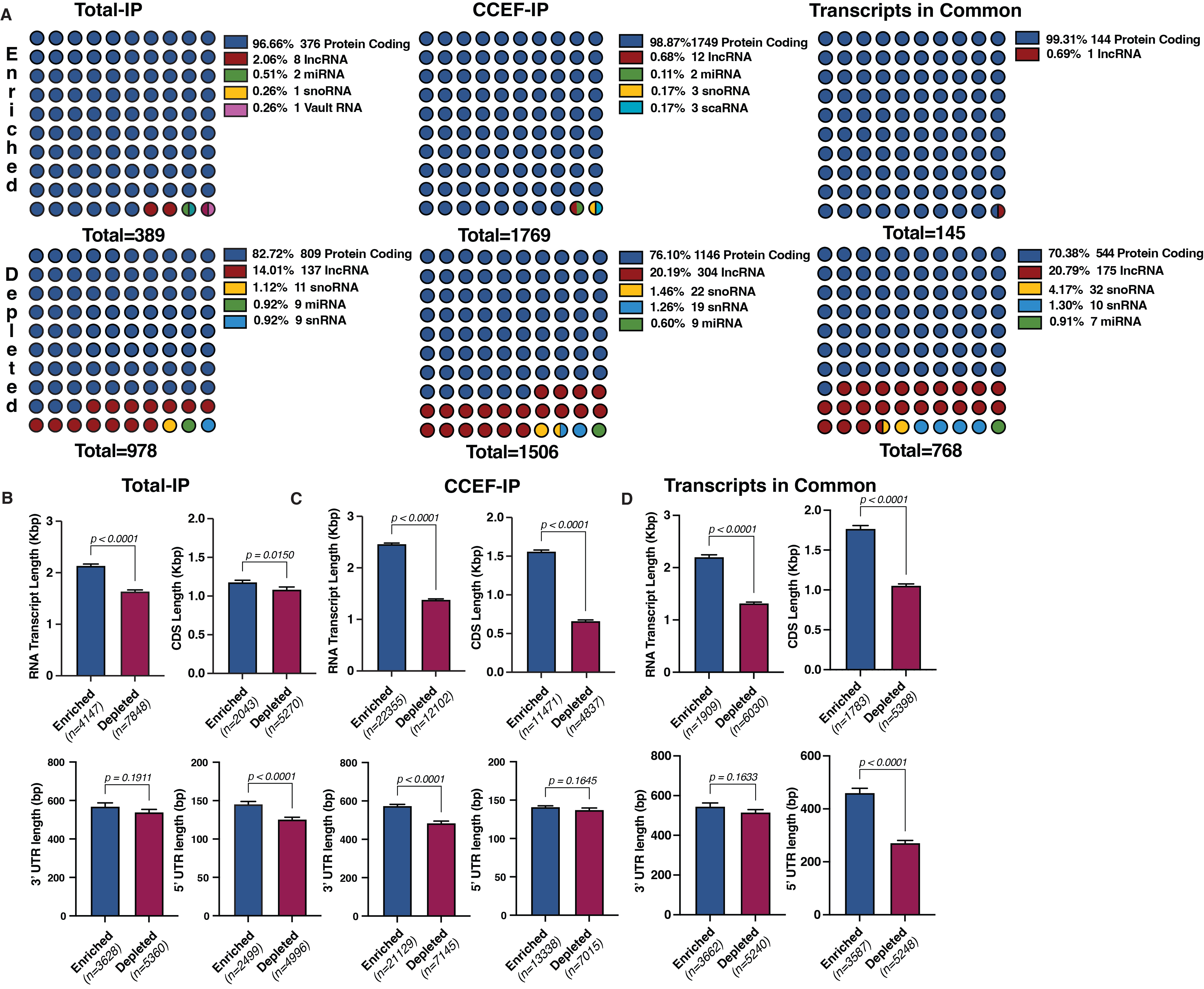

**Figure S11 | Transcript biotype and length of RNAs enriched and depleted across the different transcriptomes. (A)** Dot plot pie charts showing the biotype distribution of transcripts enriched or depleted in Total-IP (left), CCEF-IP (middle) or common to both fractions (right). The majority of enriched transcripts are protein-coding, while depleted transcripts include a broader range of non-coding RNA biotypes. Transcript biotype annotations were obtained from Ensembl via BioMart **(B–D)**. Comparisons of transcript features between enriched and depleted RNAs from: **(B)** Total-IP, **(C)** CCEF-IP, **(D)** Transcripts in Common. Bar plots display mean values for total transcript length, coding sequence (CDS) length, and 5′ and 3′ UTR lengths (in base pairs). Data are shown as mean ± SEM. Enriched RNAs tend to be longer overall and contain extended coding sequences compared to depleted RNAs, suggesting that transcript architecture may influence identity of RNAs localised to condensate enriched fractions

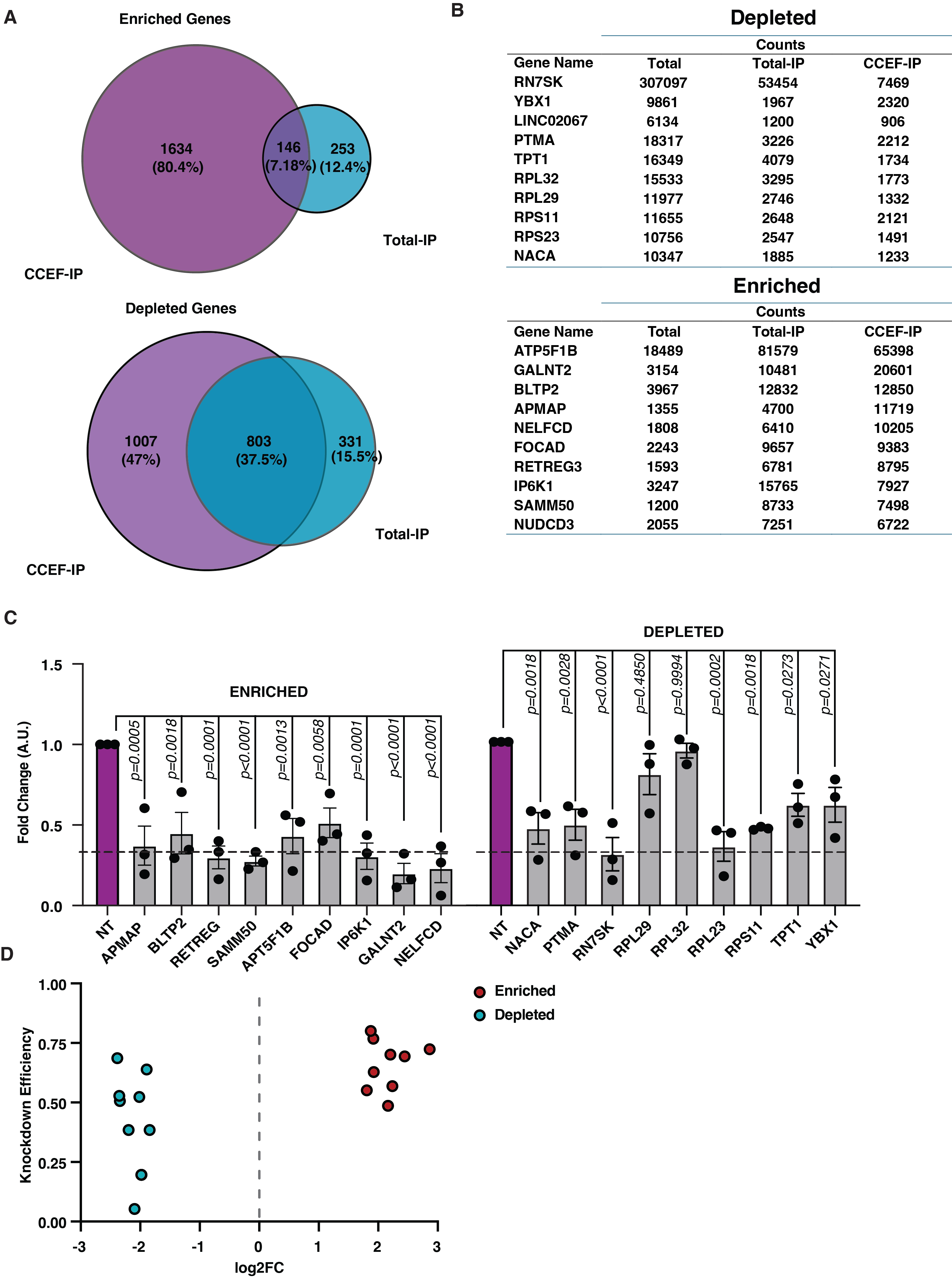

**Figure S12 | Abundance and differential association of RNA targets selected for Cas13 silencing assay. (A)** Venn diagram of genes enriched and depleted within both IP fractions. Genes were selected above threshold (fdr < 0.01, log2 fold change - </> 1.5 and high abundance. **(B)** Table of the 10 enriched and 10 depleted transcripts associated with Cas13b from both IP fractions (Total-IP and CCEF-IP) ranked by abundance. Raw RNA-seq read counts are shown for each gene across Total, Total-IP, and CCEF-IP samples. **(B)** Bar plot of log₂(fold change) values for the 10 enriched and 10 depleted RNAs identified in both IP fractions compared to total cytoplasmic input. Genes enriched in IP are shown in red, depleted genes in cyan. These transcripts correspond to targets shown in Figure 5F–G. **(C)** RT-qPCR showing average mean silencing efficacy of genes that are high associated or depleted from PspCas13b. Values were normalised to non-targeting condition. P-values are ordinary one-way Anova with multiple comparisons against NT. n=3. 2 crRNAs were used for each target. **(D)** Knockdown Efficiency plotted against log2FC (association). Log2FC of >0 represents transcripts enriched (Red) while log2FC < 0 represents transcripts depleted (Blue) from Cas13.

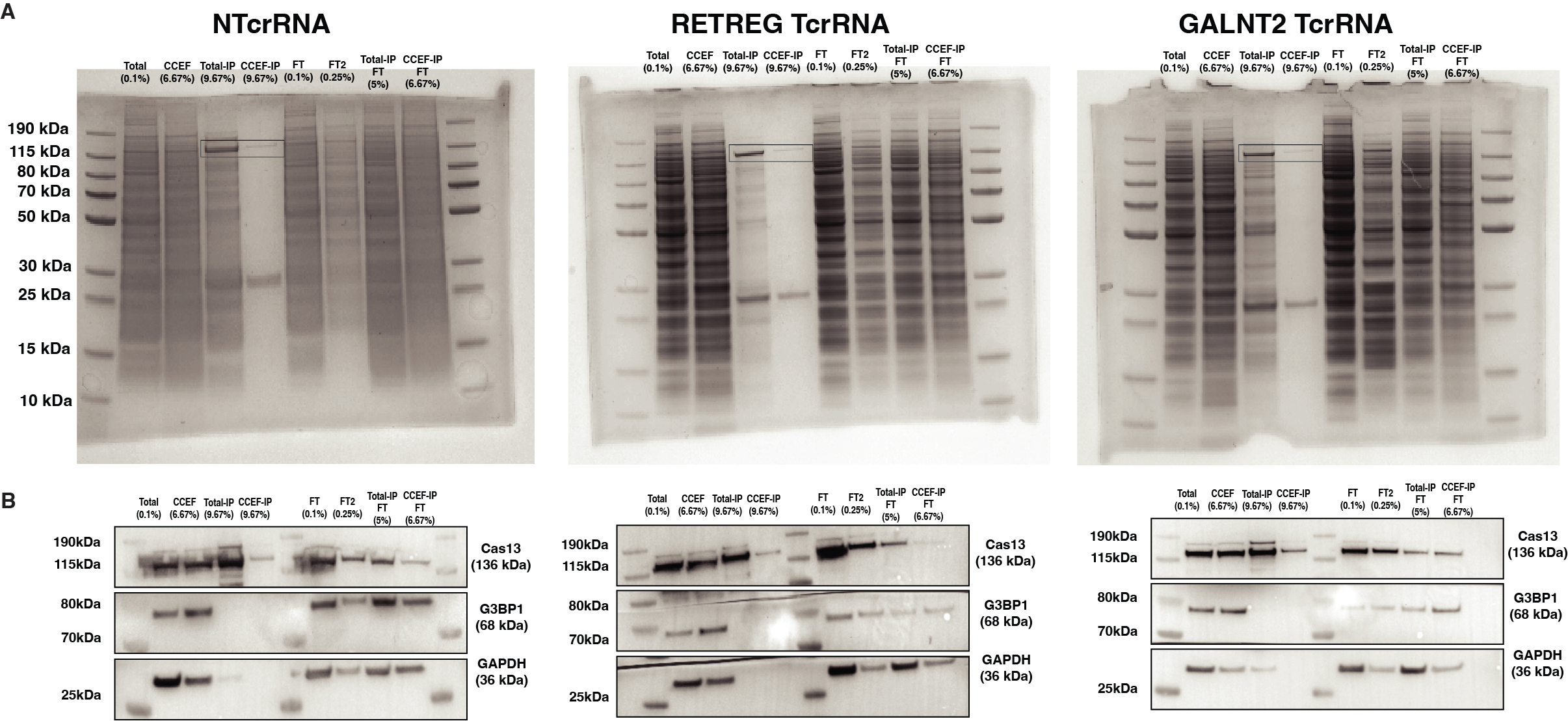

**Figure S13 | Validation of Immunoprecipitation against PspCas13b in the presence of a catalytic active Cas13 and targeting crRNAs. (A)** Coomassie-stained SDS-PAGE gels showing protein profiles from total cytoplasmic lysate, condensate-enriched fraction (CCEF), and immunoprecipitation (IP) from both total and CCEF fractions against FLAG-tagged PspCas13b. Samples were collected from cells expressing a non-targeting crRNA (NTcrRNA), or crRNAs targeting **RETREG3** or **GALNT2**. Flow-through fractions (FT, FT2) are included to assess depletion post-IP. Boxed regions highlight PspCas13b protein bands. **(B)** Western blot validation of the corresponding samples, probed for Flag-tagged **Cas13,** stress granule marker **G3BP1**, and cytoplasmic marker **GAPDH**. Cas13 is robustly enriched in both IP conditions while G3BP1 is selectively recovered in condensate-associated fractions, and GAPDH is largely absent, confirming specificity of biochemical fractionation and immunoprecipitation.

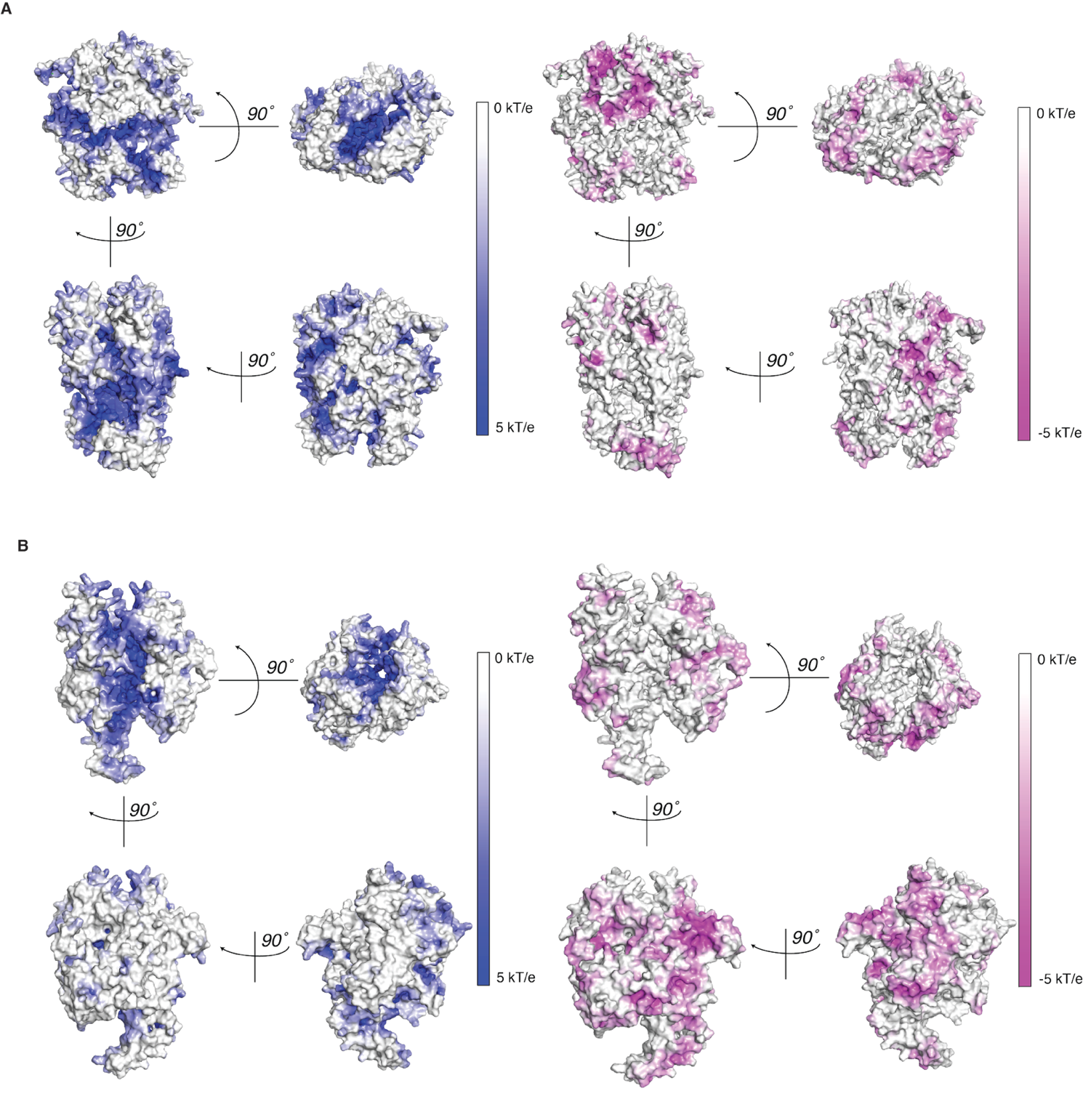

**Figure S14 | Predicted electrostatic surface potentials of Cas13 orthologues reveal asymmetric charge distributions and conserved positively charged RNA-binding surfaces**. (A) Electrostatic surface representation of LwaCas13a and (B) RfxCas13d, calculated using APBS in PyMOL. Structures are rotated in 90° increments to visualise surface charge topology. Positively charged regions are shown in blue (+5 kT/e) and negatively charged regions in magenta (−5 kT/e). Both Cas13a and Cas13d exhibit patchy and asymmetric electrostatic surface distributions, including prominent positively charged surface grooves consistent with previously described RNA-binding regions in Cas13 structures. These features resemble the electrostatic landscape observed for PspCas13b in Figure 8A, suggesting that broad RNA association may be facilitated by conserved electrostatic properties across Cas13 subtypes.

These analyses are qualitative and provide structural context for the experimentally observed RNA association patterns described in this study.

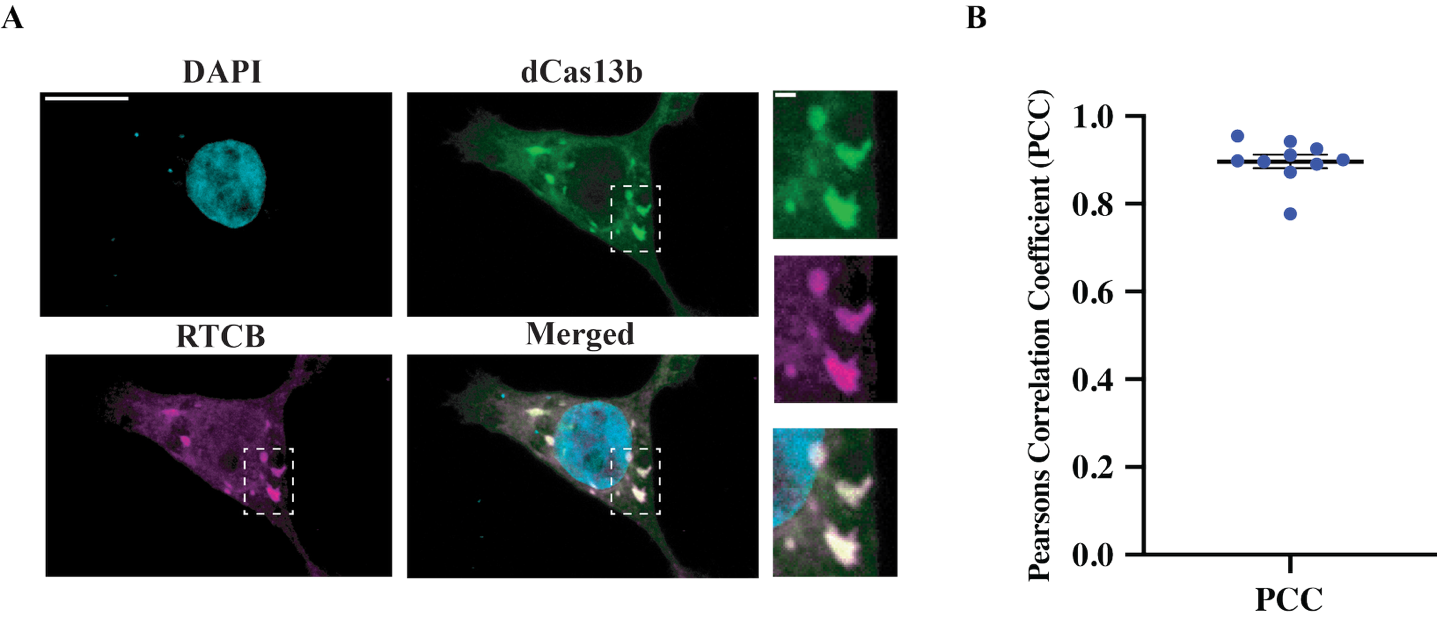

**Figure S15 | Endogenous RTCB is detectable within condensates enriched with d*Psp*Cas13b. (A)** Representative image of HEK293T cells co-transfected with d*Psp*Cas13b-2xmNeonGreen and its cognate NTcrRNA, depicting colocalization with RTCB, visualized using immunofluorescence. Scale bar = 10 µm. **(B)** Pearsons Correlation Coefficient (p ~ 0.89) between RTCB and d*Psp*Cas13b within cytoplasmic granules. n = 10 cells. 1 BR.

**Supplementary Tables**

**Table S1: crRNA Spacers used in this study**

| crRNA Name | Orthologue | crRNA sequence (Fwd 5’--- 3’) |
| --- | --- | --- |
| NTcrRNA | *PspCas13b* | *TAGATTGCTGTTCTACCAAGTAATCCATCA* |
| NTcrRNA | *LwCas13a* | TAGATTGCTGTTCTACCAAGTAATCCAT |
| mCherry_crRNA1 | PspCas13b | GGGCGCCGTCCTCGGGGTACATCCGCTCGG |
| APMAP_crRNA1 | PspCas13b | GGGTTAGGGCGGATGGTCGACATGCCCACC |
| APMAP_crRNA2 | PspCas13b | GGGTCTCTCAGATAAGAAATCCAGCATGGA |
| ATP5F1B_crRNA1 | PspCas13b | GGGTGAAGCGAAAGATGTTATCAATAAATA |
| ATP5F1B_crRNA2 | PspCas13b | GGATCCATGATACGAGAGGTGGAGTCTAGA |
| BLTP2_crRNA1 | PspCas13b | GGTGGATAAGGGTGAAGGTATCCACAGCCC |
| BLTP2_crRNA2 | PspCas13b | GGAGGAGGTTGTTCATGGTAAACCAGCCCA |
| FOCAD_crRNA1 | PspCas13b | GGAATCGTCCTTGAGAGACTAAGTACAGTT |
| FOCAD_crRNA2 | PspCas13b | GGTATAAGCTGTGCAGAATCATCCAGGCCA |
| GALNT2_crRNA1 | PspCas13b | GGGTAGACACGTGGGATGATAGGAAGGCTG |
| GALNT2_crRNA2 | PspCas13b | GGTAGACACGTGGGATGATAGGAAGGCTGA |
| IP6K1_crRNA1 | PspCas13b | GGGGGCTAGGAGAGATGGCATCCATCAAGG |
| IP6K1_crRNA2 | PspCas13b | GGAAAATACTGAATGTAAGTGGGTGCAGCT |
| NELFCD_crRNA1 | PspCas13b | GGGAGCAGTATTGGATTGACAGCAGAGAAG |
| NELFCD_crRNA2 | PspCas13b | GGAGTTCCCCGAGTGGAGTTCCCAGCGGCC |
| NUDCD3_crRNA1 | PspCas13b | GGGACAGCAGAGCCTGTTCAGGAGGCTGCT |
| NUDCD3_crRNA2 | PspCas13b | GGGAACAGGGGACAGGGGTGGTCACAGGAG |
| RETREG3_crRNA1 | PspCas13b | GGGAGGCAGAGGCTATAGACTGGCACAGGT |
| RETREG3_crRNA1 | PspCas13b | GGATCAGCAAAGGGTTAATGCAGAAGGAAA |
| SAMM50_crRNA1 | PspCas13b | GGGGACGCAGTAATTAAGTTCCAACCGAGC |
| SAMM50_crRNA2 | PspCas13b | GGGACGCAGTAATTAAGTTCCAACCGAGCG |
| LINC02067_crRNA1 | PspCas13b | GGAGAATCCATTACAGTGGTGAAGGCAGCA |
| LINC02067_crRNA2 | PspCas13b | GGAACCCCAAAACCTGATGATTGTTCCCCA |
| NACA_crRNA1 | PspCas13b | GGGGTTTAGAGGGTGATTCCAACACCAGCC |
| NACA_crRNA2 | PspCas13b | GGGTTTAGAGGGTGATTCCAACACCAGCCC |
| PTMA_crRNA1 | PspCas13b | GGGTTGGGTGGTGGAGAGCGCGTGTCATCT |
| PTMA_crRNA2 | PspCas13b | GGTTGGGTGGTGGAGAGCGCGTGTCATCTG |
| RN7SK_crRNA1 | PspCas13b | GGAGTCTTGGAAGCTTGACTACCCTACGTT |
| RN7SK_crRNA2 | PspCas13b | GGATGTGTCTGGAGTCTTGGAAGCTTGACT |
| RPL29_crRNA1 | PspCas13b | GGGGCGGGGGTGGGGGGTCGCACCAGTCCT |
| RPL29_crRNA2 | PspCas13b | GGGCGGGGGTGGGGGGTCGCACCAGTCCTT |
| RPL32_crRNA1 | PspCas13b | GGATGGTTACAGAAAAACATCCTGTAAGCA |
| RPL32_crRNA2 | PspCas13b | GGAGACAATCTGTAAGATTCCTGTCTAGAC |
| RPS11_crRNA1 | PspCas13b | GGTTGTACTTGCGGATGTAGTGCAGATAGT |
| RPS11_crRNA2 | PspCas13b | GGATGTAGTGCAGATAGTCTCGGCGGATGA |
| RPS23_crRNA1 | PspCas13b | GGAGATTATTTATGGGTCAATCTGGCAGCC |
| RPS23_crRNA2 | PspCas13b | GGGTGCAGCCAACAGGGTACAGCTGCAAGC |
| TPT1_crRNA1 | PspCas13b | GGAATCTTCAAAGGTTTACCATGAGGTAAA |
| TPT1_crRNA2 | PspCas13b | GGTTTACCATGAGGTAAACAGCTGGGGTTC |
| YBX1_crRNA1 | PspCas13b | GGATGATGGTAGAGATGGTAAGCCGGCATT |
| YBX1_crRNA2 | PspCas13b | GGGTCTCCGCATGTAGTAAGGTGGGAACCT |

**Table S2: Primers used in this study**

| Name | Sequence (5’--- 3’) |
| --- | --- |
| mCherry_qPCR_Fwd | CAG GAC GGC GAG TTC ATC TA |
| mCherry_qPCR_Rev | GTC TTG ACC TCA GCG TCG TA |
| APMAP_Fwd | TTGTGGAGAACATGCCTGGAT |
| APMAP_Rev | GCGGCACAAACTTCATCACC |
| ATP5F1B_Fwd_crRNA1 | CACCTCTAAGGTAGCGCTGG |
| ATP5F1N_Rev_crRNA1 | GATAGCCCACAGCAGAAGGG |
| ATP5F1B_Fwd_crRNA2 | GCCCATTTGGATGCTACCAC |
| ATP5F1N_Rev_crRNA2 | CAGGATCTTTTGCACCCCAC |
| BLTP2_Fwd_crRNA1 | TTGCAACTGCCGAATGTACT |
| BLTP2_Rev_crRNA1 | TCATGGCATACTGAGCTGGG |
| BLTP2_Fwd_crRNA2 | TGACAGAGGAAGATGGACAGC |
| BLTP2_Rev_crRNA2 | GGGCCGCAGTACTACCTTAT |
| FOCAD_Fwd | TCGGATCGCCCAGGTTACTA |
| FOCAD_Rev | CAGTTCCAAGAGCCACTCCA |
| GALNT2_Fwd | TTGCCCCTTCACCTTTACCC |
| GALNT2_Rev | CCGCAGTAAGTCACTTTGGC |
| IP6K1_Fwd_crRNA1 | TTGCCAAGGTTAGCACCCTC |
| IP6K1_Rev_crRNA1 | AGCCCCAAAGAATGGGAACC |
| IP6K1_Fwd_crRNA2 | TGCCTCCCAGATTCCTCGAT |
| IP6K1_Rev_crRNA2 | AAGGGAGCAGCAGCAAATCC |
| NELFCD_Fwd | CGTCTTGAAAATGGGCACCG |
| NELFCD_Rev | ATCCAACCCAGCAGAAAGGC |
| NUCD3_Fwd | CAACACCAACCACACCCAAA |
| NUCD3_Rev | GGGACAGAGATGAGGCAGTT |
| RETREG3_Fwd | GTTCTGCAGCCCCTAGTGAA |
| RETREG3_Rev | GCAAACATGTCAGCACACCC |
| SAMM50_Fwd | TAAGCTGGCTGAGTGCATCC |
| SAMM50_Rev | TCCGCAGTCAAAGAATCGCT |
| LINC02067_Fwd_crRNA1 | AGTACCACCAATGCCACTGA |
| LINC02067_Rev_crRNA1 | GCTGGCACTGTGAATTTGGA |
| LINC02067_Fwd_crRNA2 | TGACCAGGTCTGTTTCAGTCA |
| LINC02067_Rev_crRNA2 | ACTCCATTCACTGCTCATGC |
| NACA_Fwd | CACTCTGCCTCCTAAACAGCA |
| NACA_Rev | TTGACGAGGACCGACTGGAA |
| PTMA_Fwd | CTCCACTTCCCGTCTCAGAA |
| PTMA_Rev | AGGGCCTTGGAAGTTTTGGTT |
| RN7SK_Fwd | CGGTCTTCGGTCAAGGGTAT |
| RN7SK_Rev | GACTGCCACATGCAGCGCC |
| RPL29_Fwd | GGCCCCTACAAAGGCTTCAG |
| RPL29_Rev | CAAACCCATCCCCAGGATCA |
| RPL32_Fwd_crRNA1 | TGAGGGATATGGGATGTGGT |
| RPL32_Rev_crRNA1 | ATACCCAGCGCAGCATTTCT |
| RPL32_Fwd_crRNA2 | AGAAATGCTGCGCTGGGTAT |
| RPL32_Rev_crRNA2 | CACGACTCGTCTCCAATTGTT |
| RPS11_Fwd | AATGTGTCCATTCGAGGGCG |
| RPS11_Rev | GTTGAAGCGCACTGTCTTGC |
| RPS23_Fwd | GGATGTGGCCTGGTTACCT |
| RPS23_Rev | GCACTAAGATCAGGCTCAGCA |
| TPT1_Fwd | TCCAATCAAGGCTGGGACAT |
| TPT1_Rev | GCCCAAAACGTCTTGACCCT |
| YBX1_crRNA1_Fwd | AGACAAAAGCAGCCGATCCA |
| YBX1_crRNA1_Rev | TCAACGGGCAAAAAGCAAGC |
| YBX1_crRNA2_Fwd | GAACGAGGGATCGGAGAGTG |
| YBX1_crRNA2_Rev | CCATCACTTCTCCCTGCACA |
| 28SrRNA_Fwd | *CGTCGTGAGACAGGTTAGTTT* |
| 28SrRNA_Rev | *CCTCAGCCAAGCACATACA* |
| 18SrRNA_Fwd | *GCCGCTAGAGGTGAAATTCT* |
| 18SrRNA_Rev | *TCGGAACTACGACGGTATCT* |
| Norad_Fwd | *TGCTGTCGGAAGAGAGAAATG* |
| Norad_Rev | *CCTTCCATAAACGGCCAGTAA* |
| ACTB_Fwd | CATGTACGTTGCTATCCAGGC |
| ACTB_Rev | CTCCTTAATGTCACGCACGAT |
| GAPDH_Fwd | GGAGCGAGATCCCTCCAAAAT |
| GAPDH_Rev | GGCTGTTGTCATACTTCTCATGG |
| hB2M_Fwd | TTGACTTACTGAAGAATGGAGAGAG |
| hB2M_Rev | TCTCGATCCCACTTAACTATCTTG |
| XIST_Fwd | CTGCTGCAGCCATATTTCTTAC |
| XIST_Rev | TACGCCATAAAGGGTGTTGG |

**Table S3: smiFISH probes used in this study**

| Name | Transcript | Sequence (5’--- 3’) | Tail Sequence (5’--- 3’) |
| --- | --- | --- | --- |
| Probe 1 | mCherry | ATGATGGCCATGTTATCCTCCTCGCCCTTG | TTACACTCGGACCTCGTCGACATGCATT |
| Probe 2 | mCherry | CCATGTGCACCTTGAAGCGCATGAACTC | TTACACTCGGACCTCGTCGACATGCATT |
| Probe 3 | mCherry | TCGATCTCGAACTCGTGGCCGTTCAC | TTACACTCGGACCTCGTCGACATGCATT |
| Probe 4 | mCherry | CTTGGTCACCTTCAGCTTGGCGGTCT | TTACACTCGGACCTCGTCGACATGCATT |
| Probe 5 | mCherry | TGAACTGAGGGGACAGGATGTCCCAG | TTACACTCGGACCTCGTCGACATGCATT |
| Probe 6 | mCherry | GCGGGGTGCTTCACGTAGGCCTTGGA | TTACACTCGGACCTCGTCGACATGCATT |
| Probe 7 | mCherry | TCGGGGAAGGACAGCTTCAAGTAGTC | TTACACTCGGACCTCGTCGACATGCATT |
| Probe 8 | mCherry | AAGTTCATCACGCGCTCCCACTTGAA | TTACACTCGGACCTCGTCGACATGCATT |
| Probe 9 | mCherry | AGGGAGGAGTCCTGGGTCACGGTCACCA | TTACACTCGGACCTCGTCGACATGCATT |
| Probe 10 | mCherry | TTCACCTTGTAGATGAACTCGCCGTC | TTACACTCGGACCTCGTCGACATGCATT |
| Probe 11 | mCherry | TTCTTCTGCATTACGGGGCCGTCGGA | TTACACTCGGACCTCGTCGACATGCATT |
| Probe 12 | mCherry | TACATCCGCTCGGAGGAGGCCTCCCA | TTACACTCGGACCTCGTCGACATGCATT |
| Probe 13 | mCherry | ATCTCGCCCTTCAGGGCGCCGTCCTC | TTACACTCGGACCTCGTCGACATGCATT |
| Probe 14 | mCherry | CGCCGTCCTTCAGCTTCAGCCTCTGC | TTACACTCGGACCTCGTCGACATGCATT |
| Probe 15 | mCherry | AGGTGGTCTTGACCTCAGCGTCGTAG | TTACACTCGGACCTCGTCGACATGCATT |
| Probe 16 | mCherry | TAGGCGCCGGGCAGCTGCACGGGCTTCTT | TTACACTCGGACCTCGTCGACATGCATT |
| Probe 17 | mCherry | TGGGAGGTGATGTCCAACTTGATGTTGAC | TTACACTCGGACCTCGTCGACATGCATT |
| Probe 18 | mCherry | TTCGTACTGTTCCACGATGGTGTAGTCCTC | TTACACTCGGACCTCGTCGACATGCATT |
| Probe 19 | mCherry | CTTGTACAGCTCGTCCATGCCGCCGG | TTACACTCGGACCTCGTCGACATGCATT |
| Probe 1 | B-Actin | GAGCGCGGCGATATCATCATCCATGG | TTACACTCGGACCTCGTCGACATGCATT |
| Probe 2 | B-Actin | GAATCCTTCTGACCCATGCCCACCAT | TTACACTCGGACCTCGTCGACATGCATT |
| Probe 3 | B-Actin | ATGCCTCTCTTGCTCTGGGCCTCGTC | TTACACTCGGACCTCGTCGACATGCATT |
| Probe 4 | B-Actin | GATGCCGTGCTCGATGGGGTACTTCA | TTACACTCGGACCTCGTCGACATGCATT |
| Probe 5 | B-Actin | TTCTCGCGGTTGGCCTTGGGGTTCAG | TTACACTCGGACCTCGTCGACATGCATT |
| Probe 6 | B-Actin | TGTTGAAGGTCTCAAACATGATCTGGGTCA | TTACACTCGGACCTCGTCGACATGCATT |
| Probe 7 | B-Actin | CAGCCTGGATAGCAACGTACATGGCT | TTACACTCGGACCTCGTCGACATGCATT |
| Probe 8 | B-Actin | ACCCCGTCACCGGAGTCCATCACGAT | TTACACTCGGACCTCGTCGACATGCATT |
| Probe 9 | B-Actin | TCGGTGAGGATCTTCATGAGGTAGTCAG | TTACACTCGGACCTCGTCGACATGCATT |
| Probe 10 | B-Actin | CTTCTCCTTAATGTCACGCACGATTTCC | TTACACTCGGACCTCGTCGACATGCATT |
| Probe 11 | B-Actin | TGCCCAGGAAGGAAGGCTGGAAGAGT | TTACACTCGGACCTCGTCGACATGCATT |
| Probe 12 | B-Actin | CTTCATGATGGAGTTGAAGGTAGTTTC | TTACACTCGGACCTCGTCGACATGCATT |
| Probe 13 | B-Actin | GAGGAGCAATGATCTTGATCTTCATTGTGC | TTACACTCGGACCTCGTCGACATGCATT |
| Probe 14 | B-Actin | CTAAGTCATAGTCCGCCTAGAAGCAT | TTACACTCGGACCTCGTCGACATGCATT |
| Probe 15 | B-Actin | GGTTTTGTCAAGAAAGGGTGTAACGC | TTACACTCGGACCTCGTCGACATGCATT |
| Probe 16 | B-Actin | GCCATGCCAATCTCATCTTGTTTTCT | TTACACTCGGACCTCGTCGACATGCATT |
| Probe 17 | B-Actin | CCAGTTTTTAAATCCTGAGTCAAGCC | TTACACTCGGACCTCGTCGACATGCATT |
| Probe 18 | B-Actin | CAACAATGTGCAATCAAAGTCCTCGG | TTACACTCGGACCTCGTCGACATGCATT |
| Probe 19 | B-Actin | AGGATGGCAAGGGACTTCCTGTAACA | TTACACTCGGACCTCGTCGACATGCATT |
| Probe 20 | B-Actin | GCCATTCTCCTTAGAGAGAAGTGGGG | TTACACTCGGACCTCGTCGACATGCATT |
| Probe 21 | B-Actin | TACACGAAAGCAATGCTATCACCTCC | TTACACTCGGACCTCGTCGACATGCATT |
| Probe 22 | B-Actin | GAGACCAAAAGCCTTCATACATCTCAA | TTACACTCGGACCTCGTCGACATGCATT |
| Probe 23 | B-Actin | AAGTCAGTGTACAGGTAAGCCCTGGC | TTACACTCGGACCTCGTCGACATGCATT |
| Probe 24 | B-Actin | AAGGTGTGCACTTTTATTCAACTGGTC | TTACACTCGGACCTCGTCGACATGCATT |
| Probe 25 | B-Actin | GGTGCCAGATTTTCTCCATGTCGTCCCA | TTACACTCGGACCTCGTCGACATGCATT |
| Probe 26 | B-Actin | GTCAGGCAGCTCGTAGCTCTTCTCCAG | TTACACTCGGACCTCGTCGACATGCATT |
| Probe 27 | B-Actin | GCGTACAGGTCTTTGCGGATGTCCAC | TTACACTCGGACCTCGTCGACATGCATT |
| Probe 28 | B-Actin | ATCCTGTCGGCAATGCCAGGGTACAT | TTACACTCGGACCTCGTCGACATGCATT |
| Probe 29 | B-Actin | TTGCTGATCCACATCTGCTGGAAGGT | TTACACTCGGACCTCGTCGACATGCATT |
| Probe 30 | B-Actin | TGAACTTTGGGGGATGCTCGCTCCAA | TTACACTCGGACCTCGTCGACATGCATT |
| Probe 1 | 18SrRNA | TGAGACAAGCATATGCTACTGGCAGG | TTACACTCGGACCTCGTCGACATGCATT |
| Probe 2 | 18SrRNA | TGCGTACTCAGACATGCATGGCTTA | TTACACTCGGACCTCGTCGACATGCATT |
| Probe 3 | 18SrRNA | GAACCATAACTGATTTAATGAGCCATTCGC | TTACACTCGGACCTCGTCGACATGCATT |
| Probe 4 | 18SrRNA | TATCCAAGTAGGAGAGGAGCGAGCGA | TTACACTCGGACCTCGTCGACATGCATT |
| Probe 5 | 18SrRNA | CGGCATGTATTAGCTCTAGAATTACC | TTACACTCGGACCTCGTCGACATGCATT |
| Probe 6 | 18SrRNA | GGGTTGGTTTTGATCTGATAAATGCAC | TTACACTCGGACCTCGTCGACATGCATT |
| Probe 7 | 18SrRNA | GCCCGAGGTTATCTAGAGTCACCAAA | TTACACTCGGACCTCGTCGACATGCATT |
| Probe 8 | 18SrRNA | GCACGGCGACTACCATCGAAAGTTGA | TTACACTCGGACCTCGTCGACATGCATT |
| Probe 9 | 18SrRNA | GGGCCTCGAAAGAGTCCTGTATTGTT | TTACACTCGGACCTCGTCGACATGCATT |
| Probe 10 | 18SrRNA | CCCTCCAATGGATCCTCGTTAAAGGA | TTACACTCGGACCTCGTCGACATGCATT |
| Probe 11 | 18SrRNA | GCAGCAACTTTAATATACGCTATTGGAGC | TTACACTCGGACCTCGTCGACATGCATT |
| Probe 12 | 18SrRNA | CGGGACACTCAGCTAAGAGCATCGAG | TTACACTCGGACCTCGTCGACATGCATT |
| Probe 13 | 18SrRNA | CGGTCCAAGAATTTCACCTCTAGCGG | TTACACTCGGACCTCGTCGACATGCATT |
| Probe 14 | 18SrRNA | CTTGGCAAATGCTTTCGCTCTGGTCC | TTACACTCGGACCTCGTCGACATGCATT |
| Probe 15 | 18SrRNA | CCTCCGACTTTCGTTCTTGATTAATG | TTACACTCGGACCTCGTCGACATGCATT |
| Probe 16 | 18SrRNA | ACCCAAAGACTTTGGTTTCCCGGAAG | TTACACTCGGACCTCGTCGACATGCATT |
| Probe 17 | 18SrRNA | TCCCGTGTTGAGTCAAATTAAGCCGC | TTACACTCGGACCTCGTCGACATGCATT |
| Probe 18 | 18SrRNA | ACGGAATCGAGAAAGAGCTATCAATC | TTACACTCGGACCTCGTCGACATGCATT |
| Probe 19 | 18SrRNA | AATCGCTCCACCAACTAAGAACGGCC | TTACACTCGGACCTCGTCGACATGCATT |
| Probe 20 | 18SrRNA | GTAACTAGTTAGCATGCCAGAGTCTC | TTACACTCGGACCTCGTCGACATGCATT |
| Probe 21 | 18SrRNA | GAACGCCACTTGTCCCTCTAAGAAGT | TTACACTCGGACCTCGTCGACATGCATT |
| Probe 22 | 18SrRNA | CATCACAGACCTGTTATTGCTCAATC | TTACACTCGGACCTCGTCGACATGCATT |
| Probe 23 | 18SrRNA | TCCTCGTTCATGGGGAATAATTGCAA | TTACACTCGGACCTCGTCGACATGCATT |
| Probe 24 | 18SrRNA | TTCCTCTAGATAGTCAAGTTCGACCG | TTACACTCGGACCTCGTCGACATGCATT |
| Probe 1 | Poly-A | TTTTTTTTTTTTTTTTTTTTTTTTTT | TTACACTCGGACCTCGTCGACATGCATT |
| Probe 1 | 16SrRNA | TTCTGATCCACGATTACTAGCGATTCCGA | TTACACTCGGACCTCGTCGACATGCATT |
| Probe 2 | 16SrRNA | TCATGGAGTCGAGTTGCAGACTCCAATCC | TTACACTCGGACCTCGTCGACATGCATT |
| Probe 3 | 16SrRNA | AGTACGACGCACTTTATGAGGTCCGCTTG | TTACACTCGGACCTCGTCGACATGCATT |
| Probe 4 | 16SrRNA | CATTGTAGCACGTGTGTAGACCCTGGTCG | TTACACTCGGACCTCGTCGACATGCATT |
| Probe 5 | 16SrRNA | AGGGCCATGATGACTTGACGTCATCC | TTACACTCGGACCTCGTCGACATGCATT |
| Probe 6 | 16SrRNA | CAAGACCAGGTAAGGTTCTTCGCGTTGCAT | TTACACTCGGACCTCGTCGACATGCATT |
| Probe 7 | 16SrRNA | CGGTGCTTCTTCTGCGGGTAACGTCAATG | TTACACTCGGACCTCGTCGACATGCATT |
| Probe 8 | 16SrRNA | AGTTCCAGTGTGGCTGGTCATCCTCT | TTACACTCGGACCTCGTCGACATGCATT |
| Probe 9 | 16SrRNA | ATCTGGGCACATCCGATGGCAAGAGG | TTACACTCGGACCTCGTCGACATGCATT |
| Probe 10 | 16SrRNA | CCGAAGGTTAAGCTACCTACTTCTTTTGCAAC | TTACACTCGGACCTCGTCGACATGCATT |
| Probe 11 | 16SrRNA | CTCGCGAGGTCGCTTCTCTTTGTATGC | TTACACTCGGACCTCGTCGACATGCATT |
| Probe 12 | 16SrRNA | GTGGACTACCAGGGTATCTAATCCTGTTTG | TTACACTCGGACCTCGTCGACATGCATT |
| Probe 13 | 16SrRNA | TACGCATTTCACCGCTATCACCTGGAA | TTACACTCGGACCTCGTCGACATGCATT |
| Probe 14 | 16SrRNA | TCAGATGCAGTTCCCAGGTTGAGCCC | TTACACTCGGACCTCGTCGACATGCATT |
| Probe 15 | 16SrRNA | GTGCGCTTTACGCCCAGTAATTCCGATT | TTACACTCGGACCTCGTCGACATGCATT |
| Probe 16 | 16SrRNA | CAAAGGTATTAACTTTACTCCCTTCCTCCC | TTACACTCGGACCTCGTCGACATGCATT |
| Probe 17 | 16SrRNA | CATCAGGCTTGCGCCCATTGTGCAAT | TTACACTCGGACCTCGTCGACATGCATT |
| Probe 18 | 16SrRNA | GTTATGCGGTATTAGCTACCGTTTCCAGTAGT | TTACACTCGGACCTCGTCGACATGCATT |
| Probe 19 | 16SrRNA | ACCTTCCTCCAGTTTATCACTGGCAGT | TTACACTCGGACCTCGTCGACATGCATT |
| Probe 20 | 16SrRNA | GCACAACCTCCAAGTCGACATCGTTTACG | TTACACTCGGACCTCGTCGACATGCATT |
| Probe 21 | 16SrRNA | CTTCGCCACCGGTATTCCTCCAGATC | TTACACTCGGACCTCGTCGACATGCATT |
| Probe 22 | 16SrRNA | CCCTCTACGAGACTCAAGCTTGCCAG | TTACACTCGGACCTCGTCGACATGCATT |
| Probe 23 | 16SrRNA | CCTCCATCAGGCAGTTTCCCAGACAT | TTACACTCGGACCTCGTCGACATGCATT |

**Table S4: Antibodies used in this study**

| Target (Clone) | Type | Application | Host | Manufacturer | Catalogue # | Dilution |
| --- | --- | --- | --- | --- | --- | --- |
| UBAP2L (E5x4E) | Primary | WB/IF | Rabbit | Cell Signalling Technology | 40199S | 1:1000/1:200 |
| G3BP1 (E9G1M) | Primary | WB/IF | Rabbit | Cell Signalling Technology | 61559T | 1:1000/1:500 |
| GAPDH (13C10) | Primary | WB | Rabbit | Cell Signalling Technology | 2118S | 1:1000 |
| B-Actin (AC-74) | Primary | WB | Mouse | Sigma | A2228 | 1:5000 |
| RTCB | Primary | IF | Rabbit | proteintech | 19809-1-AP | 1:50 |
| 6x-Histidine (GT359) | Primary | IF | Mouse | Sigma | SAB2702218 | 1:500 |
| HA (6E2) | Primary | WB/IF | Mouse | Cell Signalling Technology | 2367 | 1:2000/1:100 |
| FLAG (M2) | Primary | WB/IF | Mouse | Sigma | F1804 | 1:2000/1:100 |
| anti-Mouse IgG IRDye 680RD | Secondary | IF/WB | Goat | LI-COR | 925-68070 | 1:2000 |
| anti-Rabbit IgG IRDye 800CW | Secondary | IF/WB | Goat | LI-COR | 925-32211 | 1:2000 |
| anti-Rabbit Alexa Fluor 647 | Secondary | IF/WB | Goat | Invitrogen | A-21246 | 1:500 |
| anti-Mouse Alexa Fluor 594 | Secondary | IF/WB | Rabbit | Invitrogen | A-11062 | 1:500 |
| anti-Rabbit Alexa Fluor 594 | Secondary | IF/WB | Goat | Invitrogen | A-11012 | 1:500 |
| anti-Mouse Alexa Fluor 488 | Secondary | IF/WB | Goat | Invitrogen | A-11017 | 1:500 |
| anti-Mouse Alexa Fluor 546 | Secondary | IF/WB | Goat | Invitrogen | A-11003 | 1:500 |

**Table S5: Constructs used in this study**

| Construct | Addgene # | Source |
| --- | --- | --- |
| BFP-Dcp1a | 153973 | A gift from Gia Voeltz |
| pLIX403_UBAP2L_mCherry | 105286 | A gift from Eugene Yeo |
| PspCas13b-NES-HIV-T2A-BFP | 173029 | Previously established in our lab |
| pC0043-PspCas13b crRNA backbone | 103854 | A gift from Feng Zhang |
| pC0040-LwaCas13a crRNA backbone | 103851 | A gift from Feng Zhang |
| pC0049-Ef1a-dPspCas13b-NES-HIV | 103865 | A gift from Feng Zhang |
| pC0068 PspCas13b (B12) His6-TwinStrep-SUMO-Bsa1 | 115219 | A gift from Feng Zhang |
| pMSCV-IRES-mCherry | 52114 | A gift from Dario Vignali |
| pHAGE-IRES-puro-NLS-dPspCas13b-2xmNeongreen-3xFlag | 132403 | A gift from Ling-Ling Chen |
| pHAGE-IRES-puro-NLS-dRfxCas13d-EGFP-3xFLAG | 132411 | A gift from Ling-Ling Chen |
| pC015-dLwCas13a-NF | 91905 | A gift from Feng Zhang |
| pC014-LwCas13a-msfGFP | 91902 | A gift from Feng Zhang |
| p23-NES-RfxCas13d-msfGFP-NES-Flag | 165076 | A gift from Ling-Ling Chen |
| His6-EGFP | - | Established in this study |
| PspCas13b-EGFP His6-TwinStrep-SUMO-Bsa1 | - | Established in this study |
| dRfxCas13d-2xmNeonGreen-3xFLAG | - | Established in this study |
| dLwCas13a-2xmNeonGreen-3xFLAG | - | Established in this study |

**Table S6: R Packages used in this study**

| Name | Reference |
| --- | --- |
| Limma | Ritchie, M.E., Phipson, B., Wu, D., Hu, Y., Law, C.W., Shi, W., and Smyth, G.K. (2015). limma powers differential expression analyses for RNA-sequencin and microarray studies. Nucleic Acids Research 43(7), e47. |
| AnnotationDbi | Pagès H, Carlson M, Falcon S, Li N (2023). _AnnotationDbi: Manipulation of SQLite-based annotations in Bioconductor_. doi:10.18129/B9.bioc.AnnotationDb. <https://doi.org/10.18129/B9.bioc.AnnotationDbi>, R package version 1.64.1, <https://bioconductor.org/packages/AnnotationDbi>. |
| Org.Hs.eg.db | Carlson M (2023). _org.Hs.eg.db: Genome wide annotation for Human_. R package version 3.18.0. |
| pheatmap | Kolde R (2025). _pheatmap: Pretty Heatmaps_. R package version 1.0.13, <https://CRAN.R-project.org/package=pheatmap>. |
| ggplot2 | H. Wickham. ggplot2: Elegant Graphics for Data Analysis. Springer-Verlag New York, 2016. |
| edgeR | Robinson MD, McCarthy DJ and Smyth GK (2010). edgeR: a Bioconductor package for differential expression analysis of digital gene expression data.Bioinformatics 26, 139-140  McCarthy DJ, Chen Y and Smyth GK (2012). Differential expression analysis of multifactor RNA-Seq experiments with respect to biological variation. Nucleic Acids Research 40, 4288-4297  Chen Y, Lun ATL, Smyth GK (2016). From reads to genes to pathways:differential expression analysis of RNA-Seq experiments using Rsubread andthe edgeR quasi-likelihood pipeline. F1000Research 5, 1438  Chen Y, Chen L, Lun ATL, Baldoni PL, Smyth GK (2024). edgeR 4.0: powerful differential analysis of sequencing data with expanded functionality and improved support for small counts and larger datasets. bioRxiv doi:10.1101/2024.01.21.576131 |
| biomaRT | Mapping identifiers for the integration of genomic datasets with the R/Bioconductor package biomaRt. Steffen Durinck, Paul T. Spellman, Ewan Birney and Wolfgang Huber, Nature Protocols 4, 1184-1191 (2009).  BioMart and Bioconductor: a powerful link between biological databases andmicroarray data analysis. Steffen Durinck, Yves Moreau, Arek Kasprzyk, Sean Davis, Bart De Moor, Alvis Brazma and Wolfgang Huber, Bioinformatics 21, 439-3440 (2005). |
